# A comprehensive atlas of somatic mutation rates and mutational signatures in normal human cells

**DOI:** 10.64898/2026.08.28.747772

**Authors:** My H. Pham, Luke M. R. Harvey, Thomas R. W. Oliver, Ellie Dunstone, Andrew R. J. Lawson, Pantelis A. Nicola, Rashesh Sanghvi, Yvette Hooks, Emily Mitchell, Georgeina L. Jarman, Yichen Wang, Federico Abascal, Hyunchul Jung, Matthew D. C. Neville, Yoshihiro Ishida, Joanna C. Fowler, Anh Phuong Le, Sarah Moody, Henry Marshall, Natalia Brzozowska, Chuling Ding, Claudia Arnedo Pac, Heather E. Machado, Laura O’Neill, Laura Humphreys, Calli Latimer, Kourosh Saeb-Parsy, Krishnaa T. A. Mahbubani, E. Joanna Baxter, Doris M. Rassl, Rocio Vicario, Frédéric Geissmann, Kenji Kabashima, Ronald L. A. W. Bleys, Luiza Moore, Rakesh Heer, Tim H. H. Coorens, Sam Behjati, Matthew Hoare, Peter J. Campbell, Philip H. Jones, Iñigo Martincorena, Raheleh Rahbari, Michael R. Stratton

**Affiliations:** Somatic Genomics Programme, Wellcome Sanger Institute, Cambridge, UK; Great Ormond Street Hospital, London, UK; University College London Cancer Institute, London, UK; Wellcome-MRC Cambridge Stem Cell Institute, Cambridge, UK; Department of Haematology, University of Cambridge, Cambridge, UK; Early Cancer Institute, University of Cambridge, Cambridge, UK; Medical Research Council Toxicology Unit, University of Cambridge, Cambridge, UK; Department of Surgery, University of Cambridge, Cambridge, UK; Cambridge Biorepository for Translational Medicine, NIHR Cambridge Biomedical Research Centre, University of Cambridge, Cambridge, UK; Cambridge Blood and Stem Cell, Cambridge, UK; Royal Papworth Hospital NHS Foundation Trust, Cambridge, UK; Immunology Program, Sloan Kettering Institute, Memorial Sloan Kettering Cancer Center, New York, New York, United State; Department of Dermatology, Kyoto University, Kyoto, Japan; Department of Anatomy, Division of Surgical Specialties, Utrecht University, University Medical Center Utrecht, The Netherlands; Clinical Diagnostics, AstraZeneca; Charing Cross Hospital, Imperial College Healthcare NHS Trust, London, UK; European Bioinformatics Institute, European Molecular Biology Laboratory (EMBL-EBI), Cambridge, UK; Cellular Genomics Programme, Wellcome Sanger Institute, Cambridge, UK; Department of Paediatrics, University of Cambridge, Cambridge, UK

## Abstract

Over the course of a lifetime, somatic mutations accrue in normal human cells, causing variation in cell phenotype and engendering somatic evolution with outcomes ranging from the adaptive immune system to cancer. To inform understanding of somatic evolution in the human body we report the mutation rates and mutational signatures of 53 normal cell types. Most show evidence of linear mutation accumulation over time with single base substitution mutation rates ranging from ∼3.5/year/diploid genome in spermatogonia and sperm, to ∼20/year in postmitotic neurons, ∼50/year in mitotically active colorectal epithelial cells, ∼60/year in kidney proximal tubule cells and hepatocytes, 100s/year in sun-exposed skin epidermal cells and 10-50/year in the remainder. Certain cell types, including skin epidermis, cardiac myocytes, bladder urothelium, kidney proximal tubule cells, and hepatocytes, show substantial variability in mutation burdens around the linear age trend, indicating the influence of additional factors which differ between individuals and modulate mutation accumulation, including exogenous mutagen exposures. At least 18 single-base substitution and nine small insertion and deletion mutational signatures are present, some in all cell types, some in a subset and others in a single cell type. Known exogenous mutagen exposures and endogenous mutational processes account for some mutational signatures, but the origins and mechanisms underlying many are uncertain. This comprehensive survey of mutagenesis provides a foundation for understanding somatic evolution of human cell populations in health and disease.

## INTRODUCTION

Understanding of somatic evolution in human tissues is informed by knowledge of the somatic mutation rates and mutational signatures of their constituent cell types. Until recently, however, somatic mutations in normal cell genomes have been problematic to detect accurately because the error rates per base of standard DNA sequencing technologies are substantially higher than the somatic mutation rates per base of normal human cells. This problem was obviated in sequencing of cancer genomes by overlaying each interrogated genome position with multiple sequence reads^1,2^. Although each read originates from a different cancer cell, cancers are cell clones in which every cell shares a set of mutations and, thus, genuine somatic mutations are recurrently reported in these reads whereas sequencing errors are not. Normal tissues, however, are composed of myriad small cell clones, each with its own set of somatic mutations, limiting the effectiveness of this approach.

Multiple strategies have been deployed to overcome this problem. These include sequencing clones derived from single normal stem cells *in vitro*^3^, sequencing microscopically visible structures in normal tissues which are recently generated cell clones^4–6^, and sequencing small tissue areas which usually include only one or a small number of such clones^5^. For cell types in which the tissue architecture or cell biology lend themselves to such approaches, early studies have revealed substantial differences in somatic mutation rates and contributory mutational signatures^4,6–11^. However, for many normal cell types these approaches are not effective and, thus, a comprehensive survey of mutation landscapes has not been feasible.

DNA sequencing methods have recently been developed which enable somatic mutation detection in all tissues. Notably, single-molecule duplex sequencing allows accurate detection of mutations in polyclonal tissues by separately sequencing both strands of individual DNA molecules and validating that a somatic mutation is real by its presence on both strands^12^. It has been deployed recently to determine the effects of mutagen exposures, inherited DNA repair and replication defects and landscapes of clonal expansions in non-cancer tissues ^13–21^. In this study, we utilise Restriction-Enzyme NanoSeq (RE-NanoSeq)^12^, a single-molecule duplex sequencing protocol with an error rate lower than five errors per billion base pairs, to explore comprehensively normal human tissues, generating a survey of mutation rates and mutational signatures in normal human cells^13,16,18^.

## RESULTS

### Study design

810 tissue samples from 172 individuals aged 0-93 years were studied (Supplementary Tables 1). Because duplex sequencing detects somatic mutations in a randomly selected subpopulation of cells from each sample, providing an average of the mutation rates and mutational signatures of the constituent cell types, tissue samples were flow sorted or laser-capture microdissected to enrich for specific cell types (for example, neurons and subtypes of lymphocytes) or tissue structures (for example, glomeruli and layers of the adrenal cortex) (Supplementary Note 1). DNAs from 53 cell types or tissue structures were obtained from which RE-NanoSeq libraries were constructed and sequenced (Figure 1a)^12^. A sample from each individual was subject to standard whole-genome DNA sequencing at ∼20-fold read coverage to catalogue germline inherited variation and enable identification of somatic mutations. For simplicity of nomenclature, each laser-capture microdissected tissue structure or flow-sorted cell type is referred to as a “cell type” even though they all remain, to varying extents, mixtures of cell types (Extended Figure 1).

**Figure 1.**
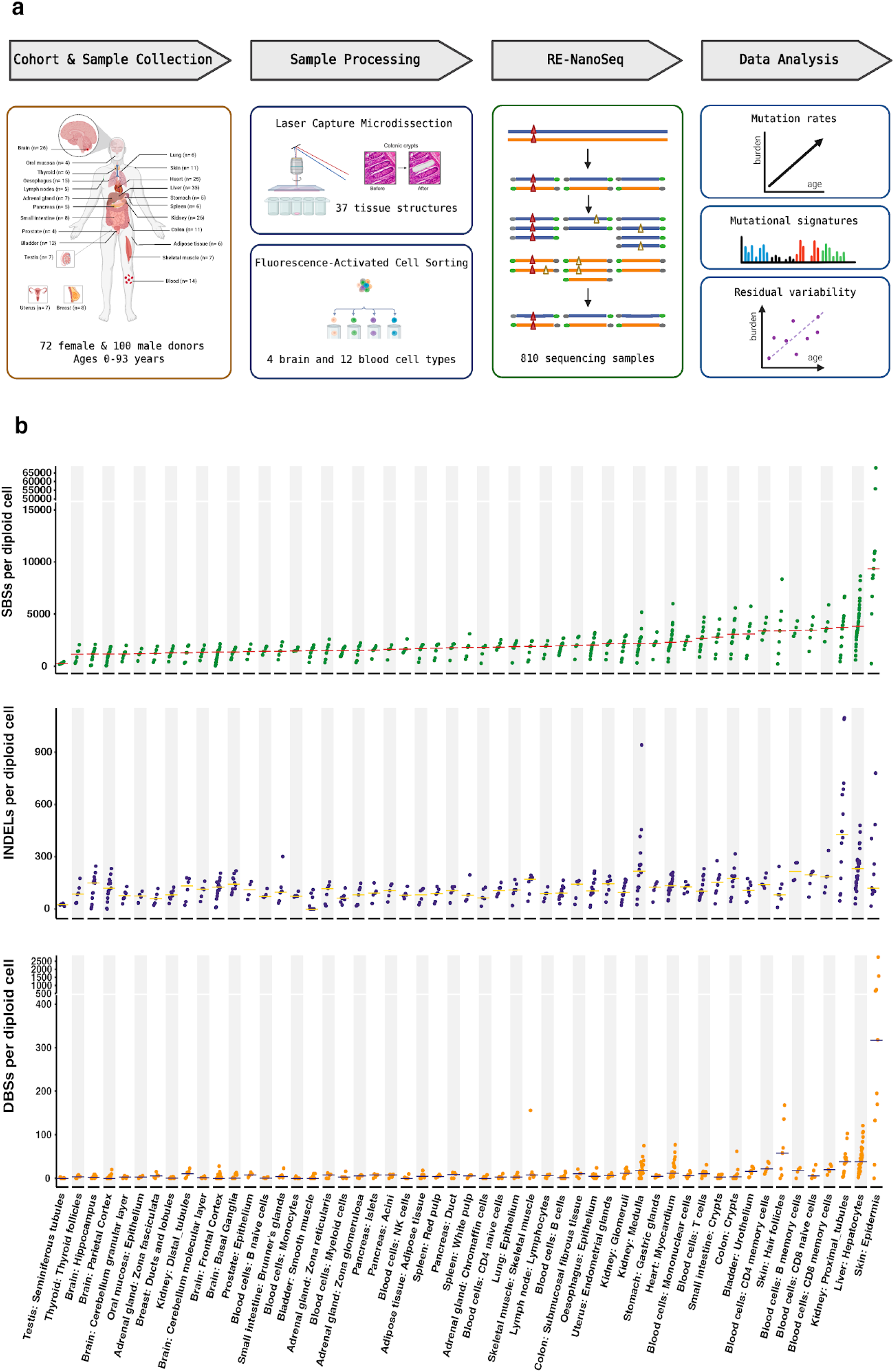
Study design and somatic mutation burden across 53 normal human cell types. (a) Overview of the experimental workflow. Solid tissues and blood/brain samples from 172 donors were collected, tissue structures and cell populations enriched by laser-capture microdissection or fluorescence-activated cell sorting, sequenced using RE-NanoSeq, and analysed for mutation rates, mutational signatures and residual variability (Methods). (b) Single-base substitution (SBS, top), small insertion/deletion (INDEL, middle) and double-base substitution (DBS, bottom) mutation burdens per diploid genome for each of the 53 cell types/tissue structures, ordered by increasing median SBS burden. Each point represents one sample; horizontal bars indicate the median burden per cell type. Note the compressed and expanded y-axis scales used to accommodate the wide dynamic range of mutation burdens across cell types.

### Somatic mutation rates

806,721 somatic single-base substitution (SBS), 10,155 double-base substitution (DBS) and 44,679 small insertion and deletion (ID) mutations were identified (Methods). RE-NanoSeq sequence coverage was used to estimate the mutation burden per diploid genome for each sample and samples from multiple individuals of different ages were analysed to characterise patterns of mutation accumulation over the human lifespan for each cell type (Figure 1b, Methods)^12^.

To investigate age-dependent mutation accumulation we compared linear, quadratic and exponential models (Supplementary Note 2). Linear accumulation of somatic SBS mutations during postnatal life provides the best fit to the observed data overall, with 39/53 cell types individually showing statistically significant evidence for linear accumulation (Figure 2, Supplementary Table 3).

**Figure 2.**
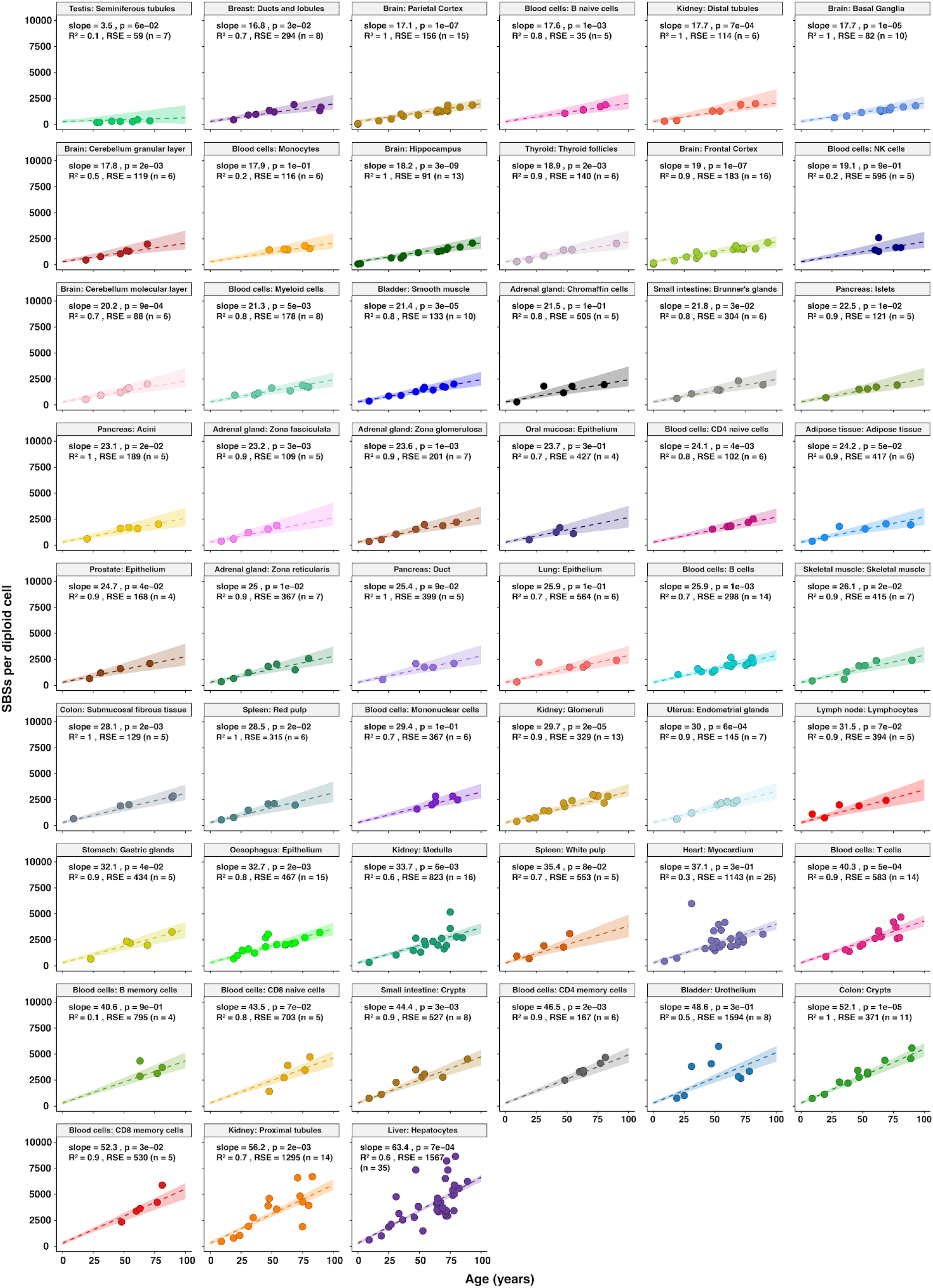
SBS mutation rates across 53 normal human cell types estimated using linear mixed-effect model. Number of SBSs per diploid cell plotted against donor age (years) for each cell type, ordered by increasing regression slope (top left to bottom right). Each point represents one sample, coloured by cell type. Dashed line, ordinary least-squares linear regression fit; shaded ribbon, 95% confidence interval estimated by bootstrapping (1,000 replicates; Methods). Slope (SBS/diploid genome/year), two-sided Student’s t-test p-value (Benjamini–Hochberg FDR-adjusted for multiple testing across cell types), coefficient of determination (R²) and residual standard error (RSE) are shown for each cell type.

Mutation rates differ, however, >20-fold between cell types. Spermatogonia and sperm from seminiferous tubules of the testis, which contribute to the male germline, exhibit substantially lower mutation rates than all other sequenced cell types (3.5 SBS/diploid genome/year). Postnatally non-dividing (postmitotic) cell types, including neurons from forebrain (frontal cortex, parietal cortex, hippocampus, basal ganglia) and hindbrain (cerebellar cortex granular and molecular cell layer) regions, show SBS mutation rates of ∼20 SBS/year demonstrating, as previously reported, that mutagenesis does not necessarily depend on genome-wide DNA replication during mitotic cell division^12,22,23^. Skeletal muscle and bladder smooth muscle cells, which are thought to be postmitotic or to have very low cell division rates, also exhibit mutation rates of ∼20 SBS/year^12^. Indeed, the mutation rates of postmitotic cell types are similar to those of many cell types which continue to proliferate throughout life, including kidney distal tubules, breast ducts and lobules, thyroid follicles, and various types of blood cells, raising the possibility that, in these too, most mutations arise between cell divisions. Nevertheless, cell types with high cell division rates, such as small intestinal and colorectal epithelial cells, show higher SBS mutation rates (40-50 SBS/year) than postmitotic cells, suggesting that genome-wide DNA replication at cell division entails a high risk of generating somatic mutations and that mutations acquired during cell division make significant contributions to mutation burdens in some tissues. However, the mutation rates of frequently dividing cell types are lower than those of infrequently dividing kidney proximal tubule cells and hepatocytes (∼60 SBS/year) indicating that other determinants of somatic mutation rate have been operative in these. Skin epidermis showed the highest mutation rate of several 100s/year. Samples of brain hippocampus, frontal cortex and parietal cortex from three individuals obtained during the first year of postnatal life show mutation burdens of 52-156 SBS, suggesting that mutation rates during the nine months of embryonic and foetal development are generally higher than those postnatally, consistent with previous observations on umbilical cord blood (55 SBS at birth)^24,25^. Cell types in which mutation rates have been measured previously using other approaches show similar results to those obtained here using RE-NanoSeq (Supplementary Table 3).

The ranking of cell types according to ID mutation rates generally mirrors that of SBS mutation rates (Extended Figure 2). Notably, spermatogonia and sperm in seminiferous tubules show very low ID mutation rates compared to all other sequenced cell types. However, kidney proximal tubule cells show disproportionately high ID mutation rates relative to SBS rates and have the highest ID mutation rate of any cell type (8.5 ID/year). The numbers of double base substitutions (DBS) identified are too small to support reliable estimation of mutation rates. Nevertheless, skin epidermis, skin hair follicles, hepatocytes, kidney proximal tubule cells and heart myocytes show high DBS mutation burdens compared to other cell types (Figure 1b, Supplementary Note 7).

### Overview of mutational signatures

Somatic mutations are caused by multiple mutational processes of exogenous and endogenous origins, including mutagen exposures and errors in DNA repair or replication, which are associated with characteristic mutational signatures. To quantify the activities of the different mutational processes operative in each cell type, SBS and ID mutational signatures were extracted from the normal tissue catalogues of somatic mutations, decomposed into COSMIC reference (v3.4, <u>cancer.sanger.ac.uk/signatures</u>) mutational signatures, and their mutation burdens in each sample estimated (Methods). Attributions of individual mutational signatures to mutation burdens across individuals and cell types and their correlations with age and exogenous exposures were then investigated (Figure 3).

**Figure 3.**
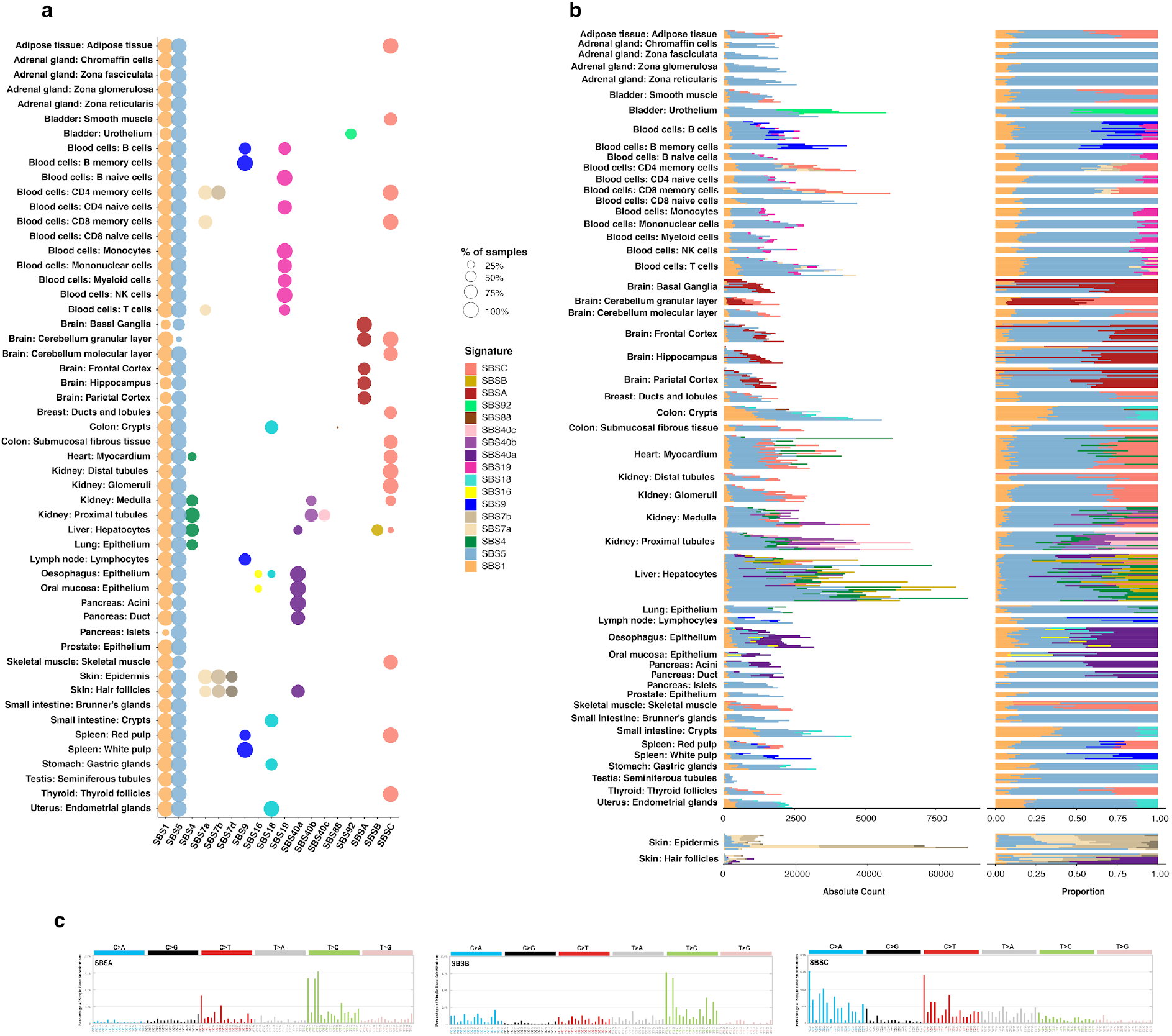
Landscape of single-base substitution mutational signatures across normal cell types. (a) Bubble plot showing the presence of each of the 18 SBS mutational signatures (columns) across the 53 cell types/tissue structures (rows). Bubble size indicates the percentage of samples of a given cell type in which the signature was attributed (Methods). Colour denotes signature identity. (b) Absolute mutation counts (left) and proportional contribution (right) of each SBS signature to the total SBS burden of each cell type, summed/averaged across all samples of that cell type and coloured as in a. (c) Mutational spectra of the three novel signatures, SBSA, SBSB and SBSC, which could not be decomposed into current COSMIC reference signatures (v3.4). Bars show the percentage of single base substitutions in each of 96 trinucleotide contexts, grouped into the six substitution classes (C>A, C>G, C>T, T>A, T>C, T>G).

Fifteen COSMIC SBS mutational signatures were identified including SBS1, SBS4, SBS5, SBS7a, SBS7b, SBS7d, SBS9, SBS16, SBS18, SBS19, SBS40a, SBS40b, SBS40c, SBS88, and SBS92, together with three additional signatures designated SBSA, SBSB, and SBSC (Supplementary Note 3). Some signatures are thought to be due to endogenous mutational processes (SBS1, SBS5, SBS9, SBS18), others to exogenous exposures (SBS4, SBS7a, SBS7b, SBS7d, SBS16, SBS88, SBS92), and others are of uncertain origin^26^ (Figure 3a). SBS1 and SBS5 are ubiquitous, present in all cell types and all individuals, with the remainder of signatures restricted to subsets of cell types and/or individuals. SBS1, SBS5, SBS18, SBS40a, SBS40c, SBSA and SBSC mutation burdens show linear correlations with age (Supplementary Note 4). The complexity of the repertoire of mutational signatures differs between cell types: some include only SBS1 and SBS5, whereas hepatocytes and kidney proximal tubule cells show five and six, respectively. Several positive correlations are observed between the mutation burdens contributed by different SBS mutational signatures likely reflecting shared components of underlying mutational processes, common cell type distributions, similar influences of biological features such as cell division rates, and other factors (Extended Figure 4).

Nine COSMIC ID mutational signatures were identified (ID1, ID2, ID3, ID5, ID8, ID9, ID11, ID13, ID21), with ID9, which is of unknown origin, almost ubiquitous across samples and cell types (Extended Figure 3). ID signatures are due both to endogenous mutational processes (ID1 and ID2, caused by polymerase slippage during DNA replication at polynucleotide repeats) and exogenous exposures (ID3 and ID13, caused respectively by tobacco smoke chemicals and ultraviolet light), with the remainder uncertain. Multiple positive correlations between the mutation burdens of SBS and ID mutational signatures likely reflect their origins from shared underlying mutational processes (examples including SBS1 and ID1, ID2, ID9; SBS5 and ID9; SBS4 and ID3, ID5; SBS7a, 7b, 7d and ID13; SBS40b and ID5; SBSA and ID21) (Extended Figure 4).

### Individual mutational signatures across cell types

SBS1 is caused by deamination of 5-methylcytosine. It shows linear accumulation with age in 21/53 cell types (Extended Figure 5). There is substantial variation in SBS1 mutation rates between cell types, with low rates in spermatogonia and postmitotic neurons (<1/year) and high rates in cell types with high mitotic rates, including small intestinal (∼17 SBS/year (95% CI: 15.8-17.7)) and colorectal epithelia (∼18 SBS/year (95% CI: 17.5-19)). Therefore, genome-wide DNA replication at mitosis appears not to be necessary for SBS1 mutagenesis but may substantially increase its rate.

SBS5 is of unknown aetiology and makes the largest contribution of any single mutational signature to the total somatic SBS burden of the human body. It shows linear accumulation with age in 40/53 cell types (Figure 3; Extended Figure 5). The SBS5 mutation rate is exceptionally low in spermatogonia (3.4 SBS/year (95% CI: 2.2-6.5), exceptionally high in hepatocytes (37.6 SBS/year (95% CI: 33.9-39.9) and at intermediate levels (∼10-30 SBS/year) in the remaining cell types. Compared to SBS1, SBS5 mutation rates appear to be influenced only to a limited extent by mitotic cell division rates. The cause of the high SBS5 mutation rate in hepatocytes is unknown, and the possibility that it is due to the activity of a different mutational process with a similar mutational signature cannot be excluded.

SBS18 is due to DNA damage caused by reactive oxygen species and is present in oesophageal, small intestinal, colorectal and endometrial epithelia, all of which are characterised by high mitotic rates (Figure 3). SBS9 is exclusively found in B lymphocytes and is due to the activity of the error-prone DNA polymerase eta on regions of the genome affected by off-target somatic hypermutation in non-naive B lymphocytes^27^ (Figure 3). SBS19 is restricted to haematopoietic and lymphoid cells (Figure 3) and has been attributed to long-lived DNA adducts the origins of which are unknown^28^.

Two SBS mutational signatures caused by exposure to tobacco smoke chemicals are found here in normal cells: SBS4 in bronchial epithelial cells, hepatocytes, cardiac myocytes, kidney proximal tubule and medulla cells, and SBS92 in bladder urothelium (Figure 3). Many individuals with a history of tobacco smoking have high SBS4 and SBS92 mutation burdens which account for the increased total mutation burdens in these cell types in exposed individuals and which indicate that tobacco smoke chemicals are circulated systemically to cause mutations in tissues beyond those directly exposed. However, the origins of small SBS4 mutation loads in some cell types are uncertain and may be due to other mutational processes or to ambiguities in signature attribution. The reasons for the sensitivities of cardiac myocytes and hepatocytes to SBS4 mutagenesis are unknown but suggest that intrinsic biological features of these cell types influence levels of mutagenesis in conjunction with ambient concentrations of the causative mutagens^13^. The cardiac muscle samples showing SBS4 are from the left side of the heart, which receives blood directly from the lung before circulation to the rest of the body, and further studies to explore whether there are differences in SBS4 mutation burden between the left and right sides of the heart would be of interest. Furthermore, the presence of SBS4 in relatively infrequently dividing cardiac myocytes and hepatocytes raises the possibility that the underlying mutational process does not depend primarily on DNA replication at mitosis. SBS4 and SBS92 exhibit very different mutation profiles. SBS4 has been observed in cells exposed *in vitro* to tobacco smoke chemicals. However, SBS92 has not, and the nature of its underlying mutational process, the reasons for its distinct mutational signature and its presence mainly in bladder urothelium are unknown^6^. It may plausibly be due to high concentrations of a subset of tobacco smoke chemicals in urine or to intrinsic features of mutagen metabolism in bladder epithelial cells.

SBS7a, SBS7b and SBS7d are caused by ultraviolet light exposure and account for the high mutation burdens of skin epidermis and hair follicles. SBS7a and SBS7b are also present in T lymphocytes^27,29^ and are likely caused by ultraviolet light exposure of circulating T cells resident for long periods in skin. SBS16 is associated with exposure to alcohol and is found in buccal and oesophageal squamous epithelia of a subset of individuals^17^. SBS88 is due to colibactin exposure, a mutagen produced by certain microbes in the colorectal microbiome, and was found in the colorectal epithelium of one individual^30,31^.

The mutational processes underlying SBS40a, SBS40b, and SBS40c are unknown. SBS40a is present in oesophageal squamous, oral squamous, hepatocellular, pancreatic acinar, and pancreatic ductal cells, and shows a linear correlation between mutation burden and age (Supplementary Note 4). SBS40b is only found in kidney proximal tubule and kidney medulla cells and SBS40c only in kidney proximal tubule cells.

SBSA, SBSB and SBSC cannot be decomposed into the current set of COSMIC reference signatures and their causes are unknown (Figure 3c). SBSA is characterised predominantly by T>C substitutions at A<u>T</u>N trinucleotides and is present in neurons of the frontal cortex, parietal cortex, hippocampus, basal ganglia, and granular cell layer of the cerebellum, but absent from the cerebellar molecular layer, a region predominantly composed of dendritic trees and glial cells. Previous reports of a similar signature were also in neurons^32–34^. SBSB is also characterised predominantly by T>C mutations at A<u>T</u>N trinucleotides together with T>A mutations at C<u>T</u>G trinucleotides. It is restricted here to hepatocytes, and previous reports of a similar signature have also been in hepatocytes and hepatocellular carcinomas^35,36^. A detailed comparison of the trinucleotide and pentanucleotide mutational profiles of SBSA, SBSB, SBS5, SBS16 and SBS92, which are all characterised by T>C mutations at A<u>T</u>N trinucleotides, highlights the differences between these mutational processes (Extended Figure 6). SBSC is characterised predominantly by C>A mutations and is found in skeletal muscle, adipose tissue, thyroid, heart and cerebellar granular cells, in which it shows linear mutation accumulation with age.

### Residual variability in somatic mutation burdens

Although most cell types exhibit linear accumulation of somatic mutations with age, the extent to which individual sample mutation burdens vary between individuals after accounting for age differs markedly between cell types (Figure 4a). To quantify this residual variability, we calculated the Residual Standard Error (RSE) for each cell type, which reflects the average deviation of individual mutation burdens from the fitted linear age model (Methods). The residual variability varies by more than an order of magnitude, with RSE values ranging from 59 (seminiferous tubules) to 1594 (bladder urothelium). Many cell types show individual SBS mutation burdens closely following the linear accumulation model (Figure 2), with accordingly low RSE values, including cerebral and cerebellar neurons (RSE 82-183), pancreatic islets (121), colonic submucosal fibrous tissue (129), bladder smooth muscle (133), endometrial epithelium (145), prostatic epithelium (168), pancreatic acini (189), adrenal gland zona glomerulosa (201), and breast epithelium (294). By contrast, high interindividual variability and scatter away from the linear model are apparent in cardiac myocytes (1143), kidney proximal tubules (1295), hepatocytes (1567) and bladder urothelium (1594) and are similarly observed for ID mutations in these cell types (Extended Figure 2). Lesser degrees of interindividual variability are observed in several subtypes of T and B lymphocytes, although the relatively small numbers of individuals studied may, in part, account for this (Supplementary Note 5). These findings indicate that, although age is the principal determinant of somatic mutation burden in most cell types, additional factors contribute substantially to certain cell types. Such factors potentially include inherited genome variation, exogenous mutagen exposures, exogenous influences on the rates of endogenous mutational processes, and supervening disease processes.

**Figure 4.**
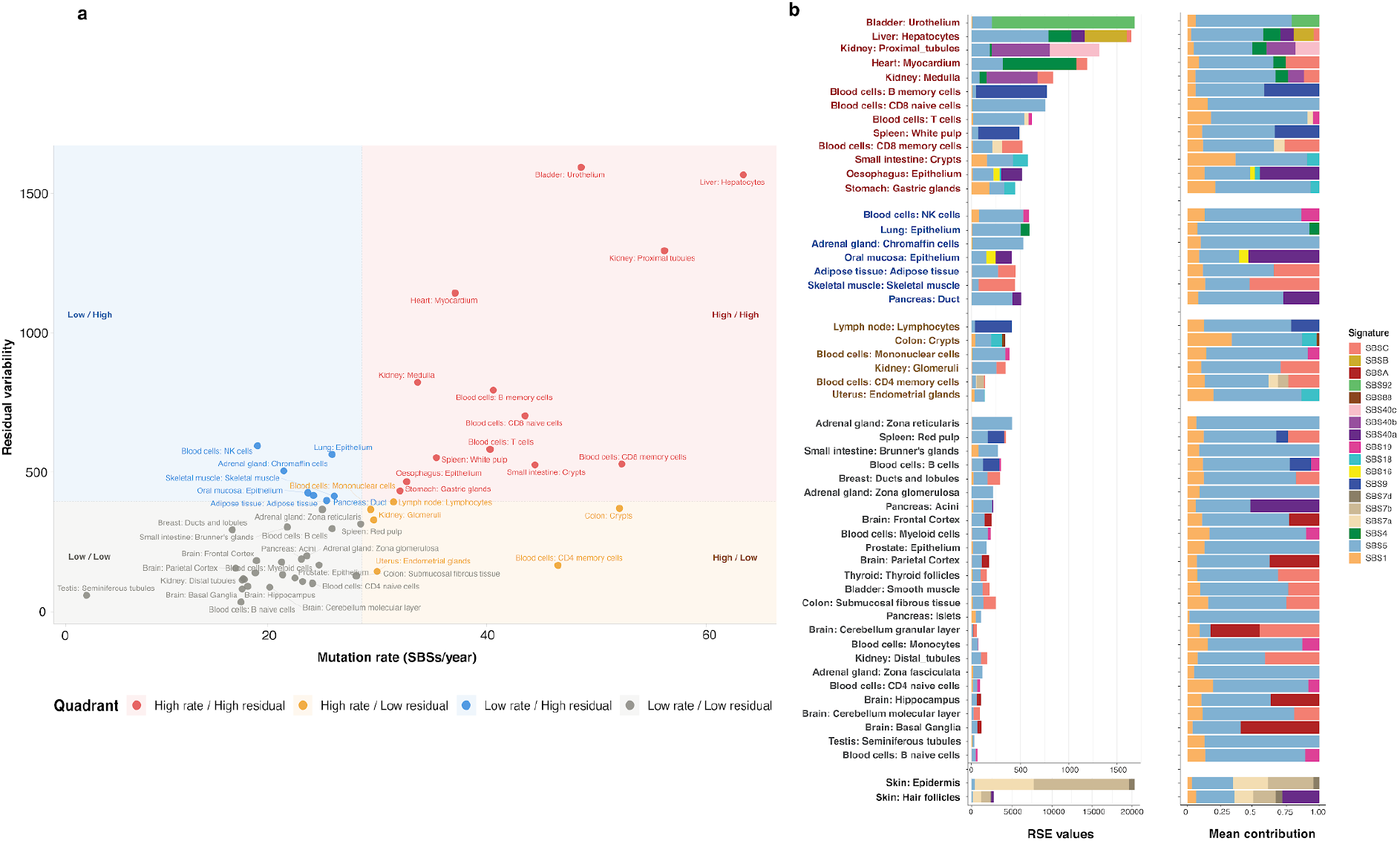
Residual variability in somatic mutation burden between cell types. (a) Quadrant scatter plot of mutation rate (SBS/diploid genome/year, x-axis, derived from the slope of the linear age-regression in Figure 2) against residual variability (RSE, y-axis, Methods) for each cell type. Cell types are partitioned into four quadrants by dashed threshold lines: high rate/high residual (red), high rate/low residual (orange), low rate/high residual (blue), and low rate/low residual (grey), and coloured/labelled accordingly. Quadrant thresholds are defined by mean of RSE and mean of mutation rates. (b) RSE values (left) and mean contribution of individual mutational signatures (right, coloured as in Figure 3) to residual variability for each cell type, ranked by decreasing RSE (top to bottom), illustrating that high residual variability in cell types such as bladder urothelium, liver hepatocytes, kidney proximal tubules and skin is predominantly attributable to variable signature exposures (for example SBS4, SBS92 and SBS7a/b/d) reflecting differences in exogenous mutagen exposure between individuals.

Mutational signature analysis indicates that the high interindividual variability of mutation burdens observed in some cell types is predominantly due to variation in intensity of exogenous exposures, with SBS4 mutation burdens caused by tobacco smoking in hepatocytes and cardiac myocytes, SBS92 due to tobacco smoking in bladder urothelium, and SBS7a, SBS7b and SBS7d due to ultraviolet light exposure in skin epidermis and hair follicles accounting for their high RSE values (Figure 4b). Variation in ultraviolet light exposure may also contribute to the relatively high interindividual variation of some T lymphocyte subtypes. By extension, therefore, it is plausible that SBS40b, which makes a substantial contribution to the RSE of mutations in kidney proximal tubules, could also be attributed to systemically circulating, currently unidentified mutagens to which individuals are variably exposed. However, other explanations of the interindividual variation of SBS40b, such as differences in inherited variation or impaired kidney function, cannot be excluded.

## DISCUSSION

This survey of mutation rates and mutational signatures charts the landscape of mutagenesis in normal human cells. Although comprehensive, it is not complete. The cell types examined constitute a subset of those known to exist through microscopy or single-cell transcriptomics and, despite enrichment by microdissection and flow sorting, the sequenced cell populations remain mixtures of cell types. Previously reported rare, low-amplitude, or geographically restricted mutational signatures in normal cells are not represented^11,20,26^ and RE-NanoSeq does not detect genome rearrangements or copy number changes. Future generations of survey may address these shortcomings by using whole-genome single-cell DNA sequencing, combined with transcriptome or methylome analysis of the same cells to classify cell type, applied to more exhaustive tissue sampling from larger numbers of people with global representation.

All cell types, including those that are postmitotic, show mutation accumulation during life. Some postmitotic/infrequently dividing cell types also show high mutation burdens from exogenous mutagenic exposures (for example, cardiac myocytes and SBS4 due to tobacco smoke mutagens). Thus, mutational processes of both endogenous and exogenous origins may not require DNA replication at cell division. In most cell types, a major component of mutation accumulation occurs linearly, indicating that, for this component, the underlying levels of DNA damage, rates of DNA repair, and fidelity of DNA replication remain the same throughout life. Differences in mutation rates between cell types are influenced by multiple factors, including both intrinsic biological features, such as cell division rate, and by exogenous mutagen exposures. Notably, mutagenesis due to exogenous exposures is not necessarily restricted to the directly exposed body surface, and intrinsic biological features of cell types may influence the extent of mutagenesis caused by a circulating mutagen. The extremely low mutation rates of spermatogonia and sperm suggest that endogenous mutagenesis can be suppressed if evolutionarily desirable but the mechanisms responsible are unknown^8^. The high mutation rates and high degrees of interindividual mutation burden variation in kidney proximal tubule cells and hepatocytes may be caused by currently unknown circulating mutagens, in a similar manner to their reporting of mutations due to the known circulating mutagens, aristolochic acids and aflatoxins^35^.

An extensive repertoire of mutational signatures is present in normal cells with contributions ranging from one to all cell types. For some, it remains uncertain whether they are of exogenous or endogenous origin and, for many, understanding of their underlying mechanisms of DNA damage/modification, DNA repair, and DNA replication is rudimentary. SBS5 accounts for most somatic mutations in the human body and for most sequence variation transmitted in the human germline. SBS5 is ubiquitous among cell types in humans and is found in all mammalian species studied, suggesting that it is of endogenous origin^37^. It generates mutations continuously through life at a more-or-less constant rate, showing modest variation between cell types, operates in postmitotic cells, is influenced only to a limited extent by cell division rates, is suppressed in spermatogonia and appears elevated in hepatocytes. Although generally assumed to be of endogenous origin, little is known about its underlying mutational process^38^. Elucidating the mutational process underlying SBS5 is a high research priority emerging from this and other studies.

The landscape of mutagenesis across normal human cell types potentially offers new perspectives on longstanding biological questions. For example, most cancers are thought to be caused by somatic mutations but different normal cell types show markedly different rates of conversion into cancer (Supplementary Note 6). Among the many factors that could account for these differences in cancer risk, a plausible candidate is the normal cell mutation rate. Indeed, the high incidence of skin epidermal cancers in sun-exposed body areas is likely to be, at least partially, explained by its extremely high mutation rate due to ultraviolet light-initiated mutagenesis. However, among other normal cell types, there is remarkably limited correlation between normal cell mutation rate and cancer risk. Although this does not preclude some influence, it suggests that the replicative potential of cell types, sizes of their cell populations, timing of mutations during life, tissue structure, patterns of normal somatic evolution, and other factors are stronger determinants of their variation in cancer incidence rates^20^.

The adult human is composed of ∼4 x 10^13^ cells. Knowledge of the range of mutation rates of its constituent cell types now enables estimation that there are ∼10^17^ somatic SBS mutations and ∼10^15^ non-synonymous single-base substitutions in protein-coding genes in the body of a 60-year-old individual. Assuming that mutations are more-or-less randomly distributed across the genome, each position in the genome is mutated ∼30 million times. Nature’s relentless experiment in saturation mutagenesis is known to contribute to cancer development but further consequences for human disease pathogenesis may yet emerge^18^. This comprehensive survey of mutation rates and mutational signatures in normal human cells provides a foundation for understanding somatic evolution in health and disease.

## METHODS

### Sample Collections and Ethics

All human biological material and associated clinical/demographic data used in this study were obtained with the approval of the relevant national or institutional research ethics bodies. Samples were contributed by 12 collecting institutions and biobanks across the United Kingdom, the Netherlands, Japan and the United States, under 16 independent ethical approvals, yielding a total cohort of 172 donors.

Samples collected at Cambridge University Hospitals NHS Foundation Trust were obtained under two approvals granted by the National Research Ethics Service (NRES) Committee East of England: reference 11/EE/0011 (1 donor), reference 11/EE/0155, approved 13/06/2011 (2 donors). Three breast samples were provided by Head Breast Pathology Group, The University of Queensland, under reference 20/PR/0905, approved by the London – Harrow Research Ethics Committee on 26/01/2021. Samples from Addenbrooke’s Hospital, part of Cambridge University Hospitals NHS Foundation Trust, were collected under reference 20/NI/0109, approved by the Research Ethics Committees Northern Ireland on 18/08/2020 (21 donors), and under reference 23/EE/0198, approved by the NRES Committee East of England on 17/10/2023 (7 donors); the same 23/EE/0198 approval also covered collection from Royal Papworth Hospital NHS Foundation Trust (21 donors).

Material from the Cambridge Biorepository for Translational Medicine (CBTM) was obtained under three separate NRES Committee East of England approvals: reference 15/EE/0152, approved 13/05/2015, covering three collection batches totalling 27 donors, among which three blood samples and 11 oesophagus samples were used in previous publication^39,40^); reference 15/EE/0351, approved 08/12/2015 (1 donor); and reference 16/EE/0227 (9 donors which were included in previous publication^12^), approved 29/07/202. Blood samples from the Cambridge Blood and Stem Cell Biobank (CBSB) and the National Institute for Health and Care Research (NIHR) BioResource were collected under reference 18/EE/0199, approved by the NRES Committee East of England on 15/07/2019 (10 donors, among which 2 donors were included in previous publication^27,39^). Paediatric samples were obtained from the Children’s Cancer and Leukaemia Group (CCLG) Tissue Bank under reference 08/H0405/22+5, approved by the East Midlands – Derby Research Ethics Committee on 17/04/2008 (1 donor). Material from the MCRC Biobank, held at The Christie NHS Foundation Trust, was collected under reference 22/NW/0237, approved by the North West – Greater Manchester South Research Ethics Committee on 30/08/2022 (1 donor), and material from Imperial College London was collected under reference 23/WM/0022, approved by the West Midlands – Black Country Research Ethics Committee on 31/08/2023 (1 donor).

Samples and data contributed by institutions outside the United Kingdom were collected under local institutional review board or equivalent ethics committee approval at the collecting site and were transferred for use in this study under the terms of the corresponding material and data transfer agreements. Material from AMSBIO and the University of Utrecht was collected under reference 17/LO/1801, approved by the London – Surrey Research Ethics Committee on 26/10/2017 (21 donors which were included in a previous publication^15^). Material from the International Agency for Research on Cancer was collected under reference 18/ES/0133, approved by the East of Scotland Research Ethics Service on 26/11/2018 (21 donors, among which seven donors also contributed to previous publications^8,41^). Material from Kyoto University was collected under reference 23/WS/0149, approved by the West of Scotland Research Ethics Service on 01/12/2023 (5 donors). Material from Memorial Sloan Kettering Cancer Center was collected under Institutional Review Board protocol IRB 15-021, approved 06/02/2020 (20 donors).

Written informed consent for the collection, storage and use of tissue, blood and associated clinical data in ethically approved research, including molecular and genomic analysis, was obtained from all donors (or from a parent/legal guardian in the case of paediatric donors contributing to the CCLG Tissue Bank) prior to sample collection.

Further metadata for all samples can be found in Supplementary Table 1.

### Laser capture microdissection of solid tissues

After a brief thaw at 4 °C, frozen solid specimens were fixed using the PAXgene Tissue FIX reagent (PreAnalytiX). Fixation was carried out at room temperature (approximately 20–25 °C) for either 12 hours (small samples, maximum 4 x 15 x 15 mm) or 24 h (large samples, maximum 20 x 20 x 20 mm). Afterwards, the fixing reagent was replaced with PAXgene Tissue STABILIZER, and samples were stored at −20 °C prior to downstream processing.

For histological preparation, the fixed samples were transferred to standard cassettes, processed using a Tissue-Tek VIP system (Sakura Finetek), and embedded in paraffin blocks. Sectioning was performed on an Accu-Cut SRM 200 microtome (Sakura Finetek). Reference slices were cut at a thickness of 4 μm and mounted onto Superfrost Plus slides (Thermo Fisher Scientific) before staining and permanent coverslipping. LCM sections were cut at 16 μm thickness, placed on polyethylene naphthalate (PEN) membrane slides, stained with hematoxylin and eosin, and temporarily coverslipped.

37 different structures from 23 solid tissues were isolated into a skirted 96-well LoBind PCR plate (Eppendorf, UK) using an LMD7 microscope (Leica Microsystems). Two to four LCM samples were collected for each tissue structure. They were then lysed in 20 μl of proteinase K from the PicoPure DNA Extraction kit (Arcturus). Samples were incubated at 65 °C for 12 hours, then the proteinase was denatured at 75 °C for 30 min. A matched normal (blood or another tissue type) sample was also dissected across structures for each patient to avoid clonal expansion bias.

### Blood and Brain single cell sorting

Peripheral blood mononuclear cells (PBMCs) were isolated via density gradient centrifugation using Lymphoprep (STEMCELL Technologies) after a 1:1 dilution of whole blood with PBS. Red blood cells (RBCs) and granulocytes were simultaneously separated during this step. Residual RBCs in the mononuclear fraction were lysed using RBC Lysis Buffer (BioLegend) at 4 °C for 15 min. CD34+ cells were enriched using the EasySep human whole blood CD34+ selection kit (STEMCELL Technologies) via a single round of immunomagnetic selection per the manufacturer’s instructions. Conversely, bone marrow mononuclear cells (BMMCs) were not pre-selected before fluorescence-activated cell sorting (FACS).

Mononuclear or enriched CD34 cells were pelleted and resuspended in PBS supplemented with 3% FBS containing the respective fluorochrome-conjugated antibody panels. To ensure single, live-cell gating, cells were stained with Zombie Aqua fixable viability dye (BioLegend). For Batch 1 (donors sourced from the NIHR Bioresource), cells were stained with anti-CD3/FITC and anti-CD19/BV785 to isolate T-cell, B-cell, and myeloid fractions. For Batch 2 (donors sourced from the Cambridge Biorepository for Translational Medicine and Cambridge Blood Spark Biobank), cells were incubated with a multi-color panel comprising anti-CD3/FITC, anti-CD14/BV605, anti-CD90/PE, anti-CD49f/PE-Cy5, anti-CD38/PE-Cy7, anti-CD33/APC, anti-CD19/Alexa Fluor 700, anti-CD34/APC-Cy7, and anti-CD45RA/BV421 to resolve monocytes, bulk T cells, CD4+ memory/naïve subsets, CD8+ memory/naïve subsets, bulk B cells, and memory/naïve B cells. All staining was performed in the dark at 4 °C for 30 min, followed by washing, centrifugation, and final resuspension in 3% FBS/PBS.

Target populations were sorted into 96-well plates using either a BD FACSAria III or BD FACSAria Fusion flow cytometer (BD Biosciences) at the NIHR Cambridge BRC Cell Phenotyping Hub. Genomic DNA was subsequently isolated using either the DNeasy 96 Blood & Tissue Kit (Qiagen) or the PicoPure DNA Extraction Kit (Arcturus). Cell lysates were stored at −20 °C prior to downstream sequencing library preparation.

Neuronal populations were isolated and sorted from frozen brain tissue specimens provided by Memorial Sloan Kettering Cancer Centre (MSKCC). All sample handling and processing were conducted within an AirClean PCR Workstation. Briefly, approximately 250–400 mg of frozen brain tissue was homogenised using a sterile Dounce tissue grinder in a sterile, non-ionic surfactant-based lysis buffer (250 mM Sucrose, 25 mM KCl, 5 mM MgCl_2_, 10 mM Tris buffer, pH 8.0, 0.1% (v/v) Triton X-100, 3 μM DAPI, Nuclease Free Water). The resulting homogenate was filtered through a 40 μm cell strainer and centrifuged at 800 g for 8 min at 4 °C. The nuclear pellet was gently resuspended in 200 μl of FACS buffer (0.5% BSA, 2mM EDTA) and incubated on ice for 10 min.

Following a subsequent centrifugation (800 g for 5 min at 4 °C), the sample was incubated with an anti-NeuN antibody (neuronal marker; 1:500, PE-conjugated, clone A60, Milli-Mark, Merck Millipore^TM^) for 40 min. Cells were centrifuged again (800 g for 5 min at 4 °C), washed with 1X permeabilisation buffer (from the Foxp3 / Transcription Factor Staining Buffer Set; eBioscience^TM^), and centrifuged at 1300 g for 5 min without braking to maximise nuclear recovery. Intranuclear staining was then performed for 40 min in 1X permeabilisation buffer using anti-PU.1 (microglial marker; 1:50, Alexa Fluor 647-conjugated, clone 9G7, Cell Signaling Technology) and anti-Olig2 (oligodendrocyte marker; 1:1,000, Alexa Fluor 488-conjugated, ab109186, Abcam).

After a final wash with FACS buffer, nuclei were sorted using a BD FACSAria flow cytometer (BD Biosciences). Flow cytometry data were acquired using FACSDiva software (v8.0.1; BD Biosciences), and post-acquisition analysis was performed using FlowJo (v10.6.2; FlowJo, LLC). Nuclei were collected into 1.5-ml Eppendorf tubes containing 100 μl of sterile PBS. For each target lineage, 100,000 nuclei were isolated with greater than 95% purity. Collected nuclear suspensions were pelleted by centrifugation at 6000 g for 20 min and immediately processed for genomic DNA extraction using the QIAamp DNA Micro Kit (Qiagen) according to the manufacturer’s instructions.

### RE-NanoSeq library preparation and sequencing

The RE-NanoSeq protocol was conducted as previously described (Abascal et al., 2021). Briefly, 20 μl of DNA was purified with a 1:1 mixture of nuclease-free water (NFW) and AMPure XP beads (Beckman Coulter) and eluted in 20 μl of NFW. DNA fragmentation was performed by incubating the bead suspension with 2.5μl 10X CutSmart buffer, 0.5 μl HpyCH4V (5 U μl^-1^, New England Biolabs), and 2μl NFW at 37 °C for 15 min, followed by 2.5X AMPure XP bead purification and 15 μl of NFW resuspension. A-tailing was conducted by incubating 10 μl of fragmented DNA with 5 μl reaction buffer (1.5 μl 10X NEBuffer, 0.15 μl Klenow fragment, 1.5 μl of 1 mM equimolar dATP/ddBTPs and 1.85 μl NFW) at 37 °C for 30 min. For adapter ligation, 22.4 μl of master mix, containing 2.24 μl 10X NEBuffer 4, 3.74 μl 10 mM ATP, 0.33 μl 15 μM xGen Duplex Seq Adapters (IDT, 1080799), 0.56 μl T4 DNA ligase (400 U μl−1, New England Biolabs) and 15.53 μl NFW, was added directly to the A-tailing product, incubated at 20 °C for 20 min, and purified using 1X AMPure XP beads.

DNA libraries were quantified using qPCR (KAPA Library Quantification Kit, Roche) and diluted in 25 μl of NFW to a target concentration of 0.3-0.5 fmol to optimise the parental strand duplication rate, yielding approximately 300 million reads (20-30X coverage). PCR amplification was performed using NEBNext Ultra II Q5 Master Mix (New England Biolabs) and unique dual index primers. Thermocycling conditions were set up at 98 °C for 30 s; n cycles of 98 °C for 10 s and 65 °C for 75 s; 65 °C for 5 min; and a 4 °C hold. The cycle number (n) was adjusted according to the input concentration. Amplicons were purified via two sequential 0.7X AMPure XP clean-ups, quantified with the AccuClear Ultra High Sensitivity dsDNA Kit (Biotium), and sequenced on an Illumina NovaSeq 6000 platform (150-bp paired-end).

### Variant calling and filtering

RE-NanoSeq and whole-genome sequencing matched normal data were processed using the reproducible computational workflow as described on https://github.com/cancerit/NanoSeq. De-multiplexed FASTQ files were trimmed off adapters, and three-nucleotide barcodes were appended to the read headers. All reads were aligned to the hs37d5 reference genome using BWA-MEM v.0.7.17^42^ with the -C flag. Alignments were sorted, duplicate-marked, and annotated using *biobambam2* v.2.0.86^43^. Reads that failed quality control, were unmapped or yielded secondary alignments, or were improper pairs or optical duplicates were excluded.

For SBS identification, high-confidence base calls required read bundles to contain at least two duplicates from each parental strand, a consensus base quality score greater than 60, and a minimum difference between primary alignment score (AS) and the secondary alignment score (XS) equal or greater than 50 across both RE-NanoSeq and matched normal samples. Grouped reads with an average of greater than 2 mismatches, 5’ clippings, improper pairs or variants within 8 bp of read ends were discarded. Somatic variants were isolated by filtering germline variants against a standard 15x WGS matched normal for each donor. Regions overlapping common SNPs and predefined low-complexity noise sites were masked.

For indel calling, the pipeline identified read bundles with indels localised in at least 90% of forward and reverse reads using *SAMtools* and *BCFtools* v.1.19^44^. Any read bundles with an AS-XS score ≥ 50, without 5’ clipping, and at least 16X coverage at the given site in the matched normal are kept. Indel candidates within 10 bp of read ends were not called. To eliminate germline contamination and local mapping artefacts, candidate loci were cross-referenced against a ±5-bp window in the matched-normal data using the *bam2R* function from the *deepSNV* R package^45^. Moreover, indels were discarded if high-quality supporting reads in the matched-normal sample exceeded a 1% allele frequency threshold.

### Cross-contamination detection

Sample authenticity and cross-contamination with foreign human DNA were evaluated using verifyBAMID2 (v1.0.6)^46^, which estimates contamination levels by benchmarking BAM files against a reference panel of known population allele frequencies. Samples with a contamination coefficient (FREEMIX values) exceeding the standard threshold of 0.005 (corresponding to greater than 0.5% contaminating reads) were excluded from downstream analysis Additionally, when a sample is likely contaminated or paired with the wrong matched normal, many common SNPs are detected, leading to a high ratio of SNP masked variants to passed variants. After investigation, samples with greater than 10-fold more mask variants were also removed from downstream analysis.

### Mutation burden correction

Single-base substitution (SBS) rates are strongly influenced by trinucleotide sequence context, especially when RE-NanoSeq targets only 30% of the human genome due to restriction enzyme and size-selection constraints. Therefore, raw SBS counts were mathematically corrected to enable cross-dataset comparisons. For each of the 96 trinucleotide contexts, a sample-specific correction factor was calculated as the ratio of the context frequency in the full GRCh37 reference genome to its observed frequency within the sequenced regions of that sample. The corrected SBS burden per bp was subsequently determined by dividing the total corrected substitution counts across all 96 contexts by the total number of reference and variant bases called.

Conversely, context-specific corrections were not applied to insertion–deletion (indel) variants. Due to the structural diversity of the 83 indel subtypes, such adjustments are exceedingly complex; furthermore, indel formation kinetics are substantially less influenced by restriction-enzyme sequence specificities than SBS mechanisms. Consequently, the raw indel burden per bp was calculated directly by dividing total unadjusted indel calls by the total duplex coverage for each sample.

To compare the RE-NanoSeq mutation burdens to the standard WGS mutation burdens, both the corrected SBS burden and the raw indel burden per bp were scaled to absolute mutation loads of a reference human diploid genome.

### Mutation rate analysis

To characterise the relationship between donor age and somatic mutation burden across the dataset, we first compared linear, quadratic, and exponential mixed-effects models, with donor as a random effect to account for non-independence of samples from the same individual (Supplementary Note 2). Linear and quadratic models were fitted by maximum likelihood using lmer (R package *lme4*^47^); the exponential model was fitted using non-linear mixed-effects regression (R package *nlme*^48^). Models were compared by Akaike Information Criterion (AIC). The linear model provided the best fit (AIC = 7358.8), with a robustly significant fixed effect of age (p = 4.29 × 10⁻^15^). The quadratic model showed a small increase in AIC (ΔAIC = 3.6) and its second-order term was not significant (p = 0.52), while the exponential model was decisively rejected (ΔAIC = 43.5). We therefore modelled somatic mutation burden as accumulating linearly with age in downstream analyses.

Mutation rates for each cell type were then estimated using linear mixed-effects regression (R package *lme4*^47^), with mutation burden (burden_wg) as the response variable and donor age as a fixed effect (Supplementary Note 2). To account for the hierarchical structure of the data, *individual* was taken as a random intercept, while *cell type* was parameterised with random effects on slopes for age *(0 + age | cell type)*. To evaluate the statistical significance of the random structures (*cell type* and *individual*) on mutation burden, multiple models were compared using the anova function in R, with maximum likelihood estimation (*REML = FALSE*; Supplementary Note 2). Parameter estimates for the final selected model were subsequently derived using Restricted Maximum Likelihood *(REML = TRUE)* as below:

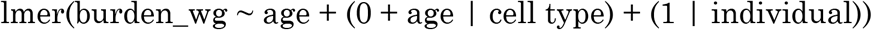

Confidence intervals (95% CIs) for the resulting regression trends were estimated via bootstrapping. Briefly, 1,000 bootstrap replicates were generated for each model configuration. Predictions and corresponding standard errors were modelled over the age range 0 to 100 years, and the 95% CIs were defined as the 2.5th and 97.5th percentiles of the bootstrapped prediction distributions.

To quantify the goodness of fit and explanatory power of our linear mixed-effects model, we calculated the conditional coefficient of determination (R^2^). The conditional R^2^ represents the proportion of variance explained by the entire model, including both fixed effects *(age)* and the random effects *(cell type and individual)*. Mathematically, the conditional R^2^ is calculated by dividing the sum of the fixed-effects variance and all random-effects variance components by the total theoretical variance of the model.

Statistical significance of the age-related accumulation rate was initially evaluated using a two-sided Student’s t-test in a simple linear model for each structure. To account for multiple hypothesis testing across the various anatomical structures, obtained p-values were subsequently adjusted using the Benjamini–Hochberg false discovery rate (FDR) procedure. Findings were considered statistically significant at an adjusted p-value < 0.05. Residual standard errors (RSE) were computed as the square root of the residual sum of squares divided by the degrees of freedom. These values were used to quantify the absolute variation in mutation burden around the predicted regression lines, serving as a measure of structure-specific biological deviations from steady, age-dependent accumulation.

### Mutational signature extraction and deposition

For SBS signatures, mutations are first classified into 6 categories, using pyrimidine bases as a reference (C > A, C > G, C > T, T > A, T > C and T > G). Then they are subclassified into 96 trinucleotide contexts by adding one base before and one base after the mutated position. Small insertions and deletions (INDELS) were grouped into 83 classes according to whether the mutation is an insertion or a deletion, its length, how many times the reference sequence repeats, or whether it exhibits microhomology.

De novo mutational signatures were extracted independently using three algorithms to cross-validate discovered patterns: a Bayesian hierarchical Dirichlet process (HDP; v0.1.5)^49^, non-negative matrix factorisation (NMF) via SigProfilerExtractor (v1.1.23)^50^, and minimum-volume NMF via MuSiCal (v1.0.0)^51^ (Supplementary Note 3). To eliminate background noise and prevent the noisy signatures, samples with fewer than 100 SBSs or 20 indels were excluded from downstream extraction workflows.

For HDP^49^, the algorithm was initialised without hierarchical structural constraints or informative prior settings across 20 independent, parallel Markov chain Monte Carlo (MCMC) chains. The alpha and beta parameters for the clustering were set to 1. Posterior sampling was conducted using a Gibbs sampler with a 30,000-iteration burn-in period. Subsequently, a total of 100 posterior samples were collected at intervals of 200 iterations to minimise autocorrelation. Following each Gibbs iteration, three concentration-parameter update cycles were performed, ultimately resolving 21 de novo somatic signatures.

For SigProfilerExtractor^50^, NMF-based factorisations were performed over a predefined range of solutions, from 2 to 20 signatures *(min_sigs = 2, max_sigs = 20)*. To robustly estimate solution stability and account for sampling variance, 500 independent factorisation replicates were performed at each solution using Poisson-distributed bootstrapped versions of the original mutation counts. The optimal solution was selected by objectively balancing the maximisation of the average silhouette width (representing signature stability) against the minimisation of the mean reconstruction error (measured via cosine similarity). This selection framework identified the most stable solution of 9 stable de novo signatures.

For MuSiCal^51^, signature extraction was performed utilising its minimum-volume NMF objective function, which incorporates a regularisation penalty to enforce compact signature geometry and diminish ambient noise assignment. The extraction was executed under default parameterisations across 100,000 algorithmic replicates. The final number of signatures was selected based on MuSiCal’s internal cross-validation and reconstruction error minimisation criteria, which identified 11 high-confidence de novo signatures.

Fifteen high confidence de novo signatures were strictly selected based on two criteria: (1) if they were independently extracted by at least two distinct algorithms, or (2) if they exhibited greater than 0.9 cosine similarity to reference signatures in the COSMIC database (v3.4, <u>cancer.sanger.ac.uk/signatures</u>) (Supplementary Note 3). These signatures were further deconvoluted into known COSMIC signatures. A subset of 12 signatures is composed of SBS1, SBS2, SBS4, SBS5, SBS7a, SBS7b, SBS7d, SBS13, SBS16, SBS17b, SBS18, SBS19,

SBS40a, SBS40b, SBS40c, SBS88, SBS92. HDP8 with cosine similarity between the original and reconstructed signatures of less than 0.9 were considered unclassified and taken forward as SBSA. HDP3 showed a close match to SBS5 (cosine similarity 0.9) and exhibited high prevalence in the hepatocytes, indicating a tissue-specific mutational process. An additional signature extraction was conducted exclusively on liver samples using HDP to isolate a cleaner version of HDP3. Consequently, the signature, which resembled HDP3 and could not be decomposed into any known signatures, was designated as SBSB. Similarly, a cleaner version of HDP8 was extracted from a subdataset of skeletal muscle, heart, and cerebellum samples and was named SBSC (Supplementary Note 3).

Since SBS5, SBS16, SBS92, SBSA, and SBSB shared a high rate of T>C mutations at ApT dinucleotides, we investigated whether the underlying mutational processes are similar. We studied the transcriptional strand bias in trinucleotide and pentanucleotide contexts of T>C mutations by examining signatures extracted from 288 and 1536 classes, respectively. The comparison across the five signatures is described in the Extended Figure 6.

### Mutational signature attribution

Samples of the same cell type from the same donor were merged into one to increase the number of absolute mutations for attribution. To prevent algorithmic overfitting and minimise false-positive signature assignments, a two-tiered attribution workflow was implemented. In the initial tier, we examined the raw HDP exposures and mapped the specific HDP components to their corresponding decomposed signatures to construct the final sample-specific active lists (Supplementary Note 3). The attribution of signatures to each sample was performed using SigProfilerAssignment^50^ with 1,000 independent bootstrap iterations. To filter out uncertain and off-target attribution, a signature was strictly chosen for each cell type only if its lower 95% CI was higher than zero and was detected in at least two samples. Consequently, each cell type had a specific list of signatures for attribution. In the second tier, samples within each cell type were re-attributed exclusively against these validated signatures to produce final attribution.

### Residual variability decomposition by mutational signature

To partition the residual variability among individual signatures, a separate linear model of mutation burden against age was fitted independently for each signature within each cell type, and the per-donor residuals from each of these signature-specific models were extracted. For each cell type, residuals were arranged into a donor-by-signature matrix, and the covariance matrix between signatures’ residuals was calculated using pairwise-complete observations. The diagonal terms of this matrix represent the variance in residual mutation burden attributable to each signature considered independently, whereas the off-diagonal terms capture covariance between signatures arising, for example, from correlated or shared exogenous exposures. Total residual variance for each cell type was calculated as the sum of all terms of the covariance matrix (diagonal plus off-diagonal), and its square root (RSE_decomp) provides a covariance-based estimate of overall residual variability that can be cross-checked against the RSE obtained from the aggregate age-regression model (RSE_total).

The percentage contribution of each signature to residual variability was calculated as its diagonal (independent) variance term divided by the sum of diagonal variances across all signatures attributed to that cell type, expressed as a percentage. To enable direct, cross-cell-type comparison of the absolute contribution of each signature to residual mutation burden variability, these percentage contributions were rescaled onto the same absolute scale as RSE_total (each percentage multiplied by RSE_total/100), yielding an RSE-equivalent value for each signature within each cell type. Cell types were ordered by ascending total RSE for presentation and downstream visualisation.

## Acknowledgement

The authors thank K. Roberts, K. Smith, S. Austin-Guest and the staff of Sequencing Operations at the Wellcome Sanger Institute for their contribution; J. Allingham from Google Deepmind, S. Sunyaev from Harvard Medical School, L. Wylie and K. Dawson from Somatic Genomics Programme at the Wellcome Sanger Institute for useful discussions; S. Aitken from Yale Cancer Center for pathology review. We appreciate the patients involved in this study and their families. We thank NIHR BioResource volunteers for their participation, and gratefully acknowledge NIHR BioResource centres, NHS Trusts and staff for their contribution. We thank the National Institute for Health and Care Research, NHS Blood and Transplant, and Health Data Research UK as part of the Digital Innovation Hub Programme. The views expressed are those of the author(s) and not necessarily those of the NHS, the NIHR or the Department of Health and Social Care. We express our gratitude to the patients and staff at Royal Papworth Hospital NHS Foundation Trust for their cooperation, support, and contribution to this study. We acknowledge the Children’s Cancer and Leukaemia Group (CCLG) Tissue Bank, the CCLG centers and the Experimental Cancer Medicine Centres Paediatric Network for the collection and provision of tissue samples (project 2016 BS 05). The CCLG Tissue Bank is funded by Cancer Research UK and CCLG. We are grateful to the Cambridge Biorepository for Translational Medicine, the Cambridge Blood and Stem Cell, the Memorial Sloan Kettering Cancer Center, the Kyoto University, the University of Queensland, the University of Utrecht, the Addenbrooke’s Hospital, Cambridge, UK for providing the tissue used in this work.

## Funding

This work was supported by the Wellcome Trust grants 206194 and 220540/Z/20/A. For the purpose of open access, the authors have applied a CC-BY public copyright license to any author-accepted manuscript version arising from this submission. M.H. is supported by a CRUK Programme Foundation Award (DRCPFA-Jun22\100001) and an MRC research grant (MR/X00970X/1).

## Author Contributions

M.R.S., R.R. and M.H.P. conceptualized the project with support from K.S., K.T.A.M, E.J.B, D.R, F.G, K.K., R.L.A.W.B, L.M, R.H., S.B, M.H., P.J.C, P.H.J, I.M.; M.H.P. generated data and led the analysis. L.M.R.H., T.R.W.O., E.D., G.L.J., Y.W., J.C.F., R.V., Y.I., H.E.M, N.B., C.D., C.A.P., and H.M. contributed to sample recruitment and data generation. A.R.J.L. and P.A.N. contributed to sample preparation and project discussions. Pathology review was carried out by T.R.W.O.; R.S., F.A. and M.D.C.N. provided bioinformatics support. Y.H. provided histology support. H.J., T.H.H.C., S.M. contributed to discussions on mutational signatures. A.P.L. contributed to statistical discussions. L.O. provided sequencing support. L.H. and C.L. provided administrative support. I.M., R.R. and M.R.S. supervised the project. M.R.S., R.R. and M.H.P. wrote the manuscript, and all authors contributed to reviewing and editing it.

## Competing interests

P.J.C., I.M., and M.R.S. are cofounders; P.J.C. is an employee; and I.M., M.R.S. and F.A. have consulted for Quotient Therapeutics Ltd. M.H. is a consultant for Quotient Therapeutics, AstraZeneca and Boston Scientific and has received unrestricted scientific grants from Pfizer. L.M. is an employee of AstraZeneca. All other authors declare no competing interests.

## Data availability

The raw sequencing data generated in this study are deposited in the European Genome-phenome Archive (EGA) under accession number EGAD00001016070. All other data are available from the authors on request.

## Code availability

Mutation-calling algorithms are available through GitHub (https://github.com/cancerit/NanoSeq). The code used to carry out all analyses and make all plots is available at https://github.com/Phamhamy/Panbody_RE_NanoSeq.

---

**Extended Figure 1.**
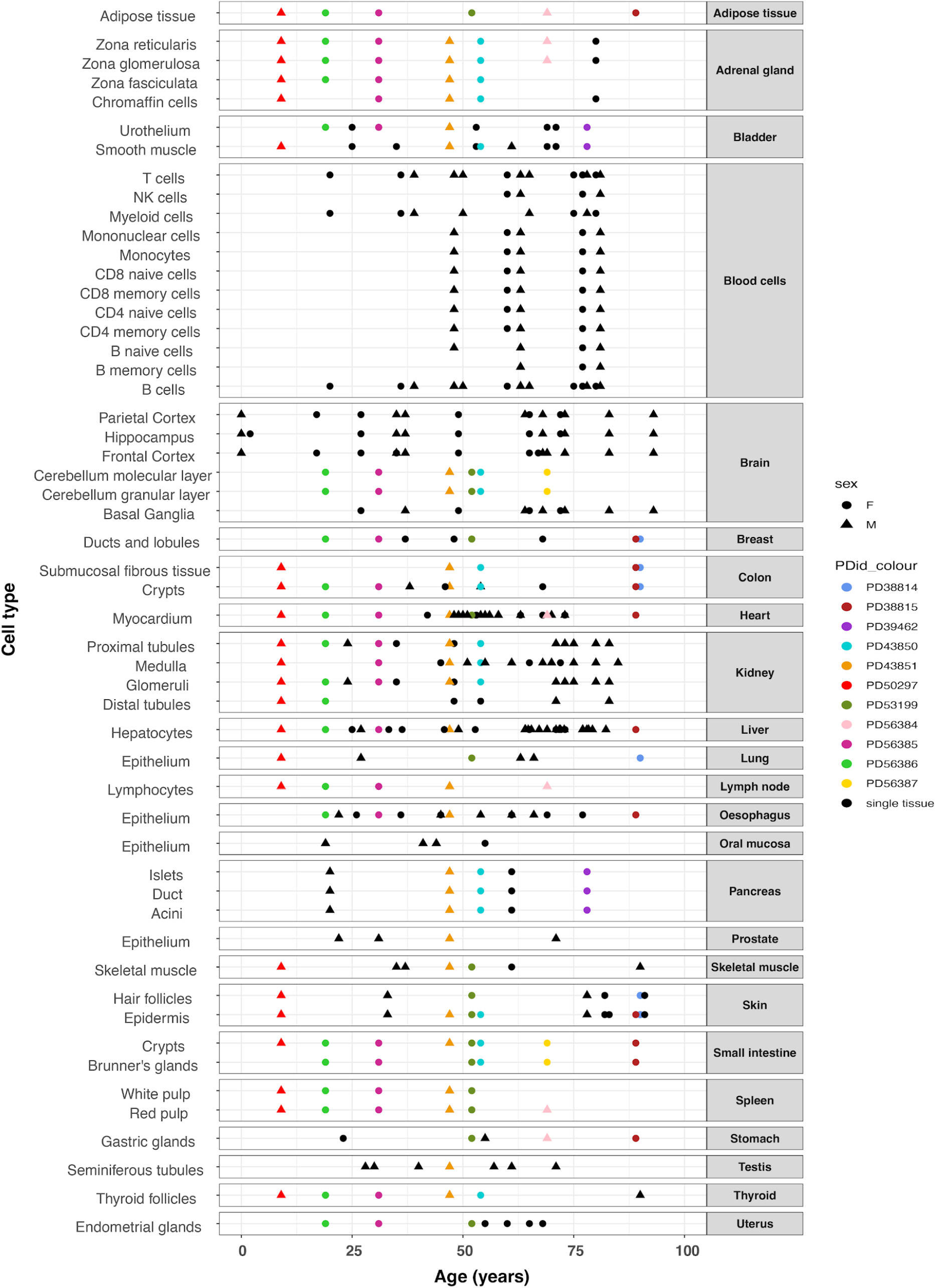
Cohort composition and sampling of tissue structures and cell types. Donor age (years, x-axis) for every sample analysed, grouped by tissue structure/cell type (y-axis) and tissue of origin (right-hand grey panels). Point shape denotes donor sex (circle, female; triangle, male). Points are coloured black where only a single tissue/cell type was obtained from that donor, or coloured by individual donor identifier (PDid, legend) where multiple tissue structures or cell types were sampled from the same donor, illustrating the extent of matched multi-tissue sampling across the cohort of 172 individuals (0–93 years).

**Extended Figure 2.**
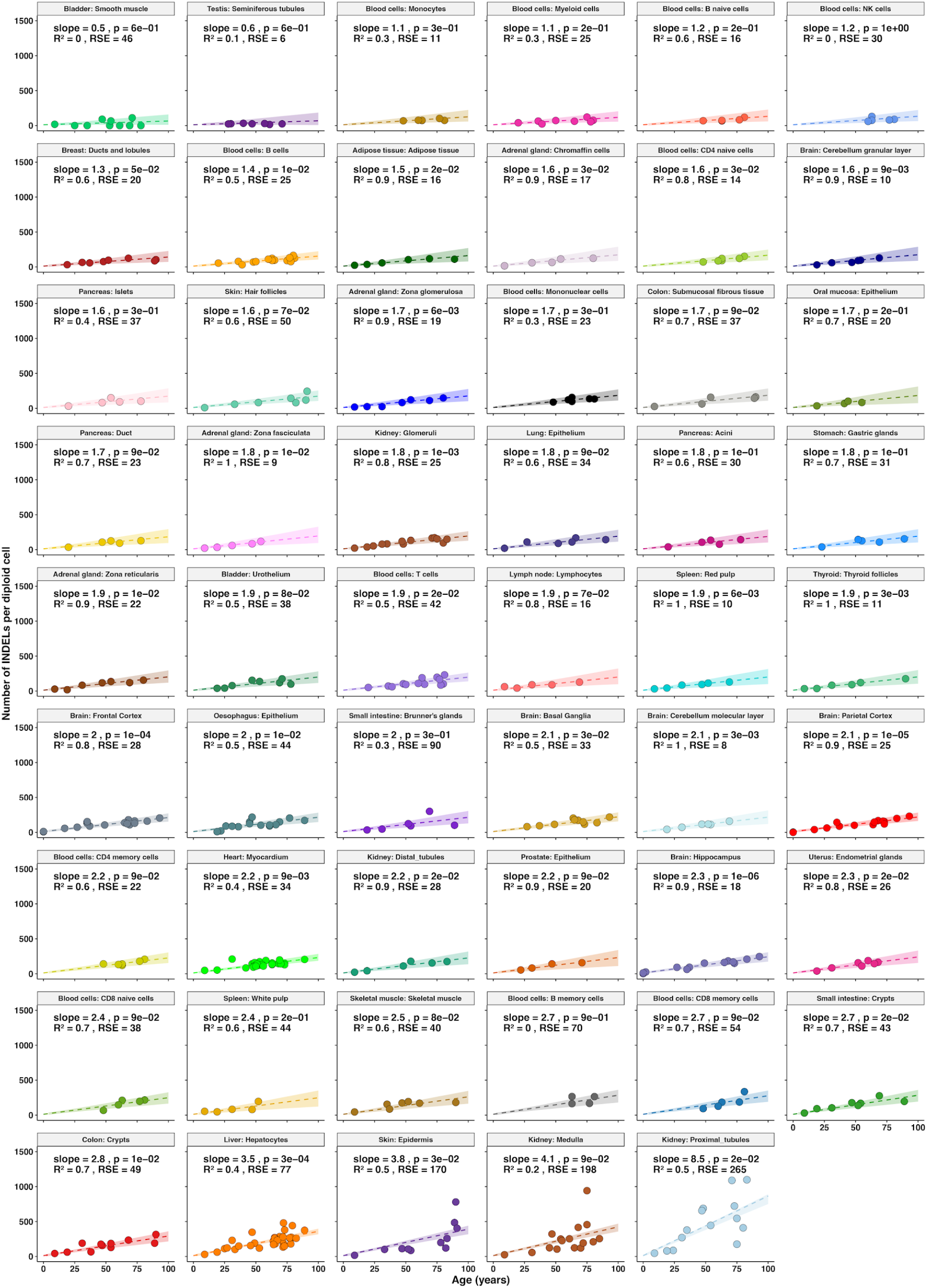
Linear accumulation of small insertions and deletions with age across normal cell types. Number of small insertion and deletion (ID) mutations per diploid cell plotted against donor age (years) for each of the 53 cell types, ordered by increasing regression slope (top left to bottom right), as in Figure 2. Each point represents one sample; dashed line, linear regression fit; shaded ribbon, 95% bootstrapped confidence interval. Slope (ID/diploid genome/year), FDR-adjusted two-sided t-test p-value, R² and residual standard error (RSE) are shown for each cell type. The ranking of cell types by ID mutation rate broadly mirrors the SBS ranking in Figure 2, except for kidney proximal tubule cells, which show a disproportionately high ID mutation rate (8.5 ID/year) relative to their SBS rate.

**Extended Figure 3.**
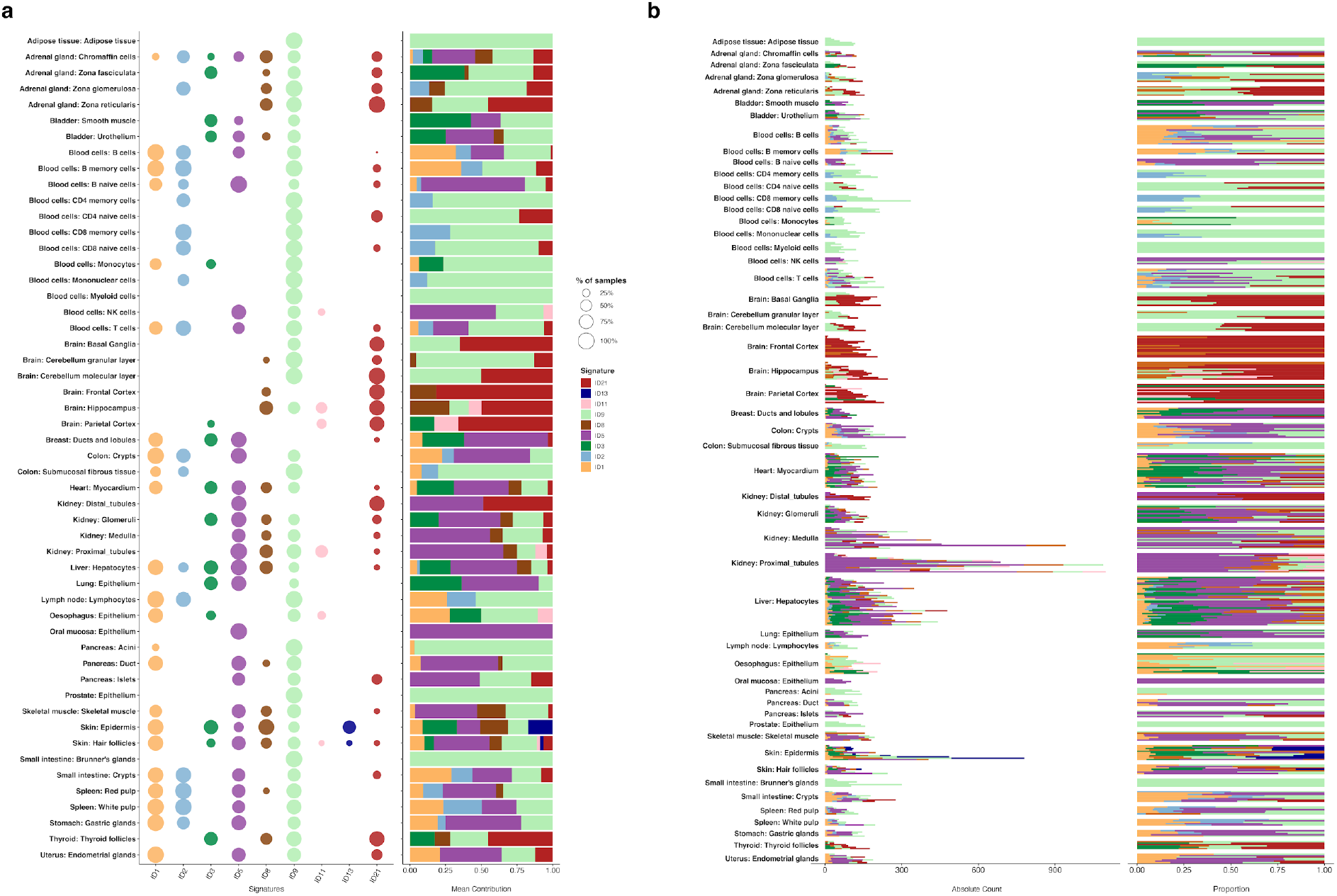
Landscape of small insertion/deletion mutational signatures across normal cell types. (a) Bubble plot showing the presence of each of the nine COSMIC ID mutational signatures (ID1, ID2, ID3, ID5, ID8, ID9, ID11, ID13, ID21; columns) across the 53 cell types/tissue structures (rows). Bubble size indicates the percentage of samples of a given cell type in which the signature was attributed; colour denotes signature identity. Stacked bars (right) show the mean proportional contribution of each signature per cell type. (b) Absolute mutation counts (left) and proportional contribution (right) of each ID signature for every individual sample, grouped by cell type, coloured as in a. ID9, of unknown aetiology, is near-ubiquitous across cell types and samples.

**Extended Figure 4.**
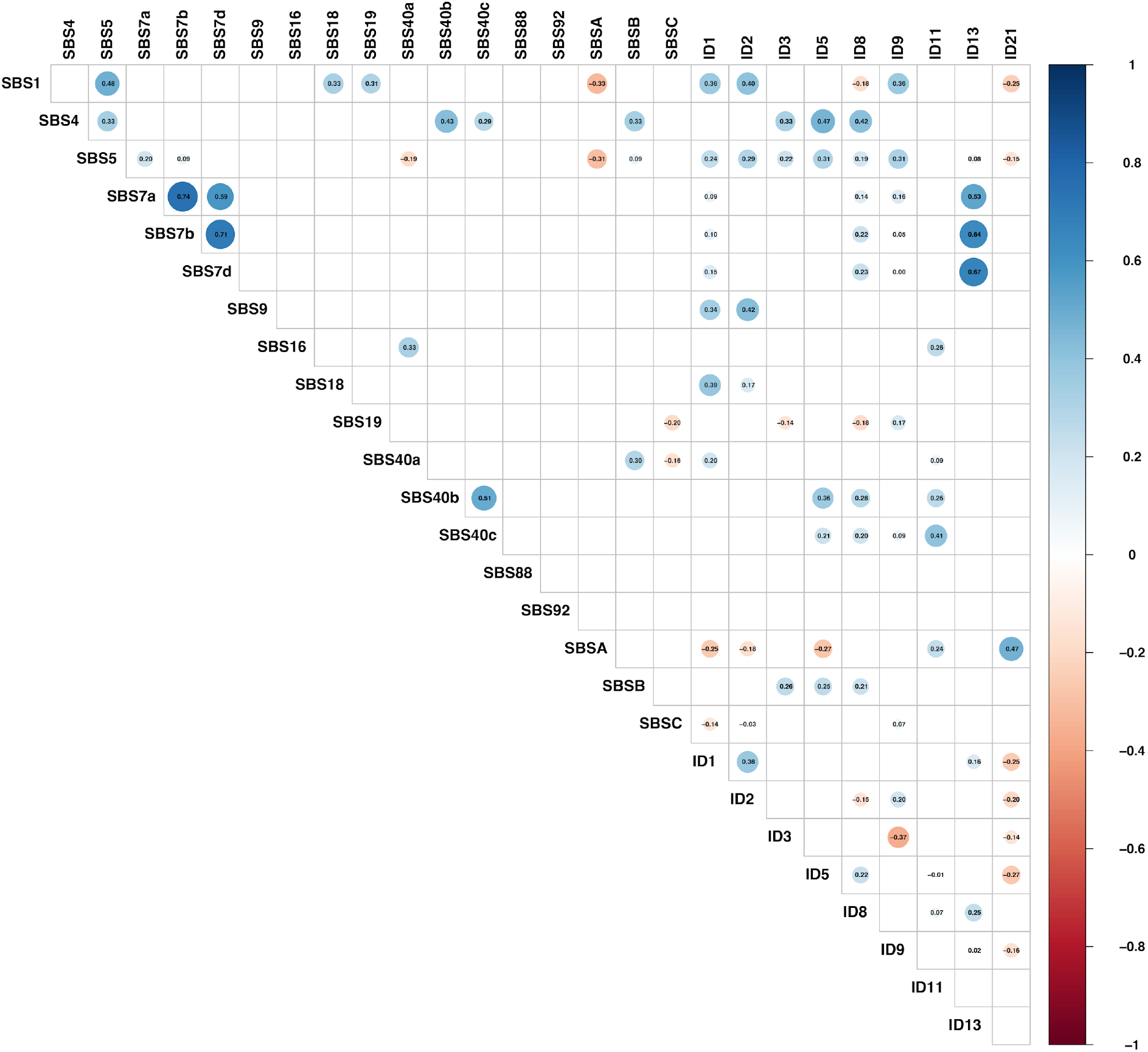
Pairwise correlations between mutational signature burdens. Correlation matrix (lower triangle) of per-sample mutation burdens attributed to each SBS (SBS1, SBS4, SBS5, SBS7a, SBS7b, SBS7d, SBS9, SBS16, SBS18, SBS19, SBS40a, SBS40b, SBS40c, SBS88, SBS92, SBSA, SBSB, SBSC) and ID (ID1, ID2, ID3, ID5, ID8, ID9, ID11, ID13, ID21) mutational signature across all samples and cell types. Circle size and colour intensity denote the magnitude of the Pearson correlation coefficient (scale, −1 to 1, right); blue, positive correlation; red, negative correlation. Numerical coefficients are shown within each circle. Statistical significance was assessed using cor.test() for each pair, and resulting p-values were adjusted for multiple comparisons using the Benjamini-Hochberg procedure. Blank cells indicate non-significant or negligible correlations.

**Extended Figure 5.**
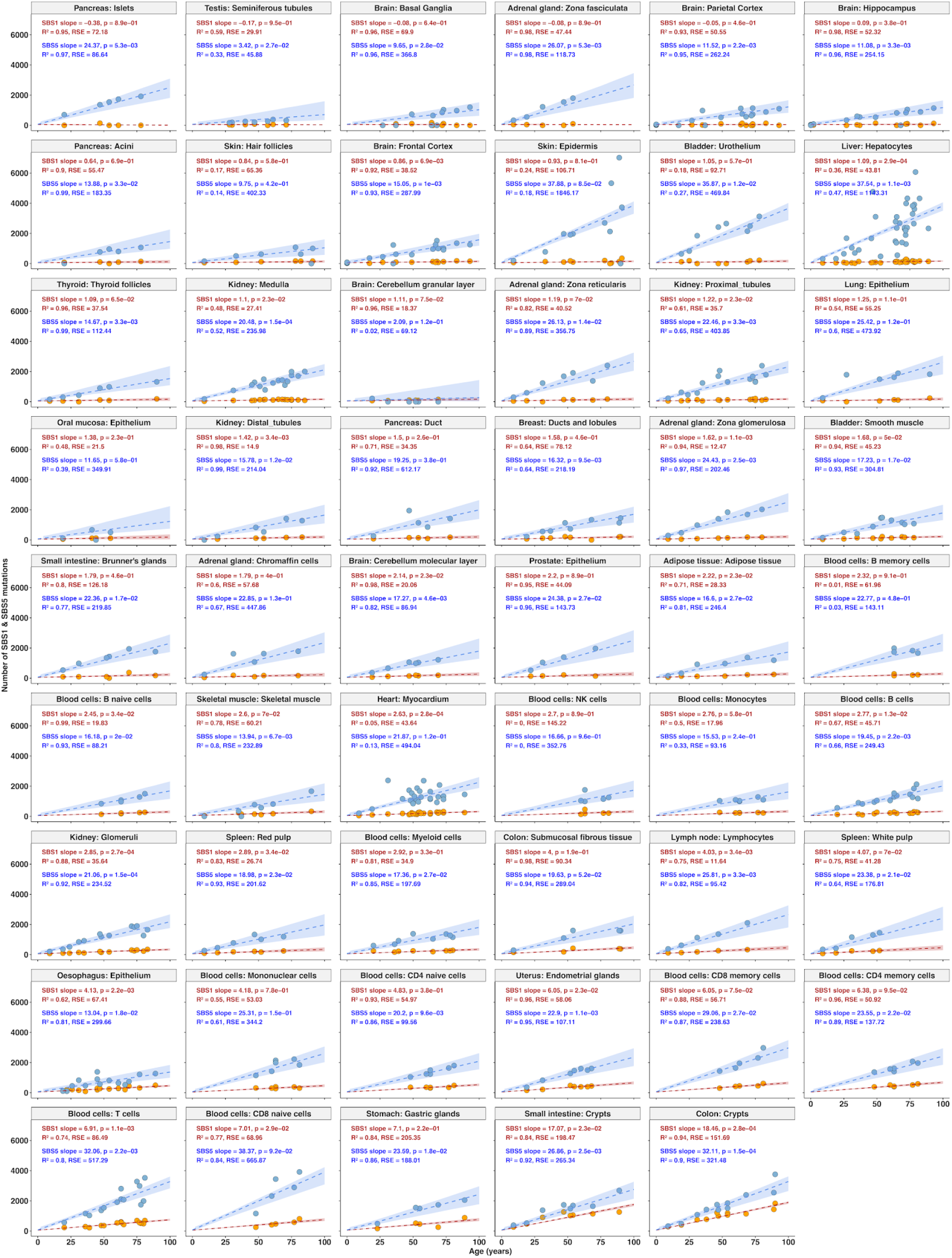
Comparison of SBS1- and SBS5-attributed mutation burdens with age across cell types. Number of mutations attributed to SBS1 (red) and SBS5 (blue) per diploid cell plotted against donor age (years) for each of the 53 cell types. Dashed lines, linear regression fits for SBS1 (red) and SBS5 (blue); shaded ribbons, 95% bootstrapped confidence intervals. Slope (mutations/diploid genome/year), R² and residual standard error (RSE) are shown for each signature and cell type (red text, SBS1; blue text, SBS5).

**Extended Figure 6.**
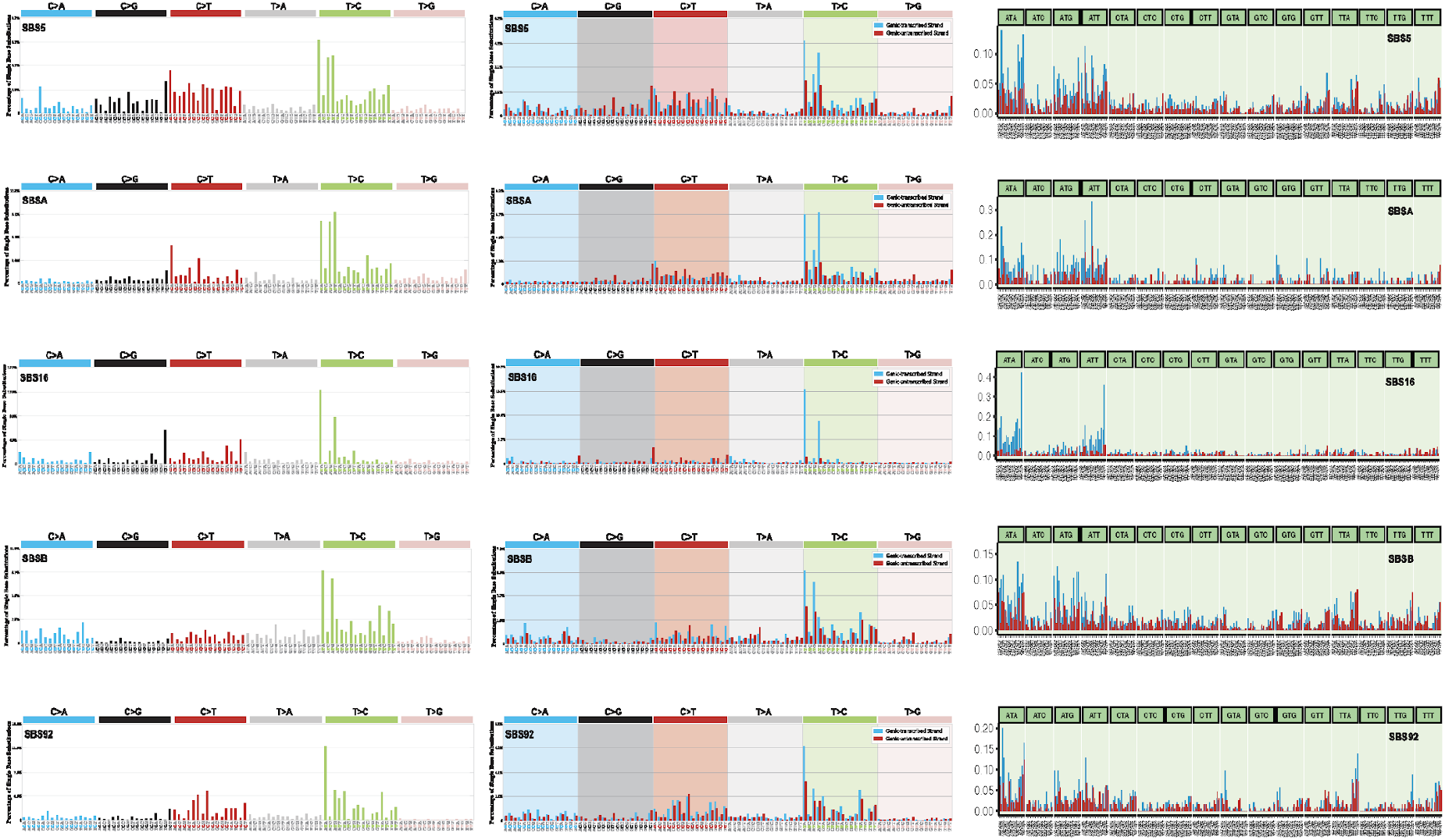
Comparison of mutational signatures characterised by T>C mutations at ApT dinucleotides. Mutational spectra of SBS5, SBSA, SBS16, SBSB and SBS92 (rows), which share a predominant pattern of T>C substitutions at ATN trinucleotides. Left column, percentage of single base substitutions in each of 96 trinucleotide contexts, grouped into the six substitution classes (C>A, C>G, C>T, T>A, T>C, T>G). Middle column, the same 96-class spectra stratified by transcriptional strand (blue, genic-transcribed strand; red, genic-untranscribed strand), showing strand bias characteristic of transcription-coupled repair. Right column, extended sequence-context spectra using 1,536 pentanucleotide classes (one base added on each side of the trinucleotide context), coloured by transcriptional strand as in the middle column, revealing additional sequence-context features that distinguish the five signatures.

