## Supplementary Notes for "A comprehensive atlas of somatic mutation rates and mutational signatures in normal human cells"

#### Note 1: LCM histology for each cell type

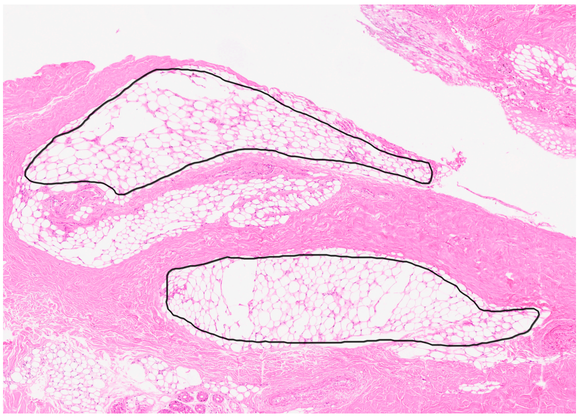

**Supplementary Figure 1.a**

**Adipose tissue**

Reference image of a 4 µm section from donor PD53199 at 4X magnification. Areas of the adipose tissue are outlined in black.

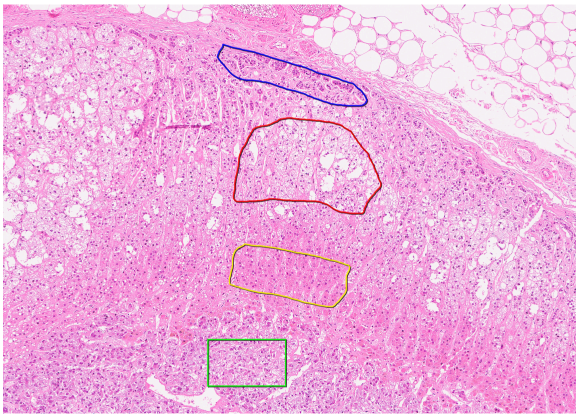

**Supplementary Figure 1.b**

**Adrenal gland**

Reference image of a 4 µm section from donor PD43851 at 6X magnification. Areas of the zona glomerulosa, zona fasciculata, zona reticularis and chromaffin cells are outlined in blue, red, yellow and green, respectively.

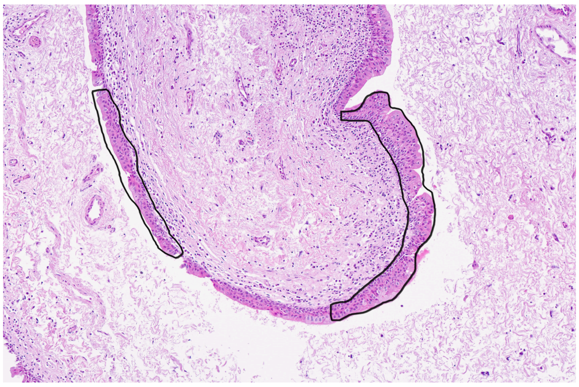

**Supplementary Figure 1.c**

**Bladder**

Reference image of a 4 µm section from donor PD41525 at 7X magnification. Areas of the urothelium are outlined in black.

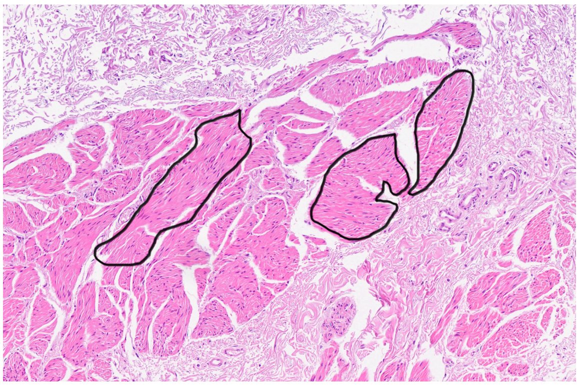

**Supplementary Figure 1.d**

**Bladder**

Reference image of a 4 µm section from donor PD41525 at 6X magnification. Areas of the smooth muscle are outlined in black.

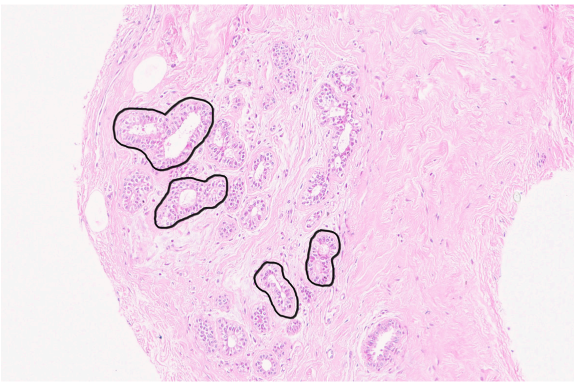

**Supplementary Figure 1.e**

**Breast**

Reference image of a 4 µm section from donor PD56386 at 10X magnification. Areas of the lobules and ducts are outlined in black.

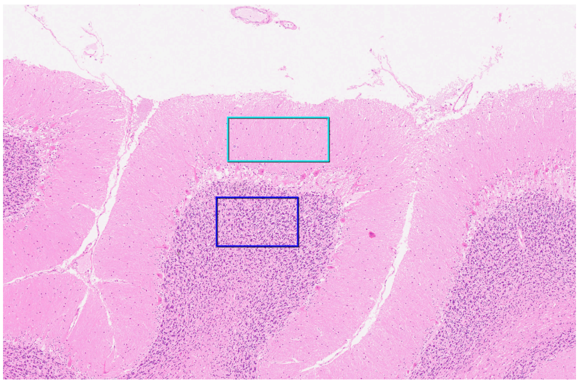

**Supplementary Figure 1.f**

**Cerebellum**

Reference image of a 4 µm section from donor PD53199 at 3X magnification. Areas of the molecular and granular layers are outlined in cyan-blue and blue, respectively.

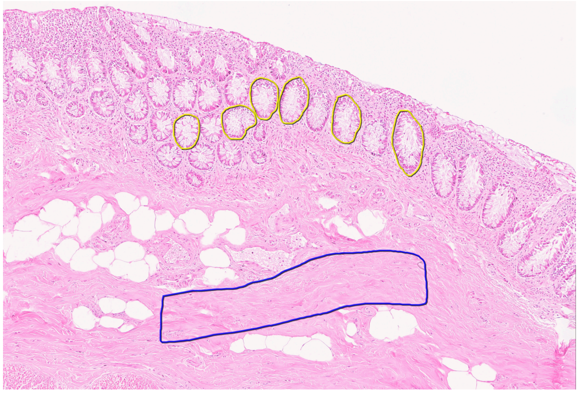

**Supplementary Figure 1.g**

**Colon**

Reference image of a 4 μm section from donor PD56385 at 6X magnification. Areas of the crypts and submucosal fibrous tissue are outlined in yellow and blue, respectively.

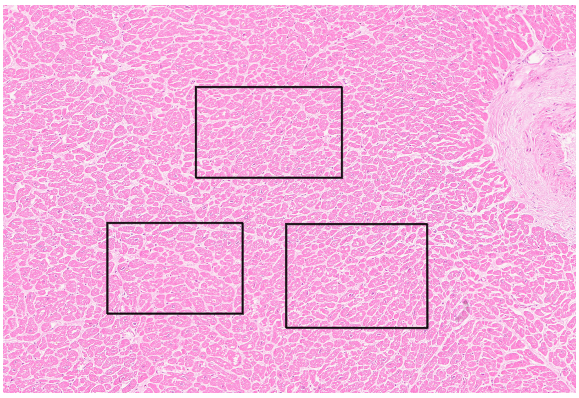

**Supplementary Figure 1.h**

**Heart**

Reference image of a 4 μm section from donor PD56385 at 6X magnification. Areas of the cardiac muscle are outlined in black.

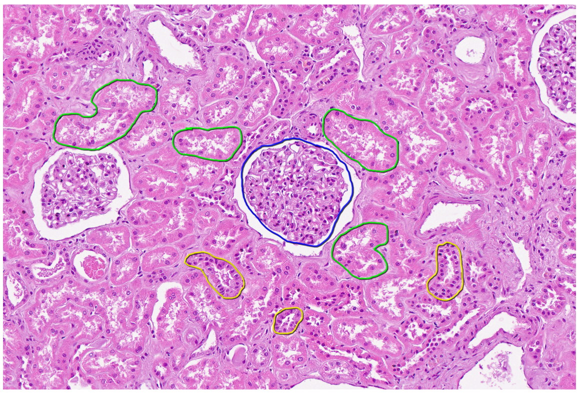

**Supplementary Figure 1.i**

**Kidney**

Reference image of a 4 μm section from donor PD41750 at 10X magnification. Areas of the glomeruli, as well as proximal and distal tubules, are outlined in blue, green, and yellow, respectively.

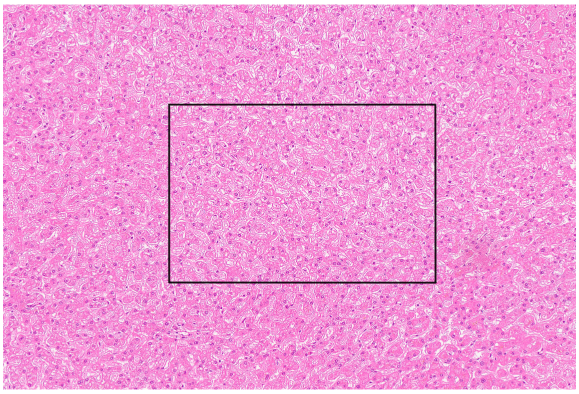

**Supplementary Figure 1.j**

**Liver**

Reference image of a 4 μm section from donor PD43851 at 6X magnification. Areas of the hepatocytes are outlined in black.

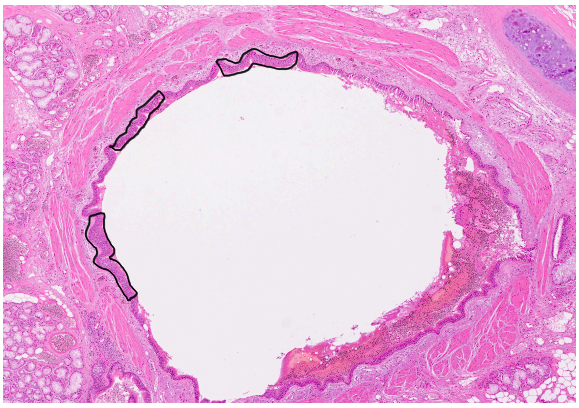

**Supplementary Figure 1.k**

**Lung**

Reference image of a 4 μm section from donor PD63074 at 2X magnification. Areas of the bronchial epithelium are outlined in black.

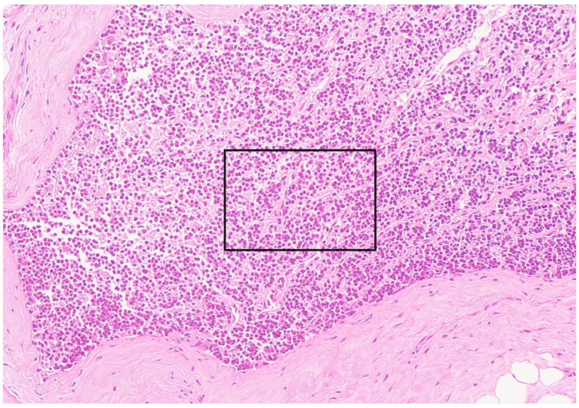

**Supplementary Figure 1.l**

**Lymph node**

Reference image of a 4 μm section from donor PD56384 at 13X magnification. Areas of the lymphocytes are outlined in black.

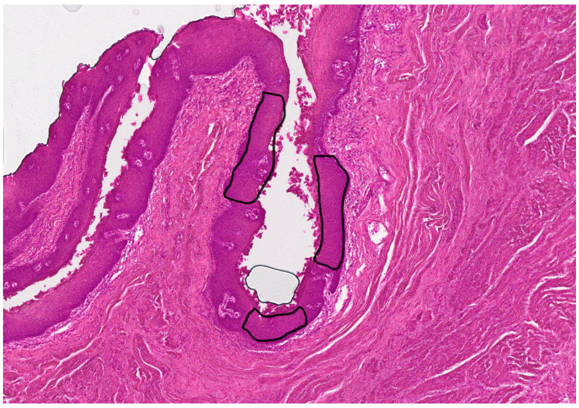

**Supplementary Figure 1.m**

**Oesophagus**

Reference image of a 16 μm section from donor PD38814 at 4X magnification. Areas of the epithelium are outlined in black.

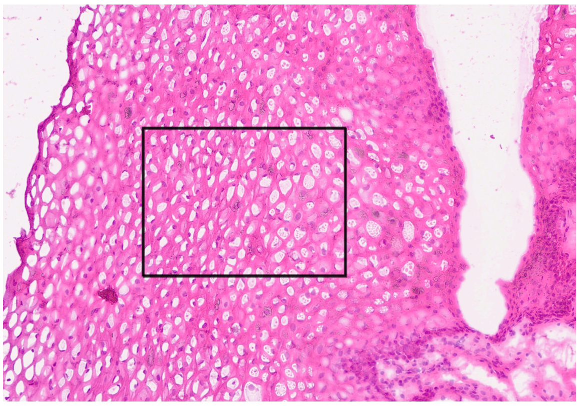

**Supplementary Figure 1.n**

**Oral mucosa**

Reference image of a 4 μm section from donor PD55073 at 6X magnification. Areas of the epithelium are outlined in black.

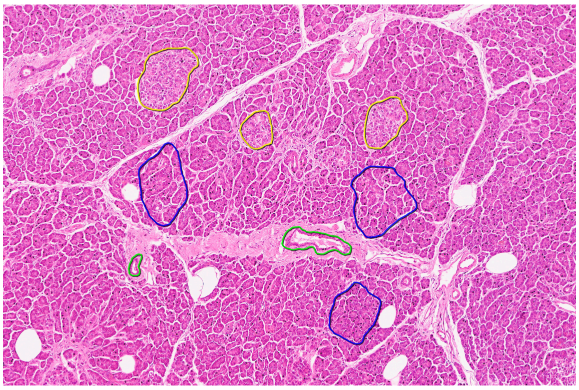

**Supplementary Figure 1.o**

**Pancreas**

Reference image of a 4 µm section from donor PD37726 at 6X magnification. Areas of the acinus, islets, and ducts are outlined in blue, yellow and green, respectively.

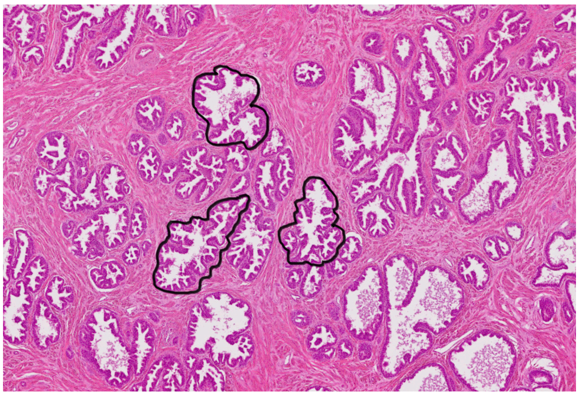

**Supplementary Figure 1.p**

**Prostate**

Reference image of a 16 µm section from donor PD43390 at 2X magnification. Areas of the epithelium are outlined in black.

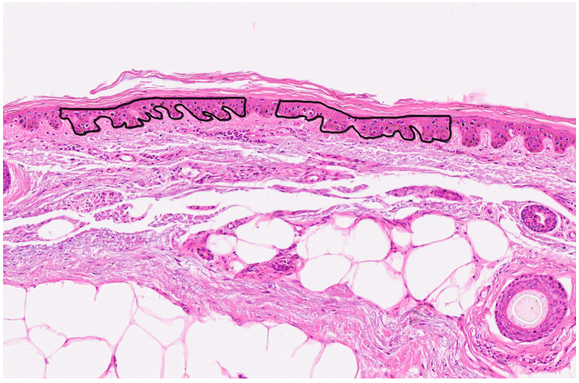

**Supplementary Figure 1.q**

**Skin**

Reference image of a 4 µm section from donor PD38814 at 8X magnification. Areas of the epidermis are outlined in black.

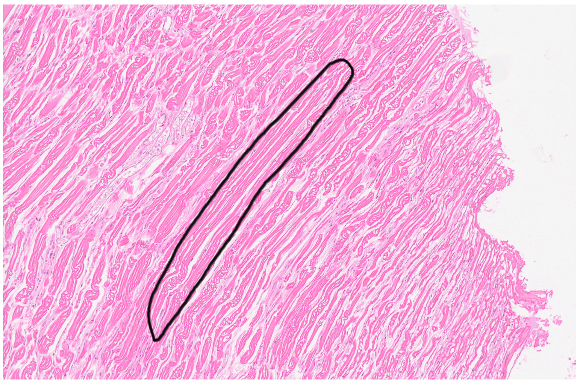

**Supplementary Figure 1.r**

**Skeletal muscle**

Reference image of a 4 µm section from donor PD52199 at 5X magnification. Areas of the skeletal muscle are outlined in black.

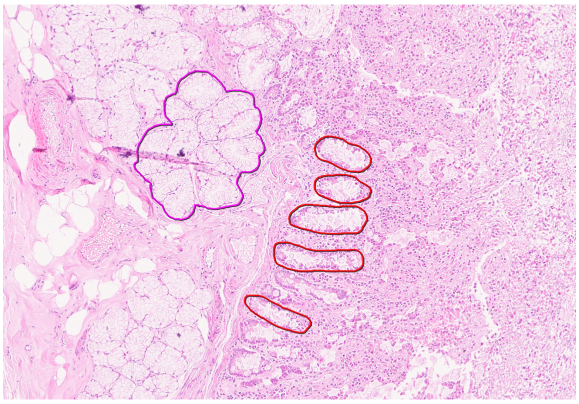

**Supplementary Figure 1.s**

**Small intestine**

Reference image of a 4 µm section from donor PD53686 at 7X magnification. Areas of the crypts and Brunner's glands are outlined in red and purple-pink, respectively.

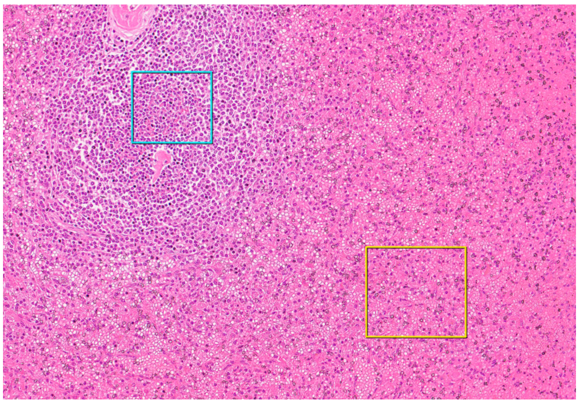

**Supplementary Figure 1.t**

**Spleen**

Reference image of a 4 µm section from donor PD43851 at 10X magnification. Areas of the white and red pulps are outlined in cyan-blue and yellow, respectively.

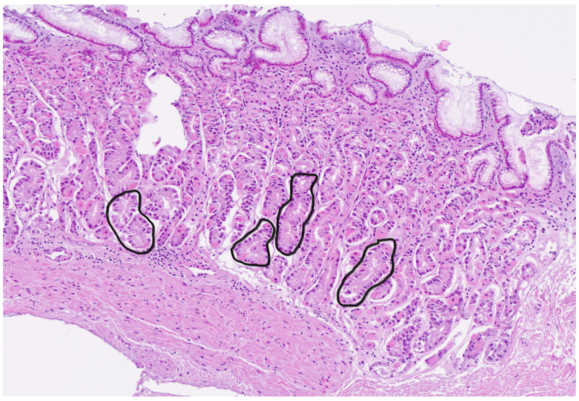

**Supplementary Figure 1.u**

**Stomach**

Reference image of a 4 µm section from donor PD45518 at 7X magnification. Areas of the gastric glands are outlined in black.

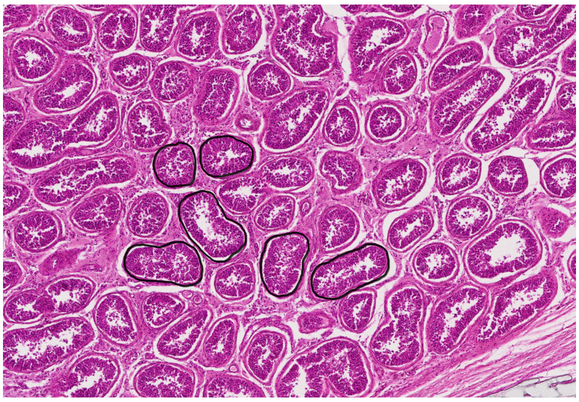

**Supplementary Figure 1.v**

**Testis**

Reference image of a 16 µm section from donor PD53625 at 4X magnification. Areas of the seminiferous tubules are outlined in black.

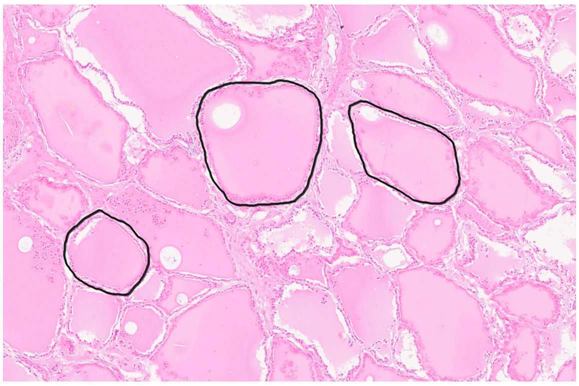

**Supplementary Figure 1.w**

**Thyroid**

Reference image of a 4 μm section from donor PD56385 at 7X magnification. Areas of the follicles are outlined in black.

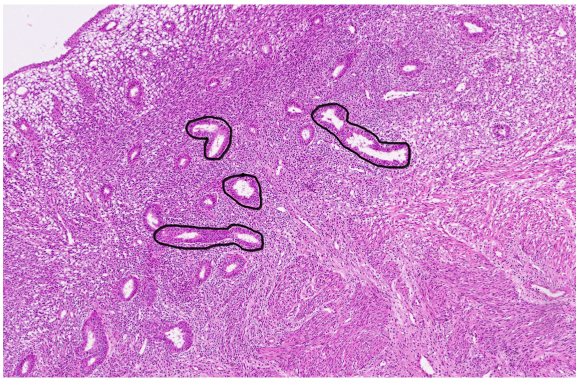

**Supplementary Figure 1.x**

**Uterus**

Reference image of a 4 μm section from donor PD60371 at 4X magnification. Areas of the endometrial glands are outlined in black.

### Note 2: Regression analysis

##### A. Whole dataset evaluation

**1. Rationale for model selection**

To robustly investigate the trend of somatic mutation accumulation in normal human cells, we evaluated three mathematical relationships between donor age (*age*) and mutation burden (*burden_wg)*: linear, quadratic, and exponential using *lme4*[1] and *nlme*[2] packages in R.

As an initial test across the whole dataset, we used mixed-effects models with donor *(individual)* as a random effect to account for the non-independence of multiple samples from the same donor.

We applied the delta Akaike Information Criterion (ΔAIC) to validate cross-model comparisons.

**2. Models for testing**

<u>Model A: Linear Mixed-Effects Model</u>

The linear model assumes a constant mutation accumulation rate with age, allowing the baseline burden (intercept) and accumulation rate (slope) to vary across cell types while accounting for individual effects on the intercept.

mm_linear = lmer(burden_wg ~ age + (1 | individual))

<u>Model B: Quadratic Mixed-Effects Model</u>

The quadratic model introduces a second-order polynomial term to capture potential acceleration or deceleration of mutation rates in later life.

mm_quadratic = lmer(burden_wg ~ age + I(age^2) + (1 | individual))

<u>Model C: Exponential Non-Linear Mixed-Effects Model</u>

The exponential model represents a non-linear acceleration of somatic mutations, estimated via a non-linear mixed-effects (nlme) framework.

mm_exp = nlme(burden_wg ~ a * exp(b * age), data = burden_model, fixed = a + b ~ 1, random = a ~ 1 | individual, start = c(a = a_start, b = b_start))

##### 3. Models comparison

AIC is measured by balancing goodness-of-fit and the number of variables included, computed using the log-likelihood and default parameters from the AIC() function in the *stats* package in R[3]. Smaller AIC scores indicate better model fit. To evaluate how well the models perform compared to the best one, we calculated ΔAIC - the difference between a model's AIC score and the minimum AIC score.

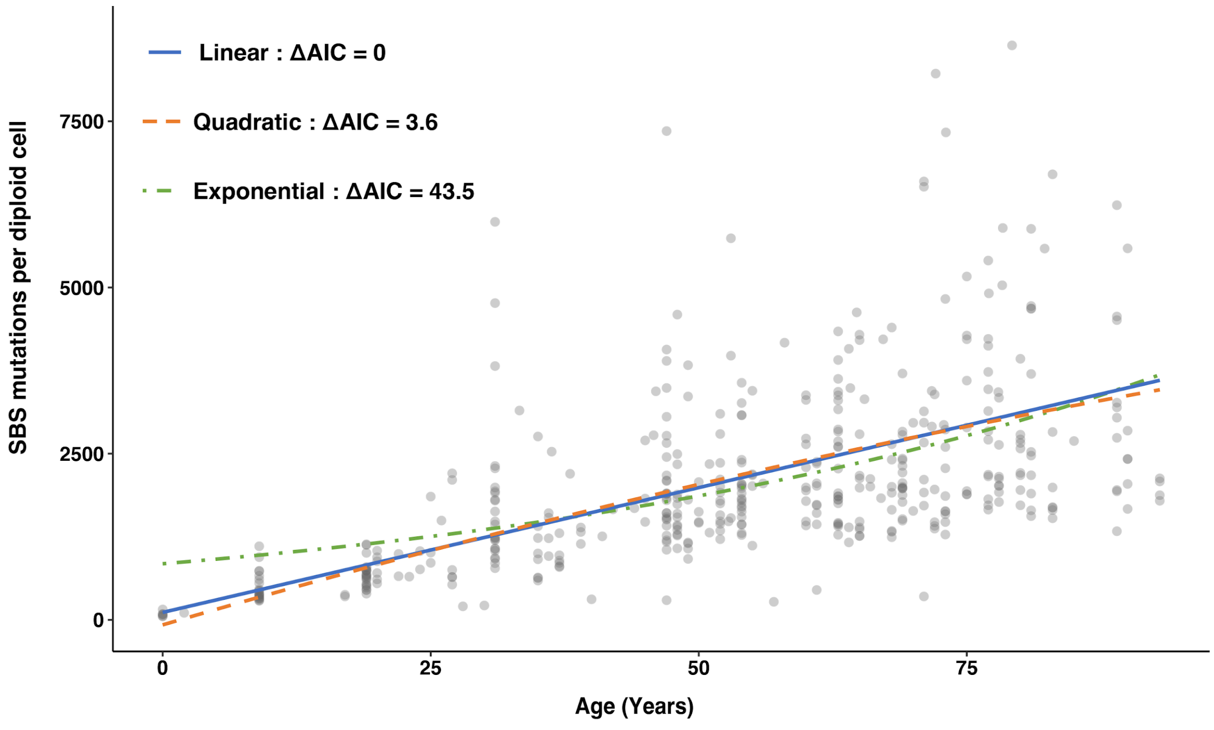


**Supplementary Figure 2. Comparison of linear, quadratic, and exponential model fits of somatic mutation burden with age.** Scatter plot of single-base substitution (SBS) mutation burden per diploid cell against donor age across all sequenced samples, overlaid with the fitted linear (blue, solid), quadratic (orange, dashed) and exponential (green, dot-dashed) models. Each grey point represents one sample. ΔAIC relative to the best-fitting model is shown for each model in the legend.

Furthermore, we decisively rejected the exponential model due to its high ΔAIC (43.5), indicating that an exponential runaway process cannot explain mutation accumulation with age across cell types.

In conclusion, we selected the linear mixed-effects model for downstream manuscript analyses, establishing that somatic mutation burden scales at a steady, linear rate across the human lifespan.

#### B. Cell-type specific mutation rates

##### 1. Rationale for model selection

Having established a linear relationship between donor age and mutation burden across the dataset, we next estimated mutation rates separately for each cell type. We compared four linear mixed-effects models that differ in how random effects are specified for cell type and donor, to identify the random-effects structure that best captures both between-cell-type variation in mutation rate and repeated sampling from the same donor, without overfitting.

All models were fitted by Maximum Likelihood (REML = FALSE) using the *bobyqa* optimiser [1] to ensure convergence and enable valid likelihood-ratio comparisons across nested models.

##### 2. Models tested

<u>Model 1: Random slope for age by cell type only, no donor effect.</u>

model1 = lmer(burden_wg ~ age + (0 + age | cell type), data = burden_model, REML = F, control = lmerControl(optimizer = "bobyqa"))

<u>Model 2: Random slope for age by cell type, plus a random intercept for donor.</u>

model2 = lmer(burden_wg ~ age + (0 + age | cell type) + (1 | individual), data = burden_model, REML = F, control = lmerControl(optimizer = "bobyqa"))

Model 3: Random intercept and slope for cell type (correlated), no donor effect.

model3 = lmer(burden_wg ~ age + (1 + age | cell type), data = burden_model, REML = F, control = lmerControl(optimizer = "bobyqa"))

Model 4: Random intercept and slope for cell type, plus a random intercept for donor.

model4 <- lmer(burden_wg ~ age + (1 + age | cell type) + (1 |individual), data = burden_model, REML = F, control = lmerControl(optimizer = "bobyqa"))

**3. Model comparison**

Nested models were compared using a likelihood-ratio test (*anova()* in *the stats* package in R[4]).

| Comparison | AIC | $\chi^2$ | df | p-value | Interpretation |
|---|---|---|---|---|---|
| Model 1 vs Model 2 | 7095.0 / 7079.2 | 17.8 | 1 | $2.47 \times 10^{-5}$ | Adding a donor random intercept significantly improves fit |
| Model 2 vs Model 3 | 7079.2 / 7097.7 | 0.00 | 1 | 1.00 | No improvement from a correlated intercept-slope structure for cell type |
| Model 3 vs Model 4 | 7097.7 / 7082.0 | 17.7 | 1 | $2.57 \times 10^{-5}$ | Adding a donor random intercept significantly improves fit |

**Supplementary Table A. Model comparison based on *anova()* results**.

Adding a random intercept for donor *(individual)* significantly improved model fit in both relevant comparisons (Model 1 vs 2; Model 3 vs 4), confirming that accounting for repeated sampling within donors is necessary. By contrast, allowing the cell-type random effect to include a correlated intercept term (Model 3) did not improve fit over the simpler slope-only structure (Model 2).

We additionally assessed model degeneracy using *isSingular()*. Model 2 converged to a non-singular fit *(isSingular = FALSE)*, whereas Model 4, despite its lower AIC, converged to a singular fit *(isSingular = TRUE)*, indicating an over-parameterised random-effects structure not properly supported by the data.

Therefore, we selected Model 2 for downstream analyses, as it provided the best-supported, non-degenerate random-effects structure: allowing mutation rate (slope) to vary by cell type while accounting for non-independence of repeated measurements from the same donor.

### Note 3: Mutational signatures analysis

#### A. SBS mutational signatures

To characterise the mutational processes operating across normal human tissues, we performed *de novo* mutational signature extraction separately for single-base substitutions (SBS) and small insertions/deletions (indels, ID) using three independent computational approaches: a hierarchical Dirichlet process model (HDP)[5], SigProfilerExtractor (SPE)[6] and MuSiCal[7]. We cross-compared signatures identified by each method using cosine similarity to assess reproducibility, and matched them against the COSMIC v3.4 [8] reference signature catalogue. Consensus *de novo* signatures (derived from HDP, the primary reference framework) were then decomposed into combinations of COSMIC reference signatures where possible, and the exposure of each signature was attributed back to individual samples. Finally, the robustness of signature attribution was assessed by bootstrap resampling.

##### 1. Mutational signatures extracted from HDP

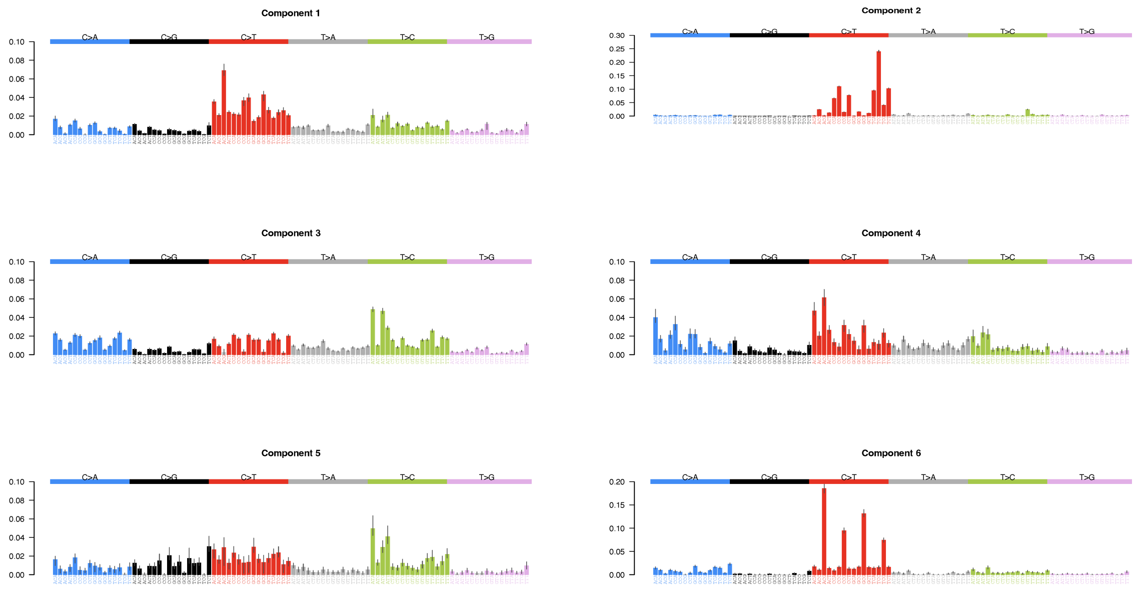

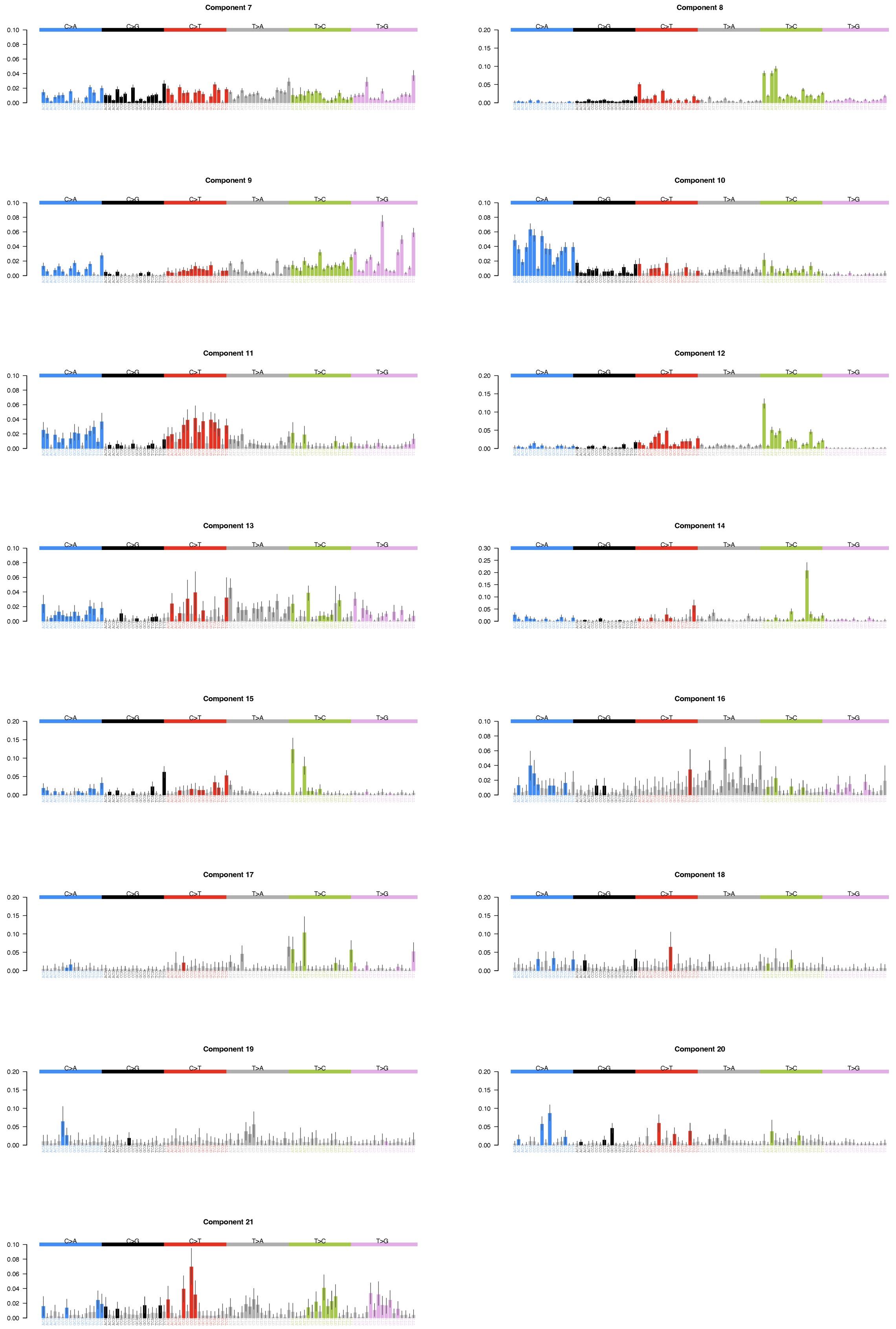
Component 7
Component 8
Component 9
Component 10
Component 11
Component 12
Component 13
Component 14
Component 15
Component 16
Component 17
Component 18
Component 19
Component 20
Component 21
C>A
C>G
C>T
T>A
T>C
T>G

**Supplementary Figure 3. Mutation spectra of 21 *de novo* SBS signatures from HDP.** The six substitution types are labelled across the top. Within each panel, the contributions from the 96-trinucleotide contexts (bases immediately 5′ and 3′ of the mutated base) are shown.

#### 2. Mutational signatures extracted from SPE

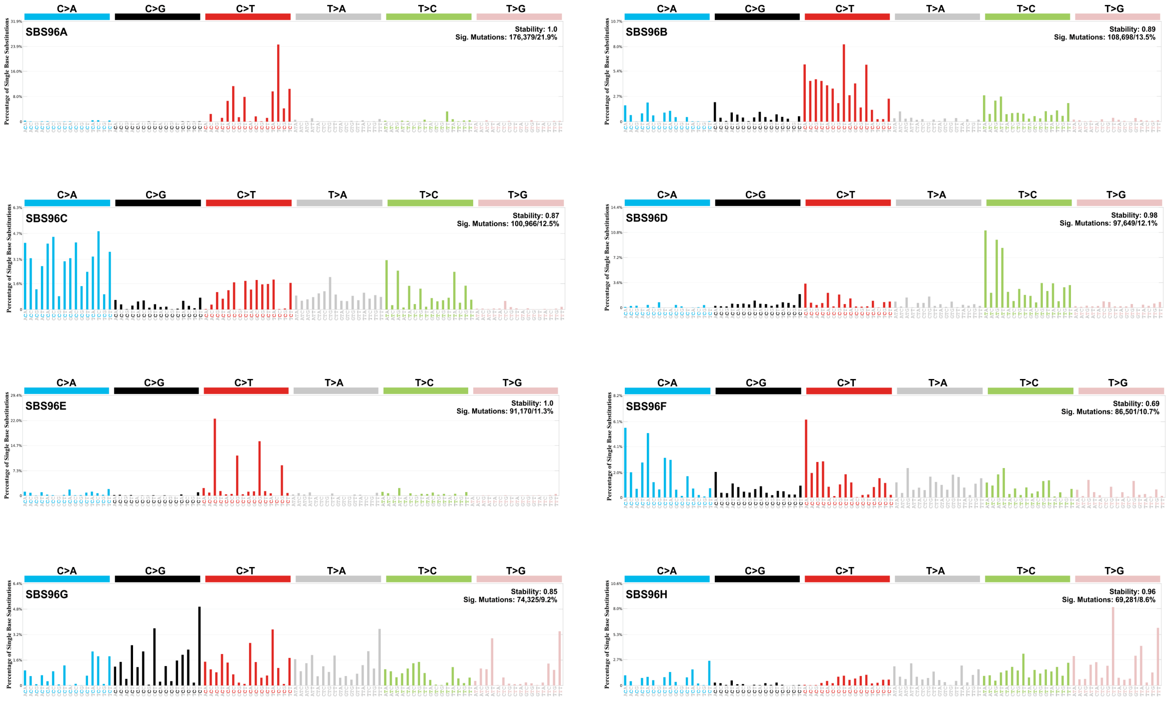


**Supplementary Figure 4. Mutation spectra of eight *de novo* SBS signatures from SPE.** The six substitution types are labelled across the top. Within each panel, the contributions from the 96-trinucleotide contexts (bases immediately 5′ and 3′ of the mutated base) are shown.

#### 3. Mutational signatures extracted from MuSiCal

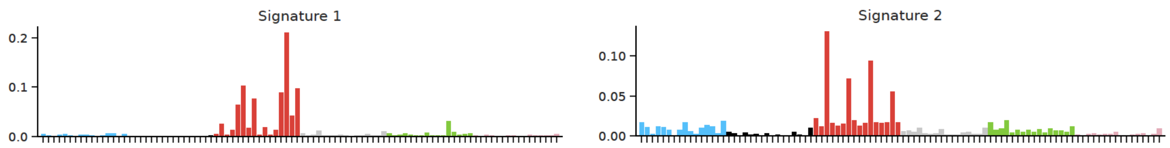

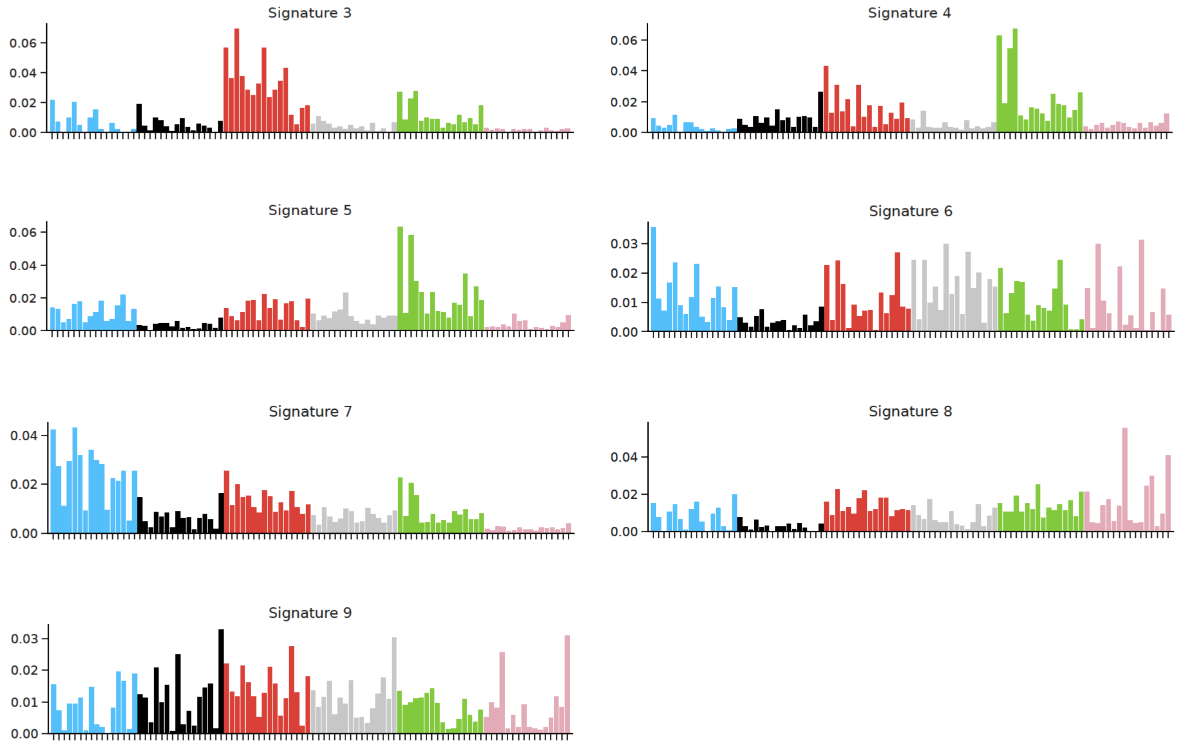


**Supplementary Figure 5. Mutation spectra of nine *de novo* SBS signatures from MuSiCal.** The six substitution types are labelled across the top. Within each panel, the contributions from the 96-trinucleotide contexts (bases immediately 5′ and 3′ of the mutated base) are shown.

#### 4. Mutational signatures comparison

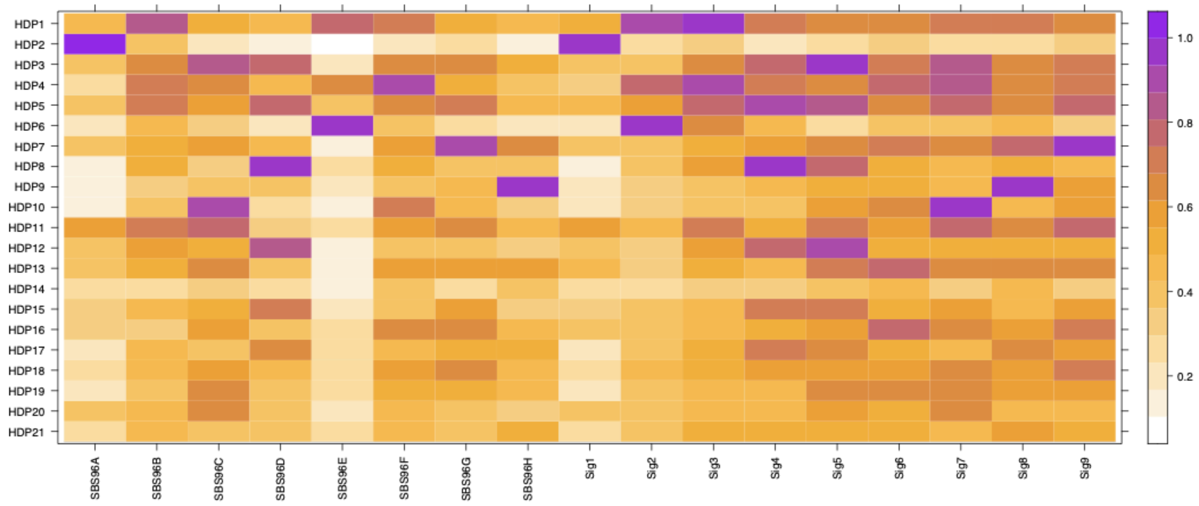

**Supplementary Figure 6. Cosine similarities between HDP96 signatures and signatures from the other two methods.** The x-axis shows 21 *de novo* signatures extracted from HDP, the y-axis shows eight *de novo* signatures from SPE, and nine from MuSiCal. The colour gradient presents cosine similarity scores. The highest similarity is shown in purple, while the least similar are in white.

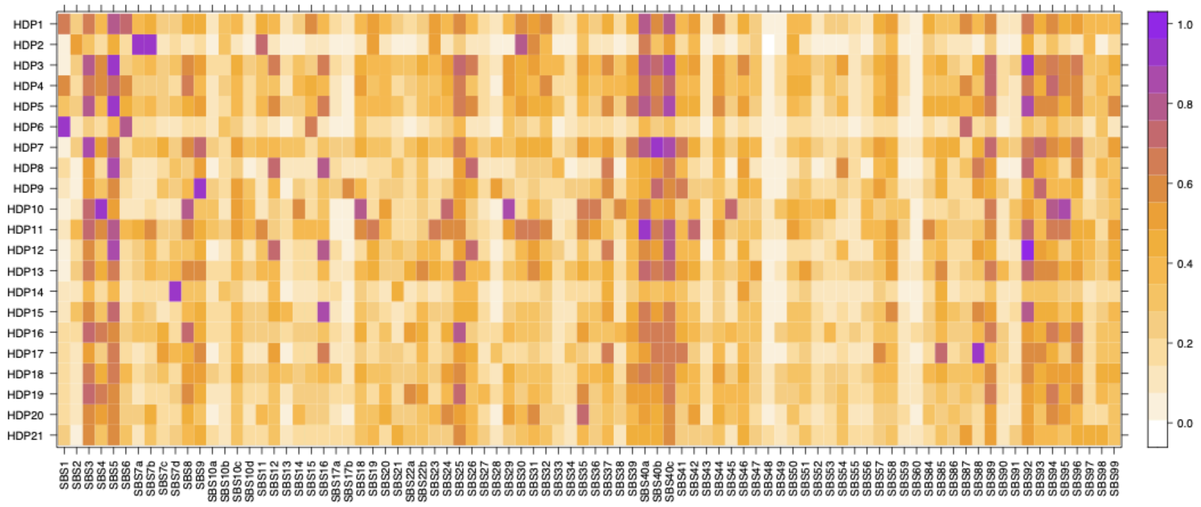


**Supplementary Figure 7. Cosine similarities between HDP96 signatures and COSMIC reference signatures.** The x-axis shows 21 *de novo* signatures extracted from HDP, and the y-axis shows a list of COSMIC reference signatures v3.4 [8]**.** The colour gradient presents cosine similarity scores. The highest similarity is shown in purple, while the least similar are in white.

| HDP | SPE | MuSiCal | Score | COSMIC | Concl. |
|---|---|---|---|---|---|
| Component 1 | | Sig3 (0.94) | 2 | | Y |
| Component 2 | SBS96A (0.99) | Sig1 (0.99) | 3 | SBS7a (0.93), SBS7b (0.92) | Y |
| Component 3 | | Sig5 (0.97) | 2 | SBS5 (0.92) | Y |
| Component 4 | SBS96F (0.9) | | 2 | | Y |

| | | | | | |
|---|---|---|---|---|---|
| Component 5 | | Sig4 (0.91) | 2 | SBS5 (0.93) | Y |
| Component 6 | SBS96E (0.99) | Sig2 (0.98) | 3 | SBS1 (0.96) | Y |
| Component 7 | SBS96G (0.93) | Sig9 (0.97) | 3 | SBS40b (0.95) | Y |
| Component 8 | SBS96D (0.97) | Sig4 (0.96) | 3 | | Y |
| Component 9 | SBS96H (0.98) | Sig8 (0.95) | 3 | SBS9 (0.96) | Y |
| Component 10 | SBS96C (0.93) | Sig7 (0.94) | 3 | SBS4 (0.95) | Y |
| Component 11 | | | | SBS40a (0.93) | Y |
| Component 12 | | | | SBS92 (0.96) | Y |
| Component 13 | | | | | N |
| Component 14 | | | | SBS7d (0.9) | Y |
| Component 15 | | | | SBS16 (0.9) | Y |
| Component 16 | | | | | N |
| Component 17 | | | | SBS88 (0.91) | Y |
| Component 18 | | | | | N |
| Component 19 | | | | | N |
| Component 20 | | | | | N |
| Component 21 | | | | | N |

**Supplementary Table B. Cross-method comparison and consensus assignment of de novo SBS signatures.** For each of the 21 de novo signatures extracted by HDP, the best-matching *de novo* signature independently extracted by SPE and MuSiCal is shown, together with the best-matching COSMIC v3.4 reference signature [8] *(cosine similarity).* Score indicates the number of methods (maximum of three) in which a corresponding *de novo* signature was independently extracted. Concl. indicates whether the HDP-derived signature was retained (Y) or excluded (N) from the final consensus signature score greater than or equal to 2 and/or a confident match to a COSMIC reference signature.

#### 5. Unclassified signatures

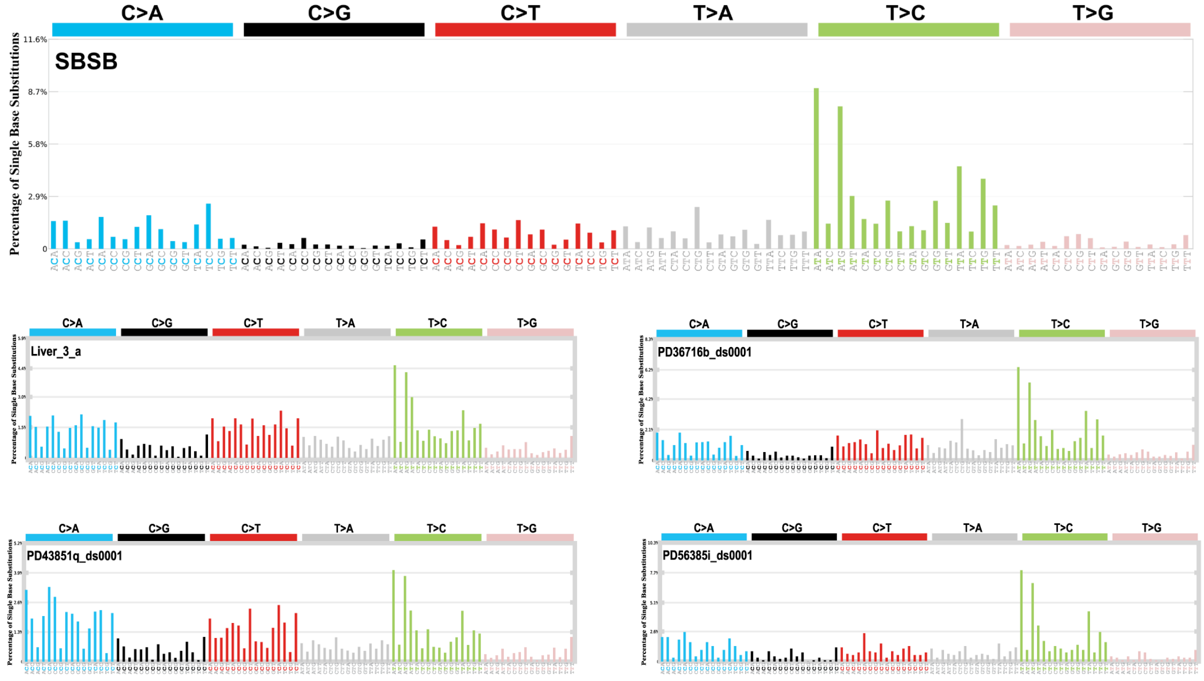


**Supplementary Figure 8. Mutation spectra of the SBSB signature extracted from liver samples.** SBS96 mutational profiles for four representative liver samples are shown alongside the extracted signature for comparison. The six substitution types are indicated above each panel. Within each panel, bars represent the relative contribution of each of the 96 trinucleotide contexts, defined by the bases immediately 5′ and 3′ of the mutated base.

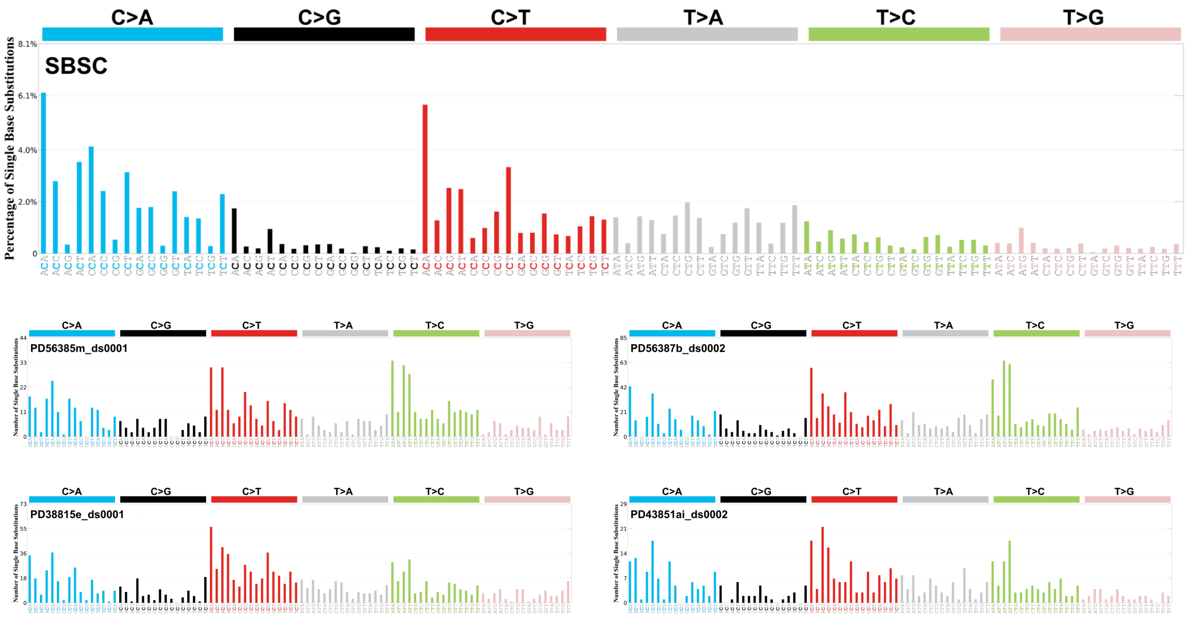


**Supplementary Figure 9. Mutation spectra of the SBSB signature extracted from brain, heart and skeletal muscle samples.** SBS96 mutational profiles for representative brain (PD56385m_ds0001 and PD56387b_ds0002), heart (PD38815e_ds0001), and skeletal muscle (PD43851ai_ds0002) samples are shown alongside the extracted signature for comparison. The six substitution types are indicated above each panel. Within each panel, bars represent the relative contribution of each of the 96 trinucleotide contexts, defined by the bases immediately 5′ and 3′ of the mutated base.

#### 6. Mutational signatures decomposition

| HDP mutational signatures | Decomposition |
| --- | --- |
| 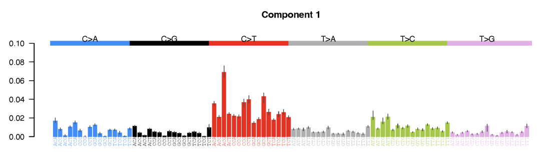 | SBS1 (17.94%) & SBS5 (70.98%) & SBS19 (11.08%) |
| 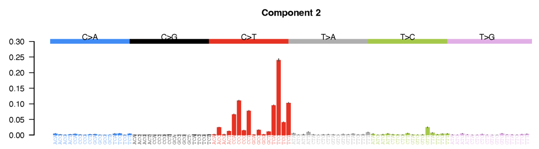 | SBS7a (42.68%) & SBS7b (49.98%) & SBS7d (7.34%) |

| | |
|---|---|
| 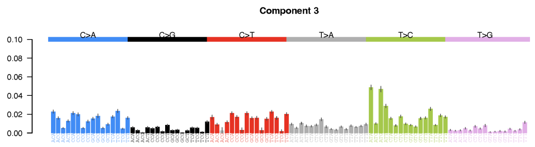  | SBS4 (17.74%) & SBS5 ( 50.12%) & SBSB (32.14%) |
| 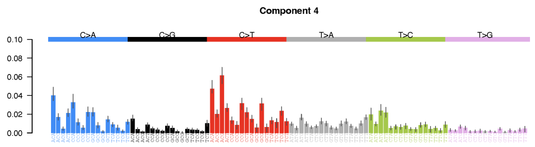  | SBS1 (10.74%) & SBS5 (32.04%) & SBSC (57.22%) |
| 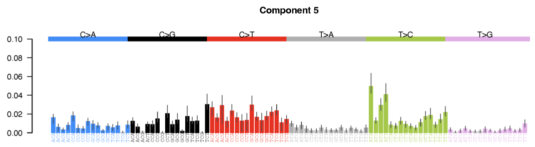  | SBS1 (3.46%) & SBS5 (96.54%) |
| 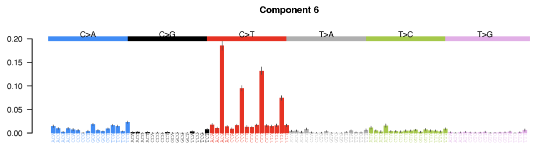  | SBS1 (51.04%) & SBS5 (32.02%) & SBS18 (16.94%) |
| 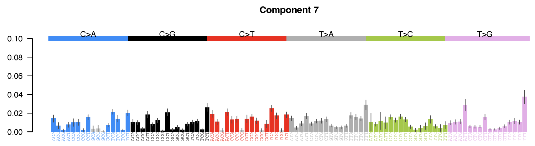  | SBS40b (69.52%) & SBS40c (30.48%) |
| 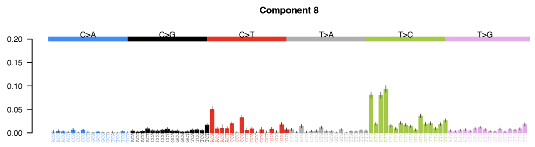  | SBSA |
| 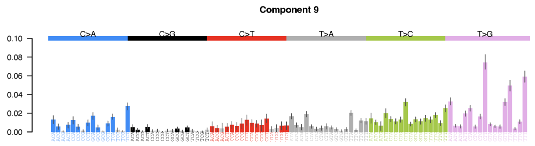  | SBS1 (0.54%) & SBS5 (7.60%) & SBS9 (85.80%) & SBS17b (6.06%) |
| 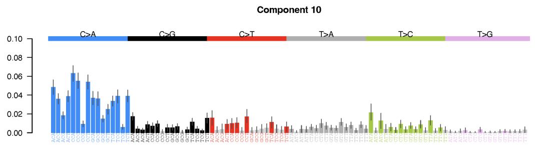  | SBS1 (0.02%) & SBS4 (78.16%) & SBS5 (21.82%) |

| | |
|---|---|
| 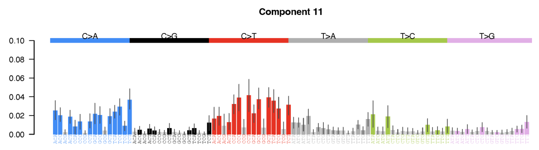 | SBS5 (3.56%) & SBS40a (96.44%) |
| 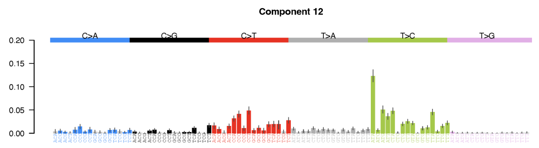 | SBS92 (100.00%) |
| 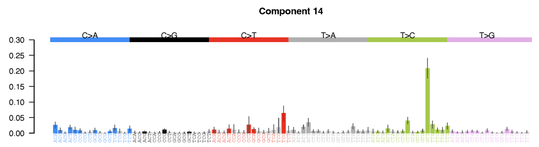 | SBS1 (1.84%) & SBS7d (82.34%) & SBS10b (15.82%) |
| 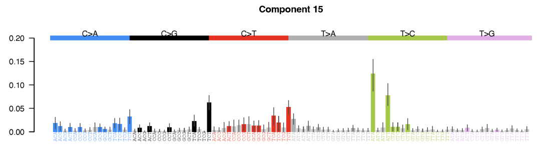 | SBS1 (1.96%) & SBS2 (7.28%) & SBS5 (29.02%) & SBS13 (11.12%) & SBS16 (50.62%) |
| 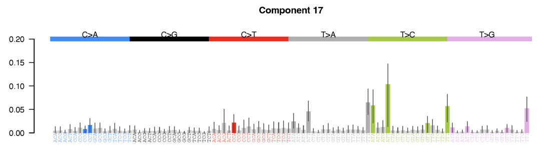 | SBS1 (2.46%) & SBS5 (40.54%) & SBS88 (57.00%) |

**Supplementary Table C. Decomposition of consensus HDP mutational signatures into COSMIC reference signatures.** Each signature was decomposed into a combination of COSMIC v3.4 reference [8] signatures using SigProfilerAssignment. Blank cells indicate signatures for which no decomposition met the reconstruction accuracy threshold (cosine similarity greater than or equal to 0.9) and which were therefore retained as novel, undecomposed signatures.

#### 7. Mutational signatures attribution from HDP

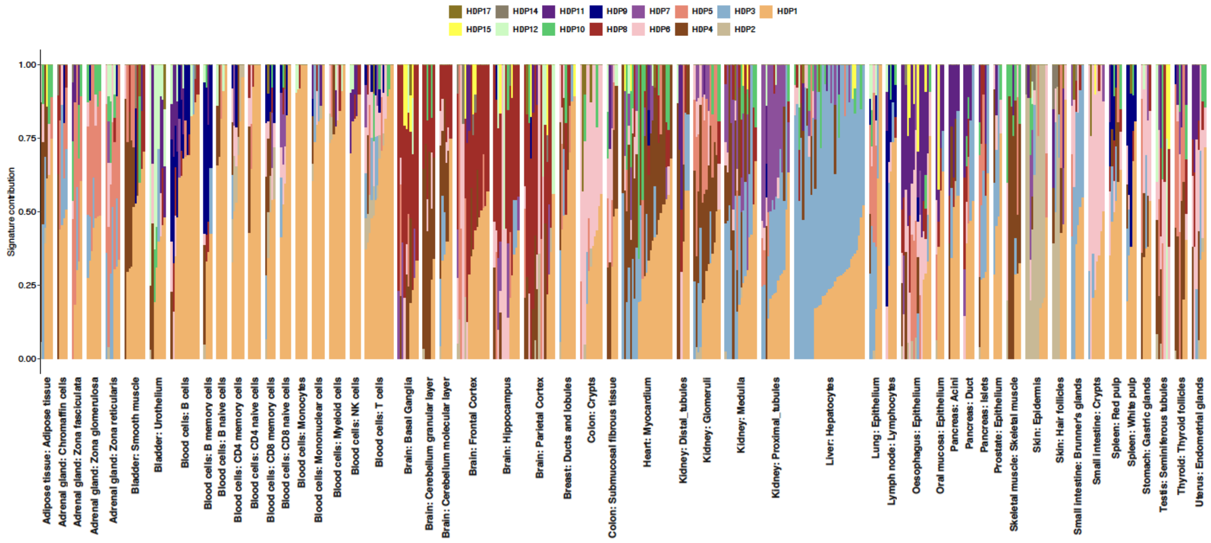


**Supplementary Figure 10. Contribution of *de novo* SBS signatures to mutation catalogues of each sample.** The 15 *de novo* SBS signatures extracted from HDP96 were attributed to each sample and grouped by tissue structure in alphabetical order. Each colour represents one signature.

#### 8. Validation of SigProfilerAssignment with bootstrapping

To evaluate the accuracy of signature attribution, we generated simulated mutational catalogues of 1000 samples. We first drew a set of signature proportions from a symmetric Dirichlet distribution (with different weights for SBS1, SBS5, and SBS19) for each sample, then scaled them by each sample's total mutation burden (up to 3,000 mutations). To incorporate realistic count-level noise, mutation counts for each sample and mutation type were then drawn from a Poisson distribution with mean equal to the corresponding expected count, yielding the final simulated mutational catalogue. We then re-attributed the simulated data with a set of COSMIC signatures (SBS1, SBS5, SBS16, SBS18, SBS19, SBS40a, SBS40b, and SBS40c)[8] using SigProfilerAssignment, and assessed confidence in the fitted exposures with 1,000 bootstrap resamplings of each mutational catalogue. For each signature, we compared the number of attributed mutations across four conditions: the ground-truth (simulated) exposure, the default fit (observed), and the bootstrap-derived exposures before (bs_unpruned) and after (bs_pruned) removal of low-confidence signature calls. Close agreement between the ground-truth and bs_pruned exposures indicated that signature attribution was robust to resampling, and that pruning removed spurious low-level calls without biasing estimates of the dominant signatures. We then used these confident signatures to re-attribute each cell type.

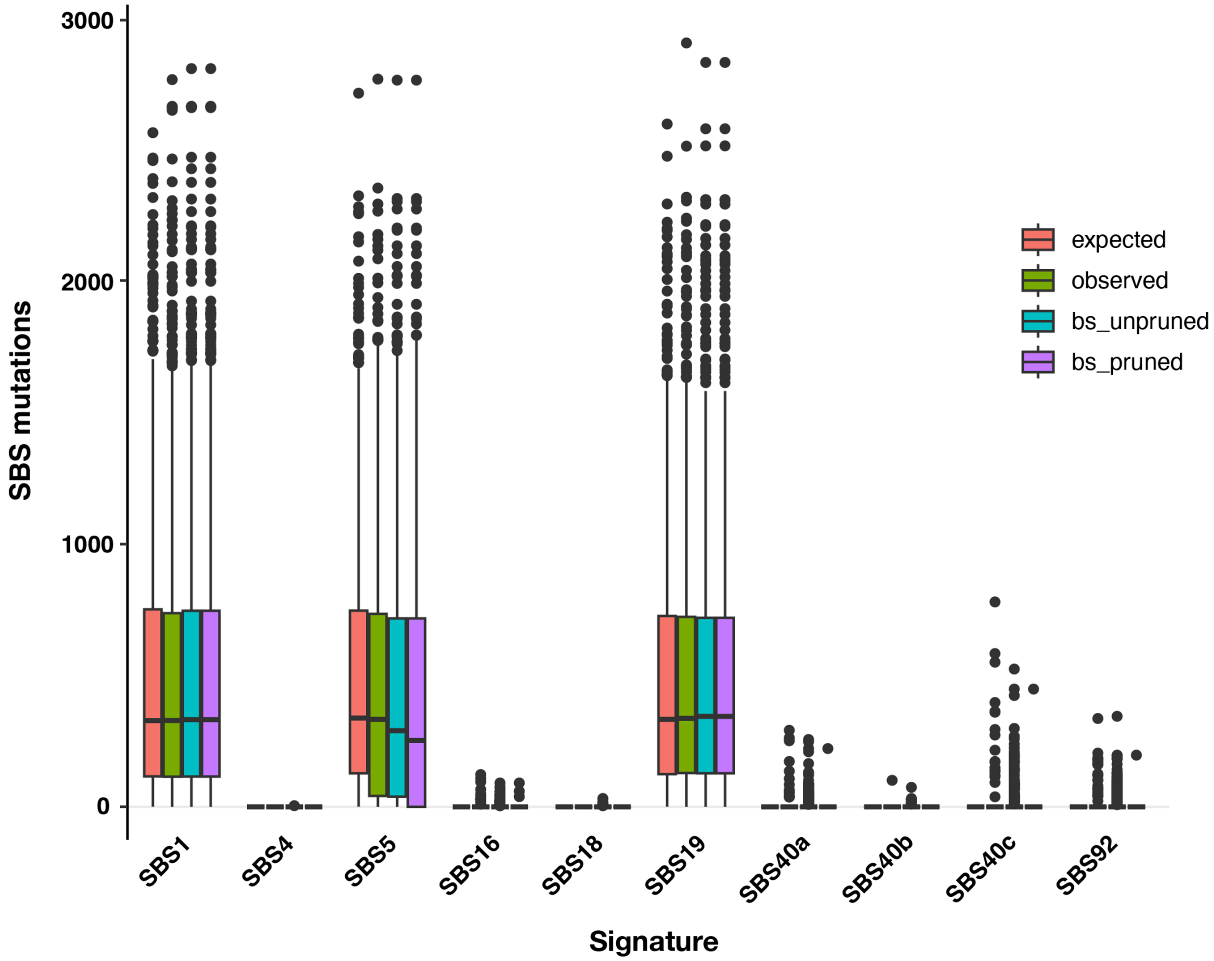


**Supplementary Figure 11. Bootstrap validation of signature attribution by SigProfilerAssignment.** The y-axis shows the number of SBS mutations in each sample, and the x-axis shows the attributed signatures. Centre lines show the median, box limits the interquartile range (IQR), whiskers 1.5× IQR, and points show values beyond this range.

#### B. INDEL mutational signatures

As for SBS signatures above, *de novo* INDEL (ID) signature extraction using HDP, SPE and MuSiCal, respectively.

##### 1. Mutational signatures extracted from HDP

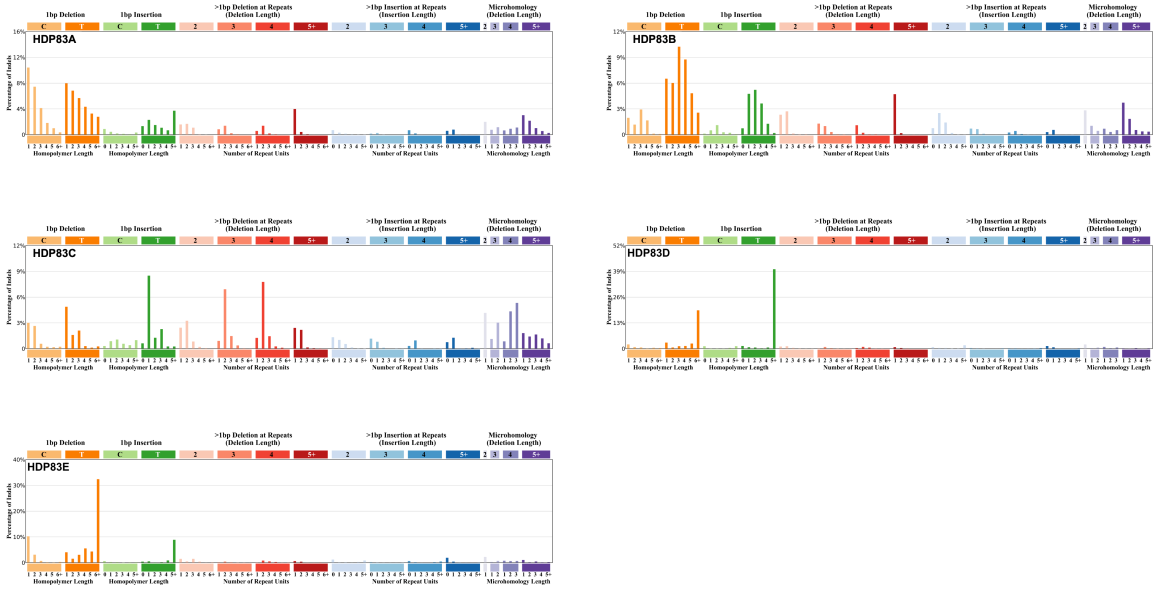


**Supplementary Figure 12. Mutation spectra of five *de novo* ID signatures from HDP.** The coloured bars at the top represent the size of the insertions or deletions. Within each panel, the x-axis depicts the count of repeat units in the reference genome corresponding to the inserted or deleted sequence.

##### 2. Mutational signatures extracted from SigProfilerExtractor

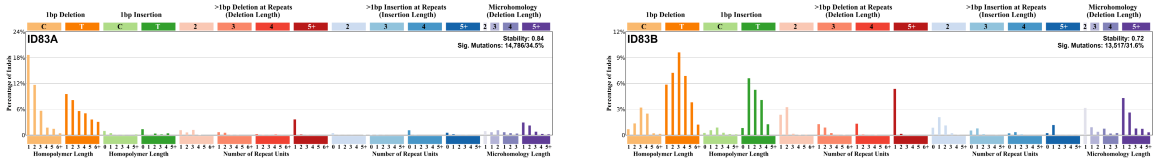

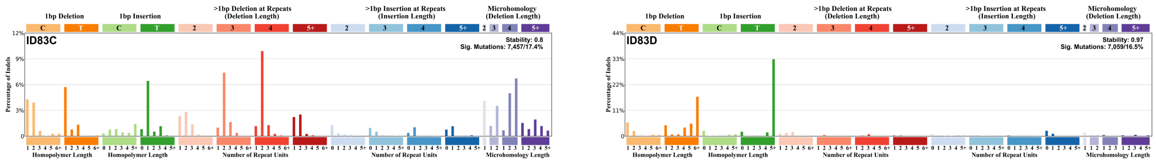


**Supplementary Figure 13. Mutation spectra of five *de novo* ID signatures from SPE.** The coloured bars at the top represent the size of the insertions or deletions. Within each panel, the x-axis depicts the count of repeat units in the reference genome corresponding to the inserted or deleted sequence.

#### 3. Mutational signatures extracted from MuSiCal

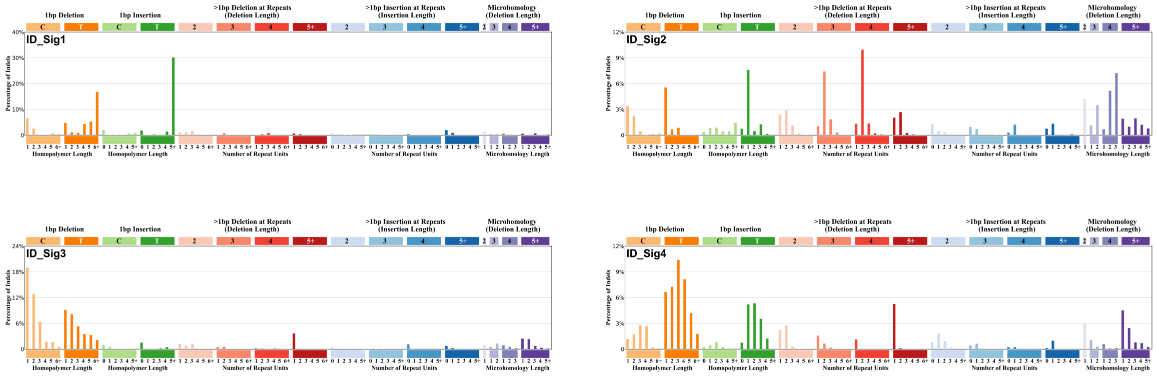


**Supplementary Figure 14. Mutation spectra of five *de novo* ID signatures from MuSiCal.** The coloured bars at the top represent the size of the insertions or deletions. Within each panel, the x-axis depicts the count of repeat units in the reference genome corresponding to the inserted or deleted sequence.

4. **Mutational signatures comparison**

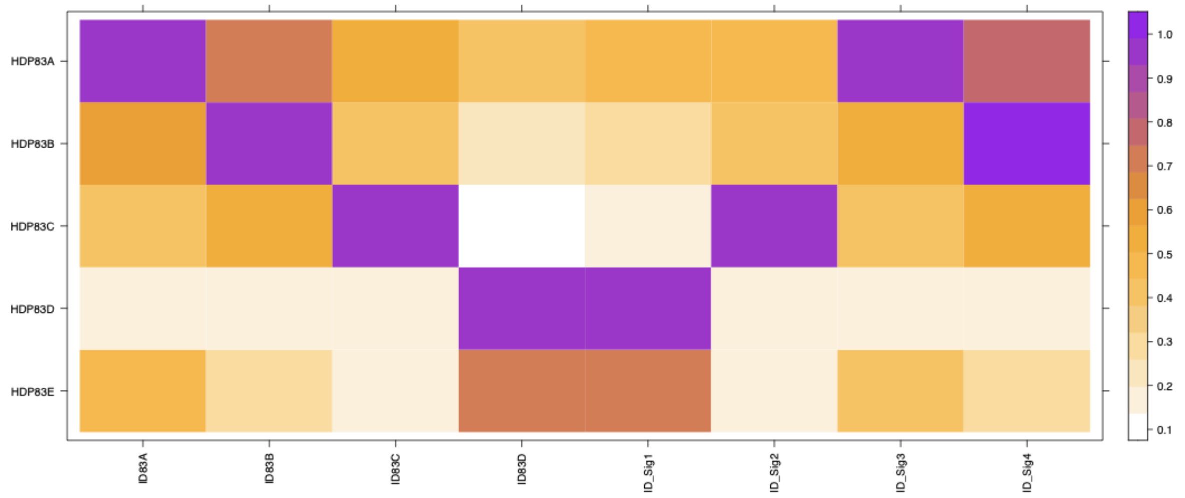


**Supplementary Figure 15. Cosine similarities between ID signatures from HDP and the other two methods.** The x-axis shows 5 *de novo* signatures extracted from HDP, the y-axis shows four *de novo* signatures from SPE, and four from MuSiCal. The colour gradient presents cosine similarity scores. The highest similarity is shown in purple, while the lowest similarity is shown in white.

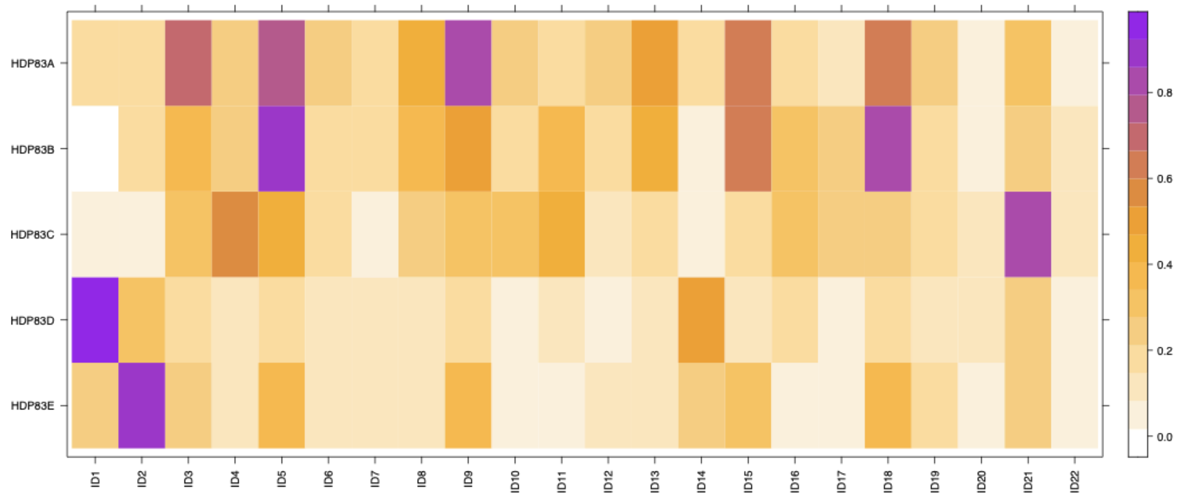


**Supplementary Figure 16. Cosine similarities between HDP83 signatures and COSMIC reference signatures.** The x-axis shows 5 *de novo* signatures extracted from HDP, and the y-axis shows a list of COSMIC reference signatures v3.4 [8]. The colour gradient presents cosine similarity scores. The highest similarity is shown in purple, while the least similar are in white.

| HDP | SPE | MuSiCal | Score | COSMIC | Concl. |
|---|---|---|---|---|---|
| HDP83A | ID83A (0.94) | ID_Sig3 (0.99) | 3 | | Y |
| HDP83B | ID83B (0.98) | ID_Sig4 (0.93) | 3 | ID5 (0.92) | Y |
| HDP83C | ID83C (0.97) | ID_Sig2 (0.98) | 3 | | Y |
| HDP83D | ID83D (0.98) | ID_Sig1 (0.97) | 3 | ID1 (0.92) | Y |
| HDP83E | | | 1 | ID2 (0.9) | Y |

**Supplementary Table D. Cross-method comparison and consensus assignment of de novo indel (ID) signatures.** For each of the five *de novo* ID signatures extracted by HDP, the best-matching *de novo* signature independently extracted by SigProfilerExtractor and MuSiCal is shown *(cosine similarity)*, together with the best-matching COSMIC v3.4 reference ID signature [8]. Score indicates the number of methods (maximum of three) in which a corresponding *de novo* signature was independently recovered. Concl. indicates whether the signature was retained (Y) in the final consensus ID signature set.

#### 5. Mutational signatures decomposition

<table>
<tr><th>HDP mutational signatures</th><th>Decomposition</th></tr>
<tr><td>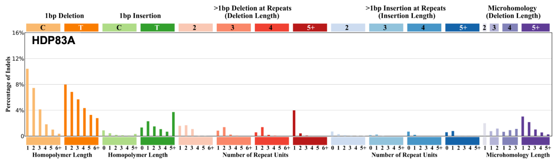
</td><td>ID3 (30.60%) & ID8 (13.14%) & ID9 (44.22%) & ID13 (12.04%)</td></tr>
<tr><td>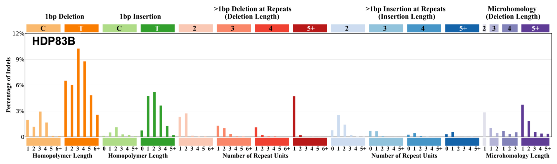
</td><td>ID5 (77.34%) & ID8 (11.20%) & ID11 (11.46%)</td></tr>
</table>

| | |
|---|---|
| 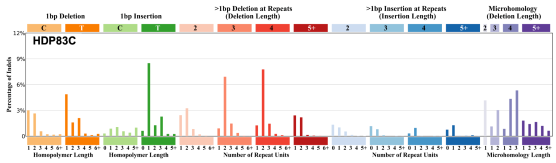 | ID9 (12.14%) & ID11 (19.82%) & ID21 (68.04%) |
| 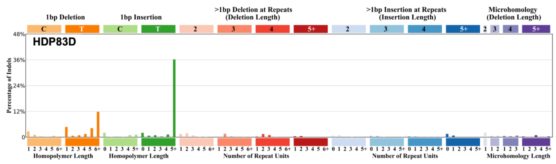 | ID1 (51.72%) & ID2 (17.56%) & ID5 (30.72%) |
| 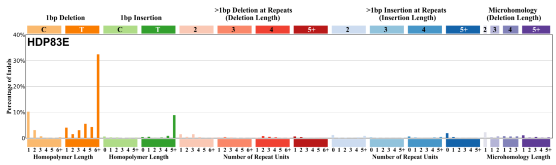 | ID1 (11.92%) & ID2 (49.22%) & ID9 (38.86%) |

**Supplementary Table E. Decomposition of consensus HDP indel signatures into COSMIC reference ID signatures.** Each HDP83-derived *de novo* ID signature was decomposed into a combination of COSMIC v3.4 reference ID signatures using SigProfilerAssignment; the relative contribution (%) of each reference signature to the reconstructed profile is shown.

#### 6. Raw mutational signatures attribution from HDP

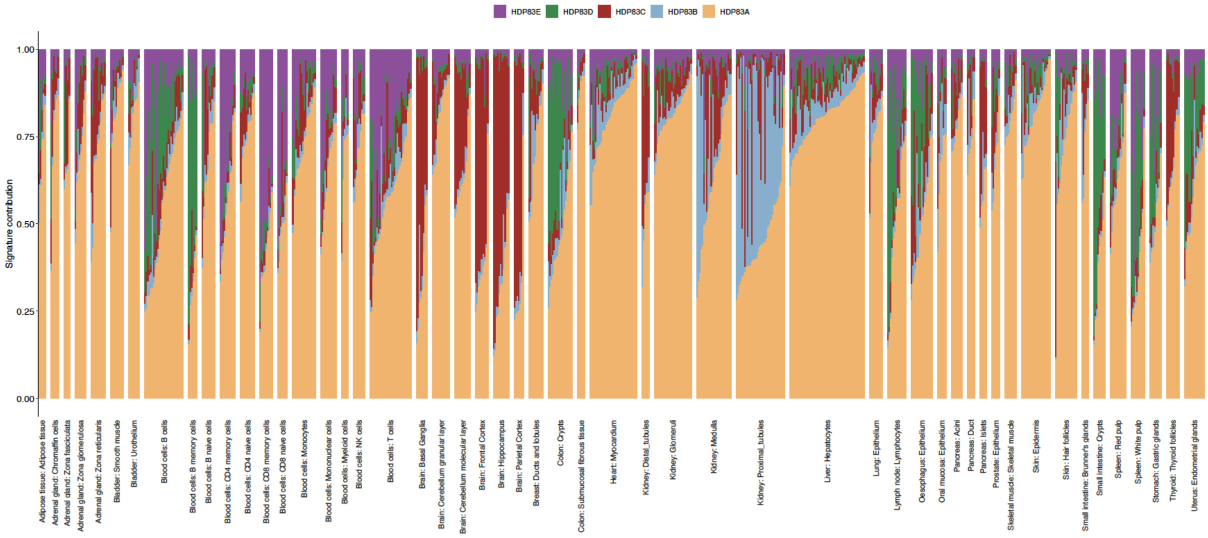


**Supplementary Figure 17. Contribution of *de novo* ID signatures to mutation catalogues of each sample.** The five *de novo* ID signatures extracted from HDP83 were attributed to each sample and grouped by tissue structure in alphabetical order. Each colour represents one signature.

### Note 4: Correlation between mutational signatures and age

For each mutational signature, we tested for accumulation of mutations within each tissue or cell type against donor age using a linear regression model. Tissue/cell types represented by fewer than five biologically independent samples were excluded from this analysis. For each regression, the fitted slope (mutations acquired per year), the two-sided p-value testing the null hypothesis that the slope is zero, and the residual standard error (RSE) of the model are reported on the corresponding panel. Shaded ribbons show the 95% confidence interval of the fitted regression line; points represent individual samples.

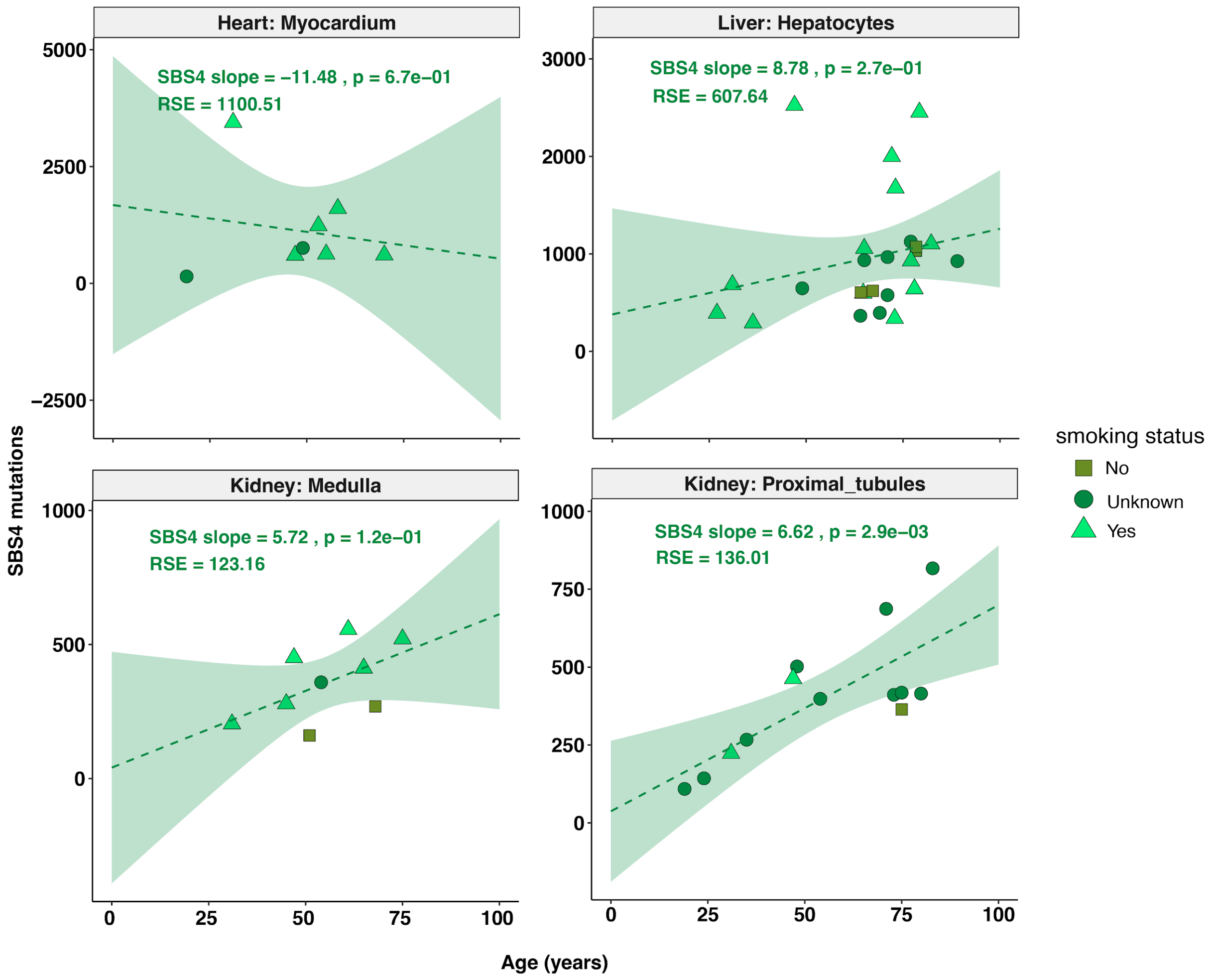


**Supplementary Figure 18. Accumulation of SBS4 mutations with age.** Linear regression of SBS4 attributed mutation burden against donor age in heart myocardium, liver hepatocytes, kidney medulla and kidney proximal tubules. Points are coloured and shaped by donor smoking status (yes, no or unknown). A significant, positive association between SBS4 burden and age was observed in kidney proximal tubules; no significant association was detected in the other tissues examined.

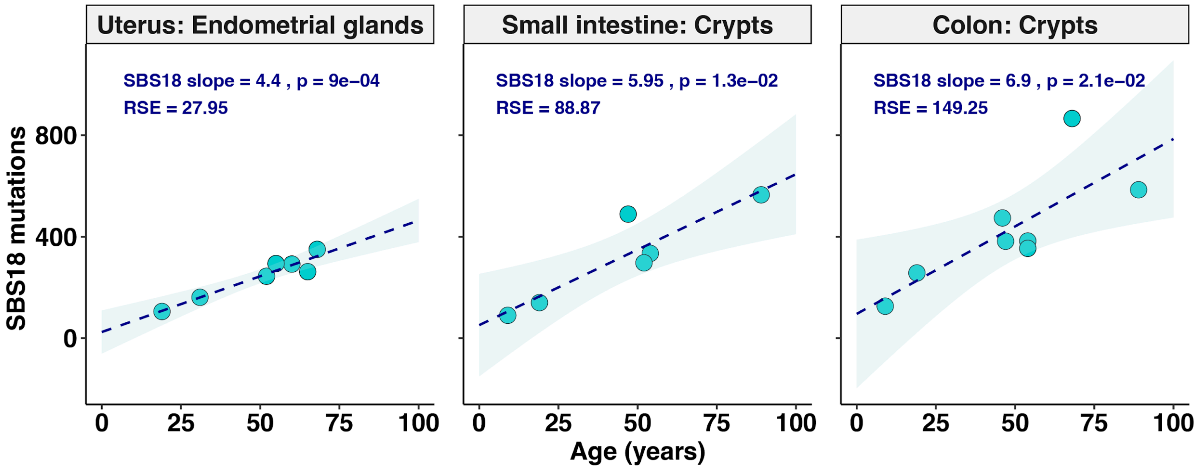


**Supplementary Figure 19. Accumulation of SBS18 mutations with age.** Linear regression of SBS18 attributed mutation burden against donor age in uterine endometrial glands, small intestinal crypts and colonic crypts. SBS18 burden increased significantly with age in all three tissues.

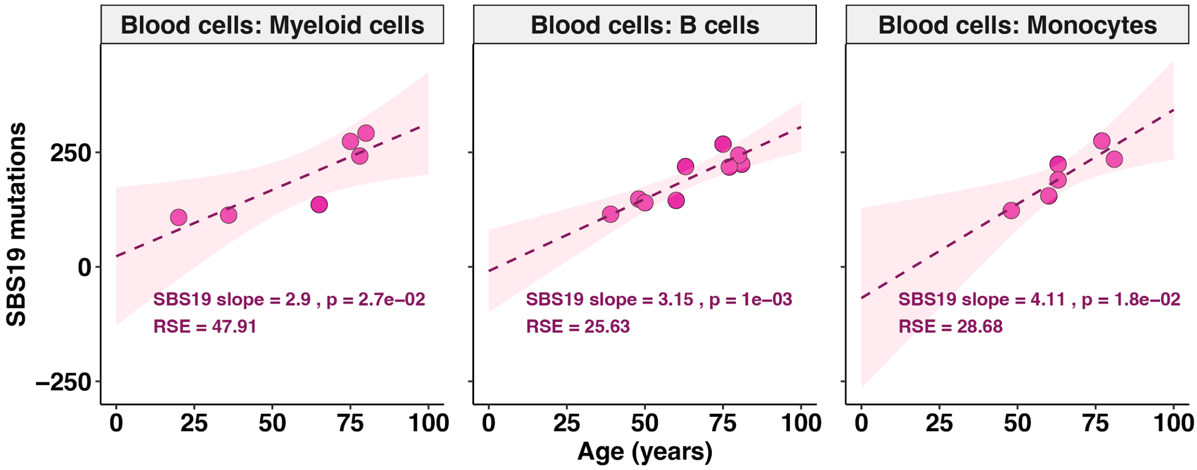


**Supplementary Figure 20. Accumulation of SBS19 mutations with age in blood cell lineages.** Linear regression of SBS19 attributed mutation burden against donor age in flow-sorted myeloid cells, B cells and monocytes. SBS19 burden increased significantly with age in all three lineages.

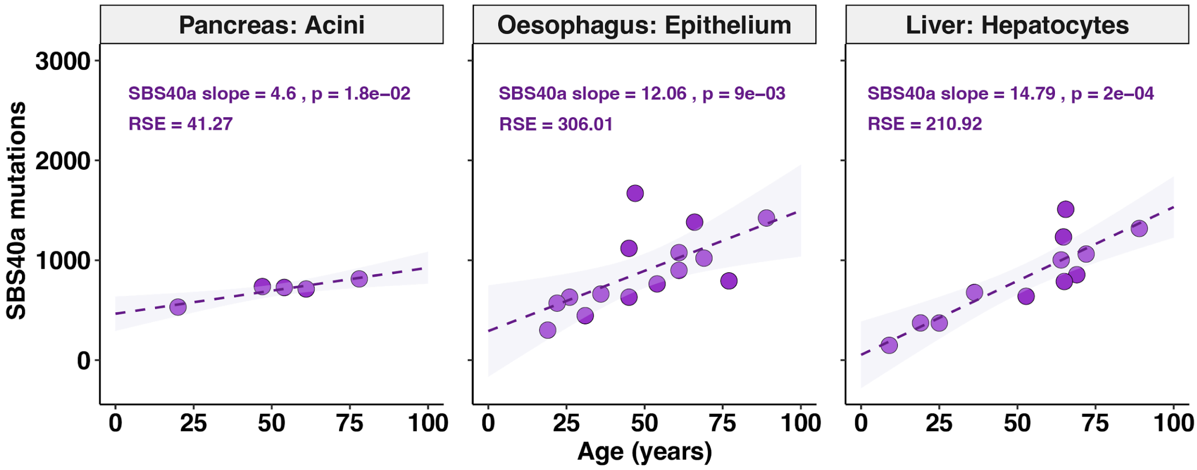


**Supplementary Figure 21. Accumulation of SBS40a mutations with age.** Linear regression of SBS40a attributed mutation burden against donor age in pancreatic acini, oesophageal epithelium and liver hepatocytes. SBS40a burden increased significantly with age in all three tissues, with the steepest rate of accumulation in liver hepatocytes.

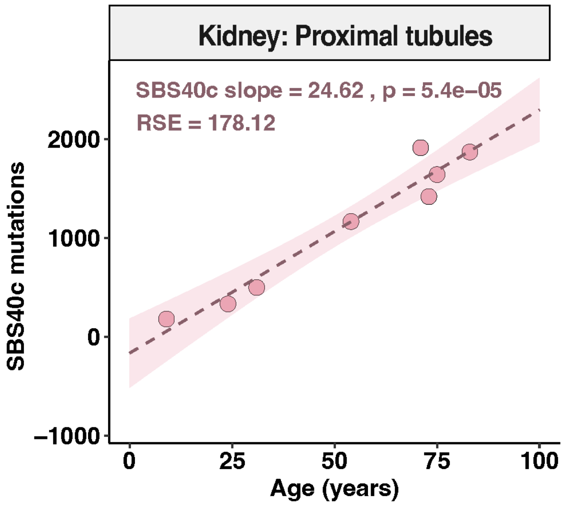


**Supplementary Figure 22. Accumulation of SBS40c mutations with age in kidney proximal tubules.** Linear regression of SBS40c attributed mutation burden against donor age. SBS40c burden increased steeply and significantly with age.

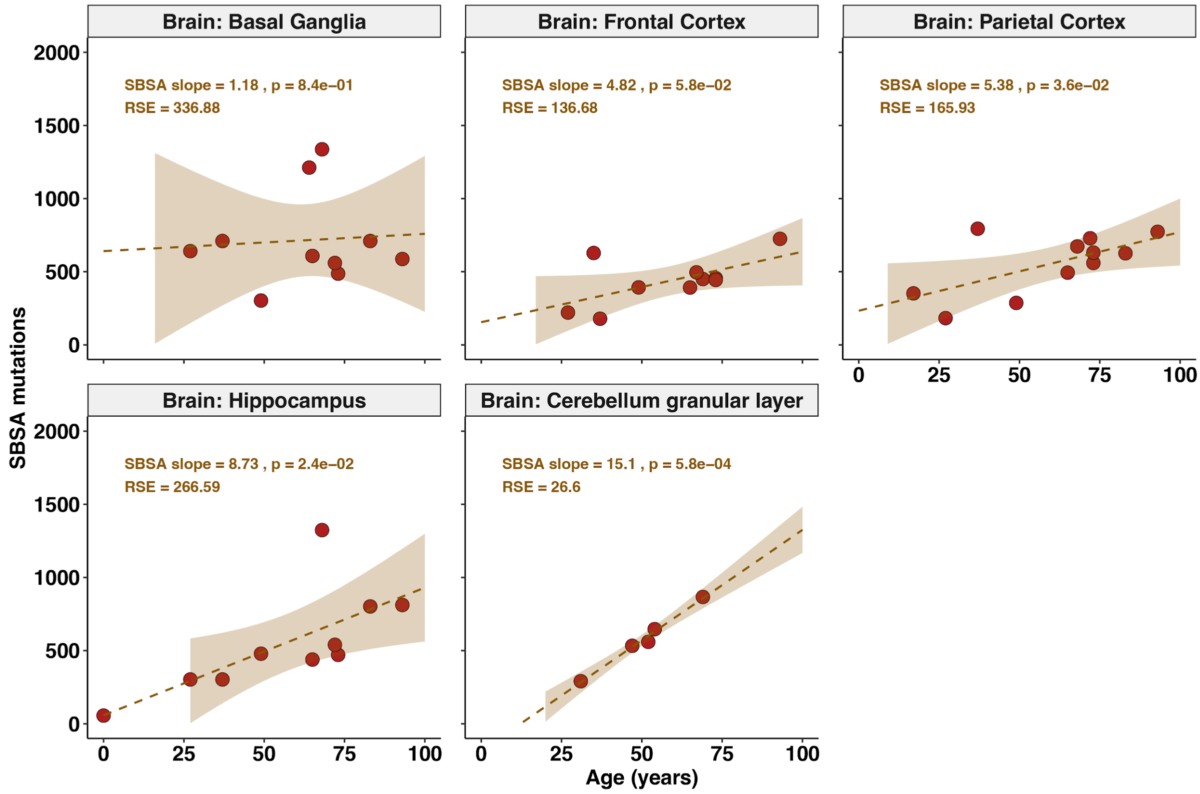


**Supplementary Figure 23. Accumulation of signature SBSA mutations with age in the brain.** Linear regression of SBSA attributed mutation burden against donor age in basal ganglia, frontal cortex, parietal cortex, hippocampus and the cerebellar granular layer. A significant, positive association with age was observed in the parietal cortex, hippocampus and cerebellar granular layer, but not in basal ganglia or frontal cortex.

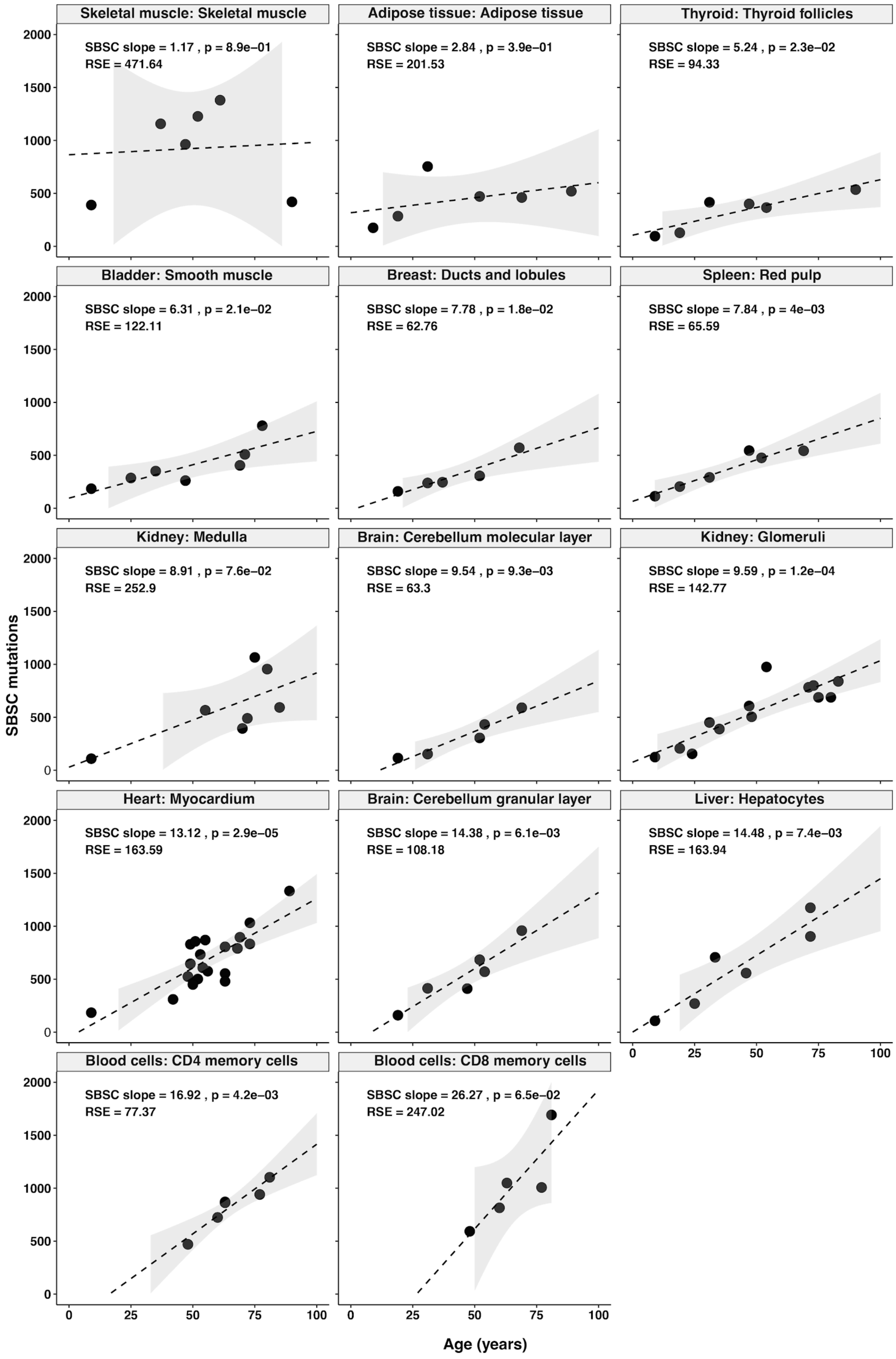
Skeletal muscle: Skeletal muscle
SBSC slope = 1.17 , p = 8.9e−01
RSE = 471.64
Adipose tissue: Adipose tissue
SBSC slope = 2.84 , p = 3.9e−01
RSE = 201.53
Thyroid: Thyroid follicles
SBSC slope = 5.24 , p = 2.3e−02
RSE = 94.33
Bladder: Smooth muscle
SBSC slope = 6.31 , p = 2.1e−02
RSE = 122.11
Breast: Ducts and lobules
SBSC slope = 7.78 , p = 1.8e−02
RSE = 62.76
Spleen: Red pulp
SBSC slope = 7.84 , p = 4e−03
RSE = 65.59
Kidney: Medulla
SBSC slope = 8.91 , p = 7.6e−02
RSE = 252.9
Brain: Cerebellum molecular layer
SBSC slope = 9.54 , p = 9.3e−03
RSE = 63.3
Kidney: Glomeruli
SBSC slope = 9.59 , p = 1.2e−04
RSE = 142.77
Heart: Myocardium
SBSC slope = 13.12 , p = 2.9e−05
RSE = 163.59
Brain: Cerebellum granular layer
SBSC slope = 14.38 , p = 6.1e−03
RSE = 108.18
Liver: Hepatocytes
SBSC slope = 14.48 , p = 7.4e−03
RSE = 163.94
Blood cells: CD4 memory cells
SBSC slope = 16.92 , p = 4.2e−03
RSE = 77.37
Blood cells: CD8 memory cells
SBSC slope = 26.27 , p = 6.5e−02
RSE = 247.02
SBSC mutations
0
500
1000
1500
2000
Age (years)
0
25
50
75
100

**Supplementary Figure 24. Accumulation of signature SBSC mutations with age across tissues.** Linear regression of SBSC attributed mutation burden against donor age in 14 tissue/cell types (skeletal muscle, adipose tissue, thyroid follicles, bladder smooth muscle, breast ducts and lobules, splenic red pulp, kidney medulla, cerebellar molecular layer, kidney glomeruli, heart myocardium, cerebellar granular layer, liver hepatocytes, and CD4+ and CD8+ memory T cells), arranged in order of increasing regression slope. SBSC burden increased significantly with age in most tissues examined, consistent with a broadly shared, clock-like mutational process.

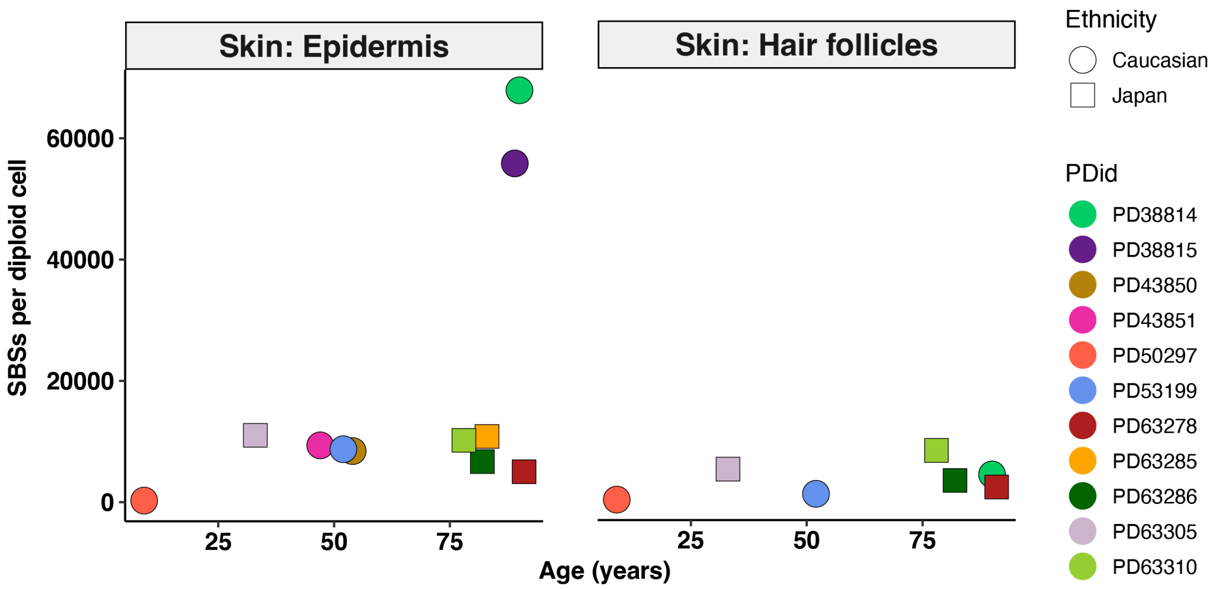


**Supplementary Figure 25. Total single-base substitution burden with age in skin.** Total number of SBS mutations per diploid genome plotted against donor age in epidermis and hair follicles. Points are coloured by donor identifier (PDid) and shaped by self-reported ethnicity (circle, Caucasian; square, Japanese). Two epidermal samples from donors PD38814 and PD38815 showed a markedly higher mutation burden than all other skin samples.

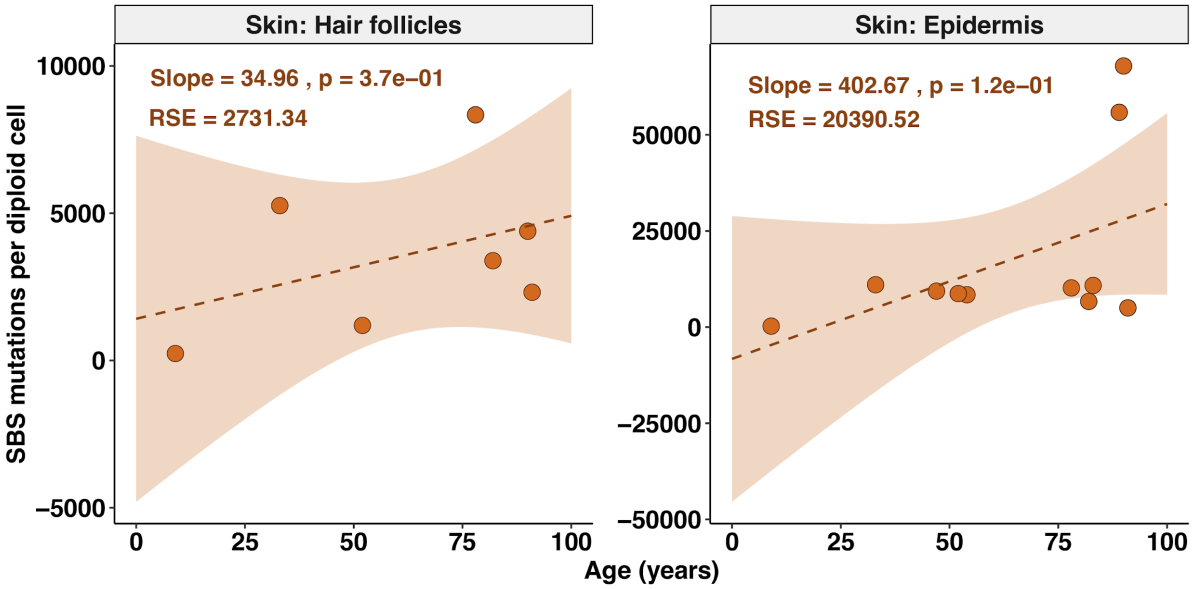


**Supplementary Figure 26. Accumulation of SBS mutations with age in the skin.** Linear regression of SBS mutation burden against donor age in hair follicles and epidermis. A negative correlation with age was observed in both cell types.

### Note 5: Sensitivity of residual standard error to donor sample size

Cell types in this study varied substantially in the number of donors contributing samples, ranging from 4 to 35 donors. Because the residual standard error (RSE) for each cell type could, in principle, be inflated or deflated simply by the amount of data available, we tested whether RSE was systematically related to the number of donors per cell type and whether this relationship was driven by any individual cell type.

We calculated Spearman's rank correlation between the number of donors and RSE across all 53 cell types (Supplementary Figure 27). This revealed no significant association (Spearman's $\rho = 0.10$, $p = 0.497$), indicating that cell types with fewer donors did not systematically show higher or lower RSE than those with more donors. Cell types with the highest RSE values, including bladder urothelium, kidney proximal tubules, and cardiac myocardium, spanned a wide range of donor numbers ($n = 8–25$), further supporting that donor number alone does not explain variation in model fit across cell types.

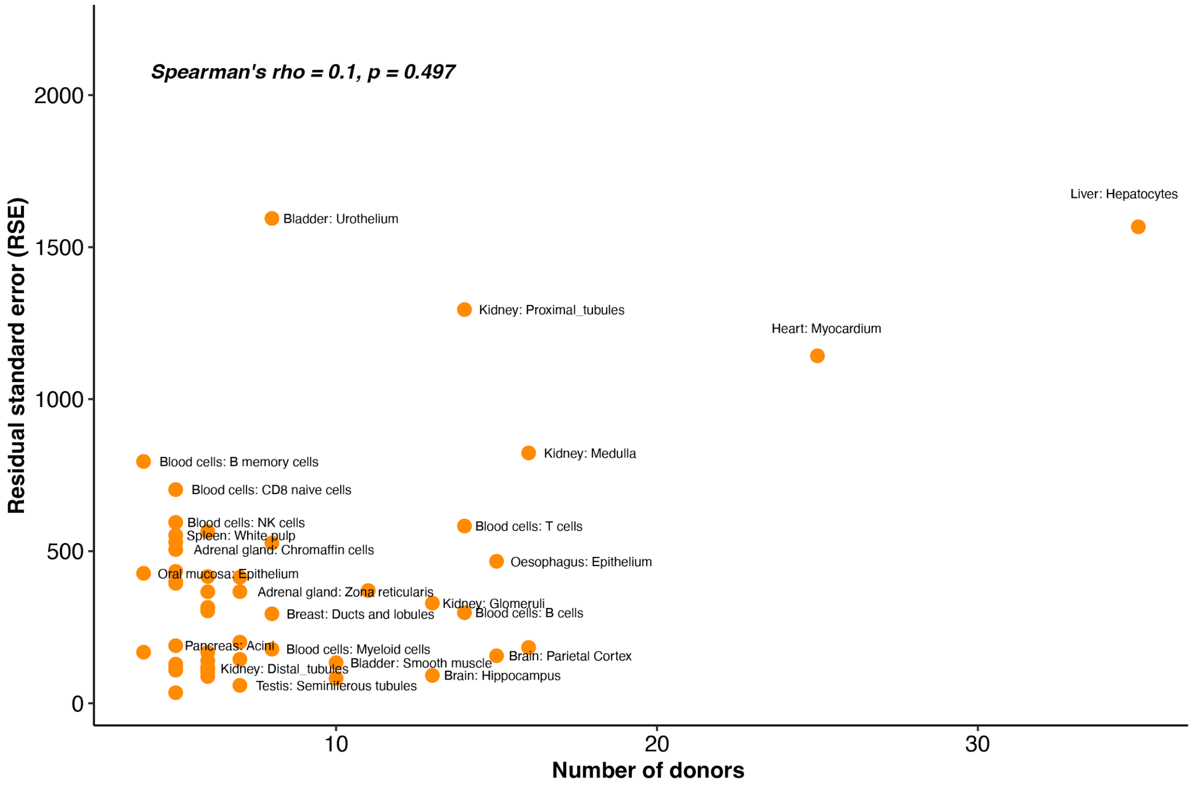

**Supplementary Figure 27. RSE versus number of donors per cell type.** Scatter plot of RSE of each cell type against the number of donors for that cell type, across 53 cell types. Each point represents one cell type, labelled by tissue and structure. Spearman's rank correlation coefficient (ρ) and associated p-value are shown.

To confirm that this null result was not driven by one or a small number of influential cell types, we performed a leave-one-out sensitivity analysis: the Spearman correlation between donor number and RSE was recalculated 53 times, each time excluding one cell type from the dataset (Supplementary Fig. 28). Across all 53 iterations, the resulting correlation remained non-significant ($p \geq 0.05$ in every case; ρ range: 0.04–0.13), with no single exclusion causing the association to cross the significance threshold.

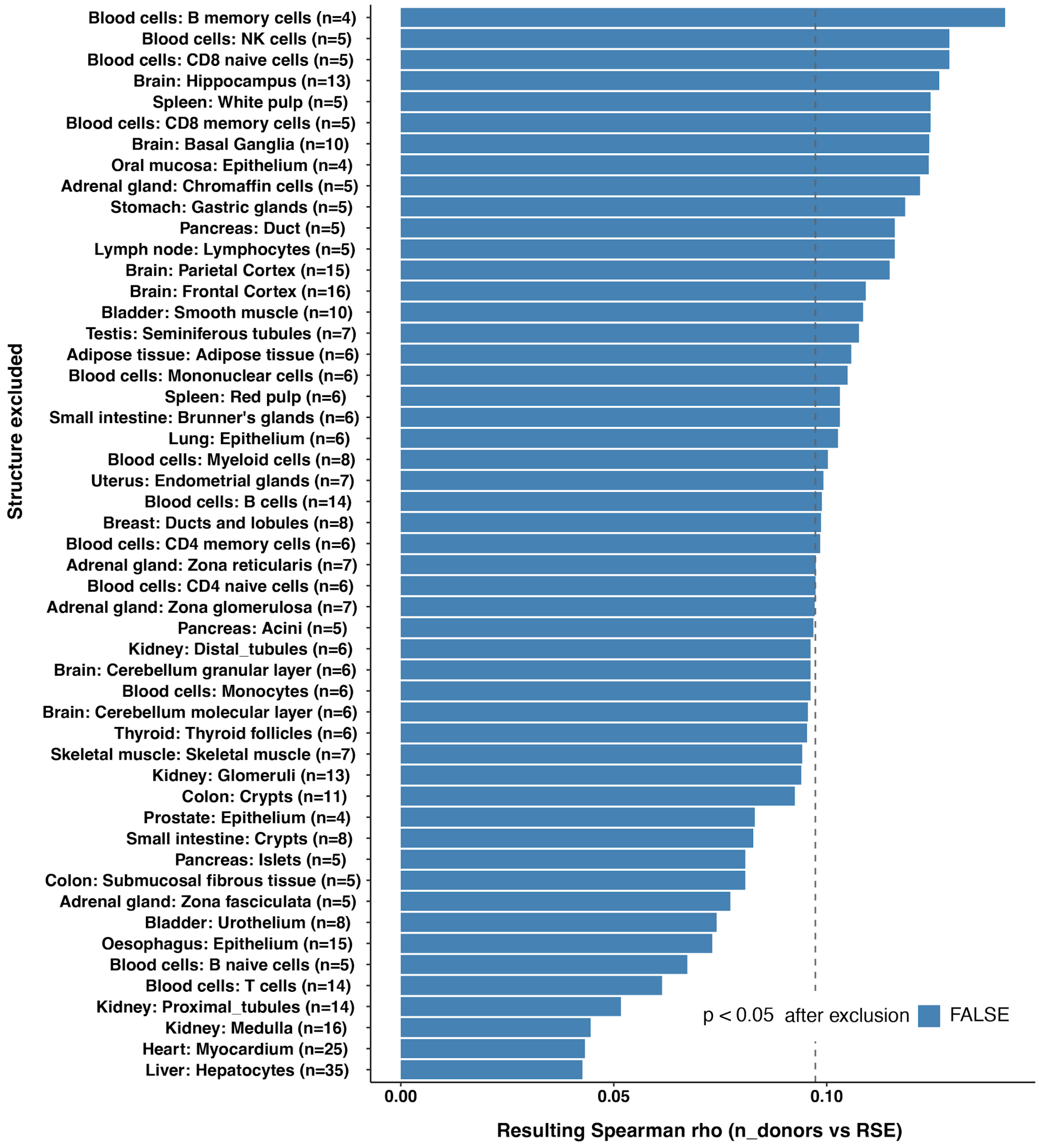


**Supplementary Figure 28. Leave-one-out sensitivity analysis confirms no association between donor number and RSE.** Bar chart showing the Spearman's rank correlation coefficient (ρ) between donor number and RSE, recalculated after excluding each of the 53 cell types in turn (one bar per exclusion, labelled with the excluded cell type and its donor number, n). The dashed vertical line marks the correlation value corresponding to $p = 0.05$. Bars are coloured by whether the correlation became significant ($p < 0.05$) after exclusion.

### Note 6: Relationship between somatic mutation burden and cancer incidence

A long-standing hypothesis in cancer biology is that tissues accumulating somatic mutations more rapidly should be at correspondingly higher risk of malignant transformation, since mutations are the substrate for oncogenic selection. We therefore tested whether the mutation rates estimated in this study, or their associated model uncertainty (RSE), correlate with population-level cancer incidence across the subset of cell types for which a corresponding cancer type exists.

Cancer incidence rates were obtained for the UK and worldwide from the International Agency for Research on Cancer (IARC) Global Cancer Observatory ( https://gco.iarc.who.int/today/). Each incidence estimate was matched to the corresponding normal cell type in our dataset by tissue of origin (n = 17 matched cell types, spanning tissues including breast, prostate, lung, colon, liver, kidney, bladder, stomach, pancreas, thyroid, uterus, oral mucosa, oesophagus, small intestine, and testis). We tested the relationship between (i) mutation rate and cancer incidence, and (ii) RSE and cancer incidence, separately for UK and worldwide incidence data, using simple linear regression (Supplementary Figure 29).

No significant association was observed between mutation rate and cancer incidence, either for UK incidence ($R^2 < 0.01$, $p = 0.750$) or worldwide incidence ($R^2 < 0.01$, $p = 0.788$). Similarly, RSE was not significantly associated with cancer incidence, for either UK incidence ($R^2 = 0.03$, $P = 0.530$) or worldwide incidence ($R^2 = 0.02$, $p = 0.629$). Cell types with among the highest mutation rates, such as liver hepatocytes and kidney proximal tubules, did not show correspondingly elevated cancer incidence, while cell types with comparatively low mutation rates, such as breast ducts and lobules and prostate epithelium, were among the tissues with the highest incidence in both datasets.

Across the cell types examined, somatic mutation rate and model uncertainty (RSE) in normal tissue did not predict population-level cancer incidence. This indicates that variation in cancer risk across tissues is not simply explained by differences in the rate of mutation accumulation in the corresponding normal cell type, consistent with the involvement of additional factors, such as differences in stem cell number and division rate, tissue-specific selection for driver mutations, and exposure to distinct mutagens, in determining cancer risk between tissues.

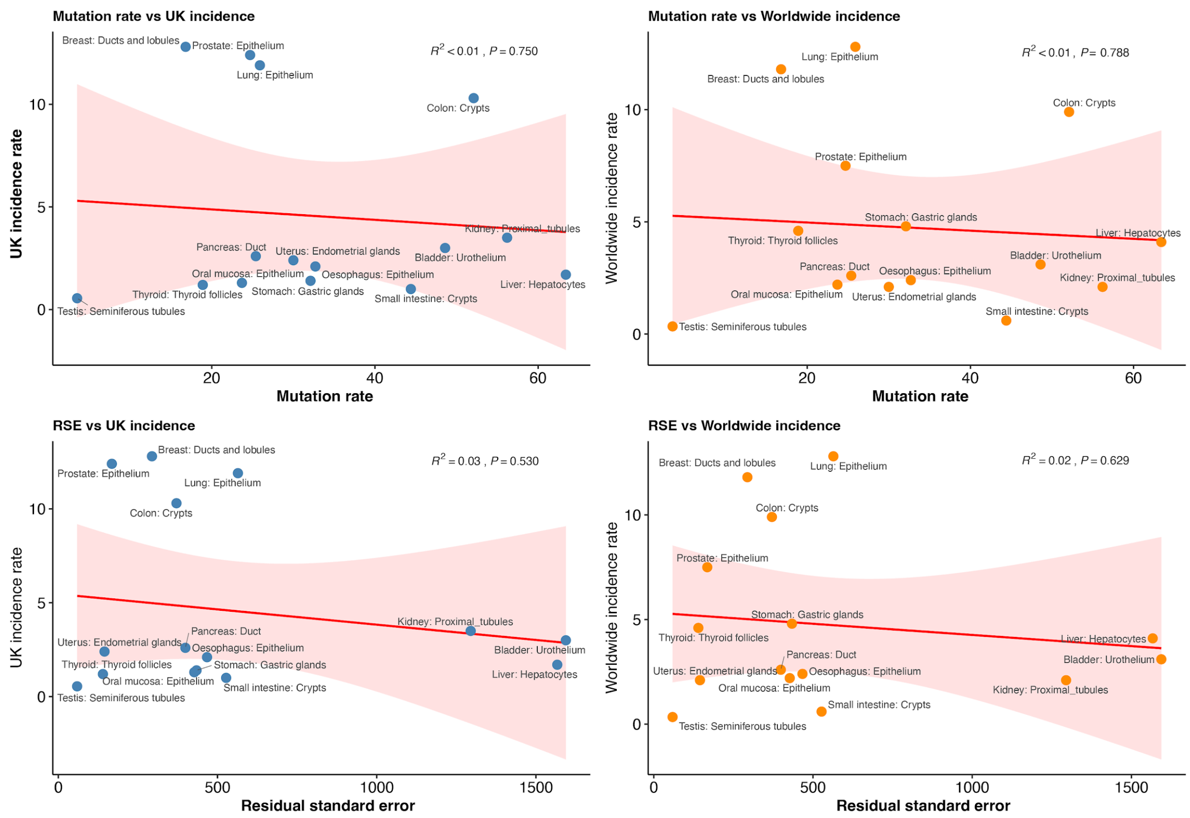


**Supplementary Figure 29. Relationship between somatic mutation burden, model uncertainty, and cancer incidence across matched cell types.** Scatter plots relating mutation rate (top row) and RSE (bottom row) from 17 cell types to their cancer incidence rates in the United Kingdom (left column) and worldwide (right column). Incidence data were obtained from the International Agency for Research on Cancer Global Cancer Observatory (GLOBOCAN). Each point represents one cell type, labelled by tissue and structure. Red line, linear regression fit; shaded ribbon, 95% confidence interval. Coefficient of determination ($R^2$) and *p-value* are shown for each panel.

### Note 7: Doublet-based substitution profiles

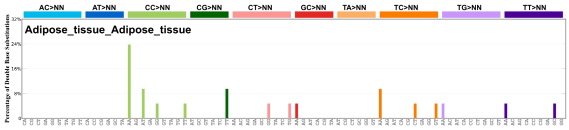


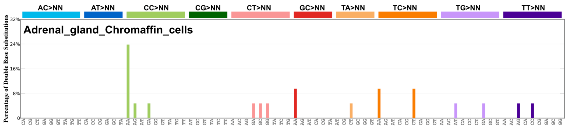


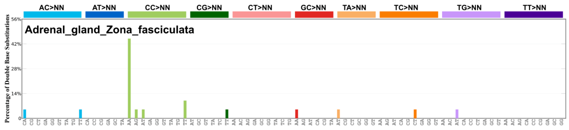


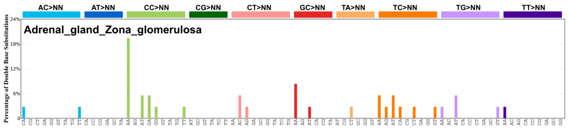


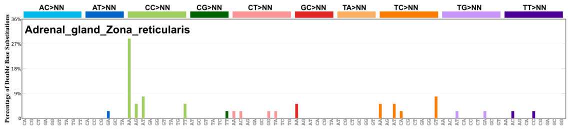


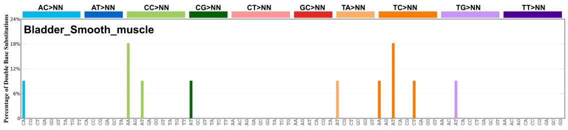


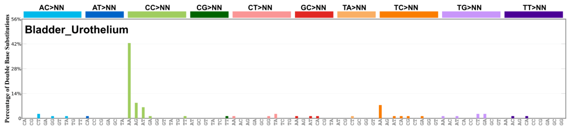


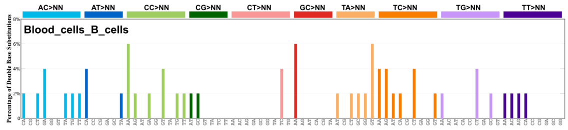


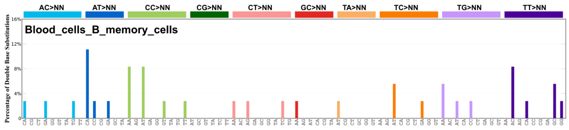


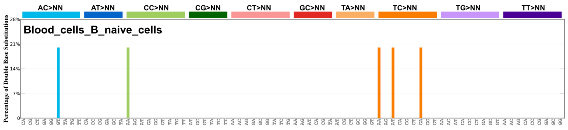


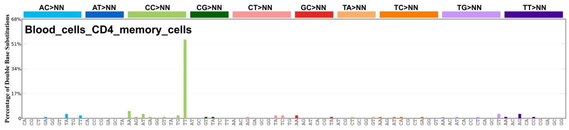


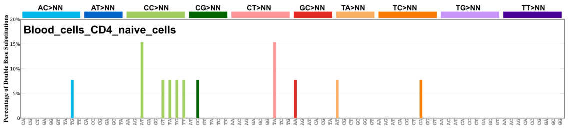


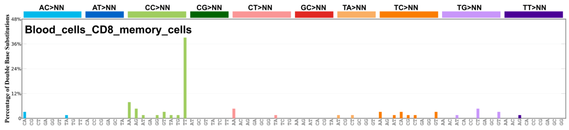


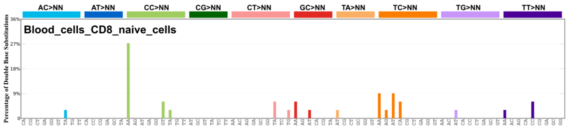

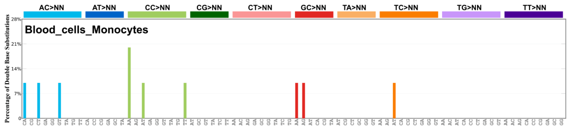
Blood_cells_Monocytes
AC>NN AT>NN CC>NN CG>NN CT>NN GC>NN TA>NN TC>NN TG>NN TT>NN
Percentage of Double Base Substitutions

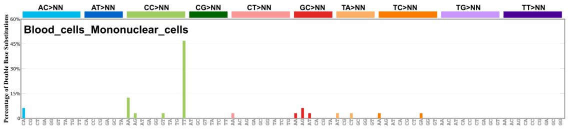
Blood_cells_Mononuclear_cells
AC>NN AT>NN CC>NN CG>NN CT>NN GC>NN TA>NN TC>NN TG>NN TT>NN
Percentage of Double Base Substitutions

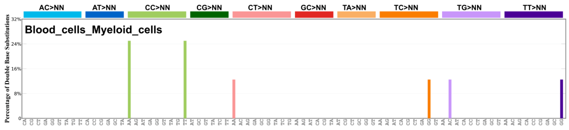
Blood_cells_Myeloid_cells
AC>NN AT>NN CC>NN CG>NN CT>NN GC>NN TA>NN TC>NN TG>NN TT>NN
Percentage of Double Base Substitutions

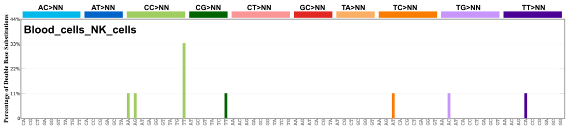
Blood_cells_NK_cells
AC>NN AT>NN CC>NN CG>NN CT>NN GC>NN TA>NN TC>NN TG>NN TT>NN
Percentage of Double Base Substitutions

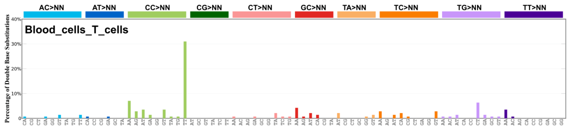
Blood_cells_T_cells
AC>NN AT>NN CC>NN CG>NN CT>NN GC>NN TA>NN TC>NN TG>NN TT>NN
Percentage of Double Base Substitutions

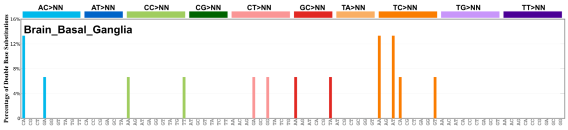
Brain_Basal_Ganglia
AC>NN AT>NN CC>NN CG>NN CT>NN GC>NN TA>NN TC>NN TG>NN TT>NN
Percentage of Double Base Substitutions

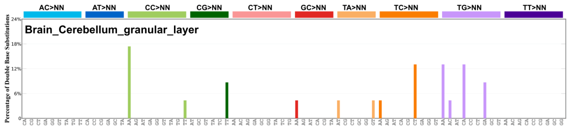
Brain_Cerebellum_granular_layer
AC>NN AT>NN CC>NN CG>NN CT>NN GC>NN TA>NN TC>NN TG>NN TT>NN
Percentage of Double Base Substitutions

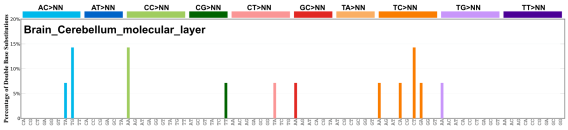
Brain_Cerebellum_molecular_layer
AC>NN AT>NN CC>NN CG>NN CT>NN GC>NN TA>NN TC>NN TG>NN TT>NN
Percentage of Double Base Substitutions

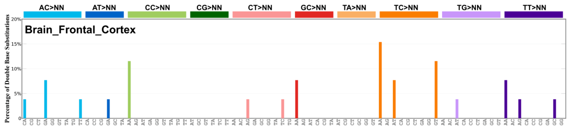
Brain_Frontal_Cortex
AC>NN AT>NN CC>NN CG>NN CT>NN GC>NN TA>NN TC>NN TG>NN TT>NN
Percentage of Double Base Substitutions

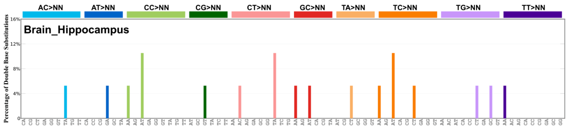
Brain_Hippocampus
AC>NN AT>NN CC>NN CG>NN CT>NN GC>NN TA>NN TC>NN TG>NN TT>NN
Percentage of Double Base Substitutions

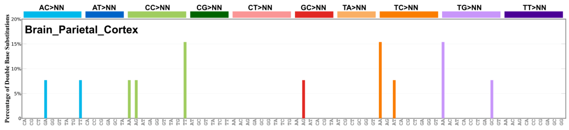
Brain_Parietal_Cortex
AC>NN AT>NN CC>NN CG>NN CT>NN GC>NN TA>NN TC>NN TG>NN TT>NN
Percentage of Double Base Substitutions

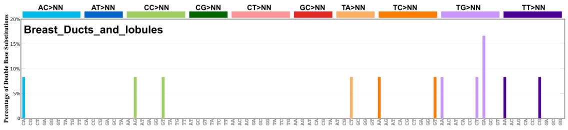
Breast_Ducts_and_lobules
AC>NN AT>NN CC>NN CG>NN CT>NN GC>NN TA>NN TC>NN TG>NN TT>NN
Percentage of Double Base Substitutions

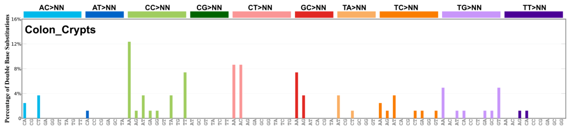
Colon_Crypts
AC>NN AT>NN CC>NN CG>NN CT>NN GC>NN TA>NN TC>NN TG>NN TT>NN
Percentage of Double Base Substitutions

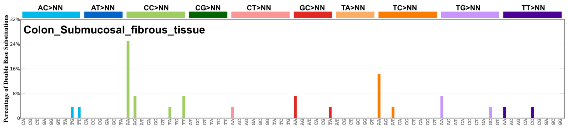
Colon_Submucosal_fibrous_tissue
AC>NN AT>NN CC>NN CG>NN CT>NN GC>NN TA>NN TC>NN TG>NN TT>NN
Percentage of Double Base Substitutions

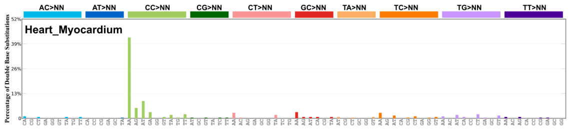
Heart_Myocardium
AC>NN AT>NN CC>NN CG>NN CT>NN GC>NN TA>NN TC>NN TG>NN TT>NN
Percentage of Double Base Substitutions

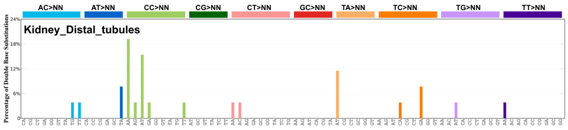
Kidney_Distal_tubules
AC>NN AT>NN CC>NN CG>NN CT>NN GC>NN TA>NN TC>NN TG>NN TT>NN
Percentage of Double Base Substitutions

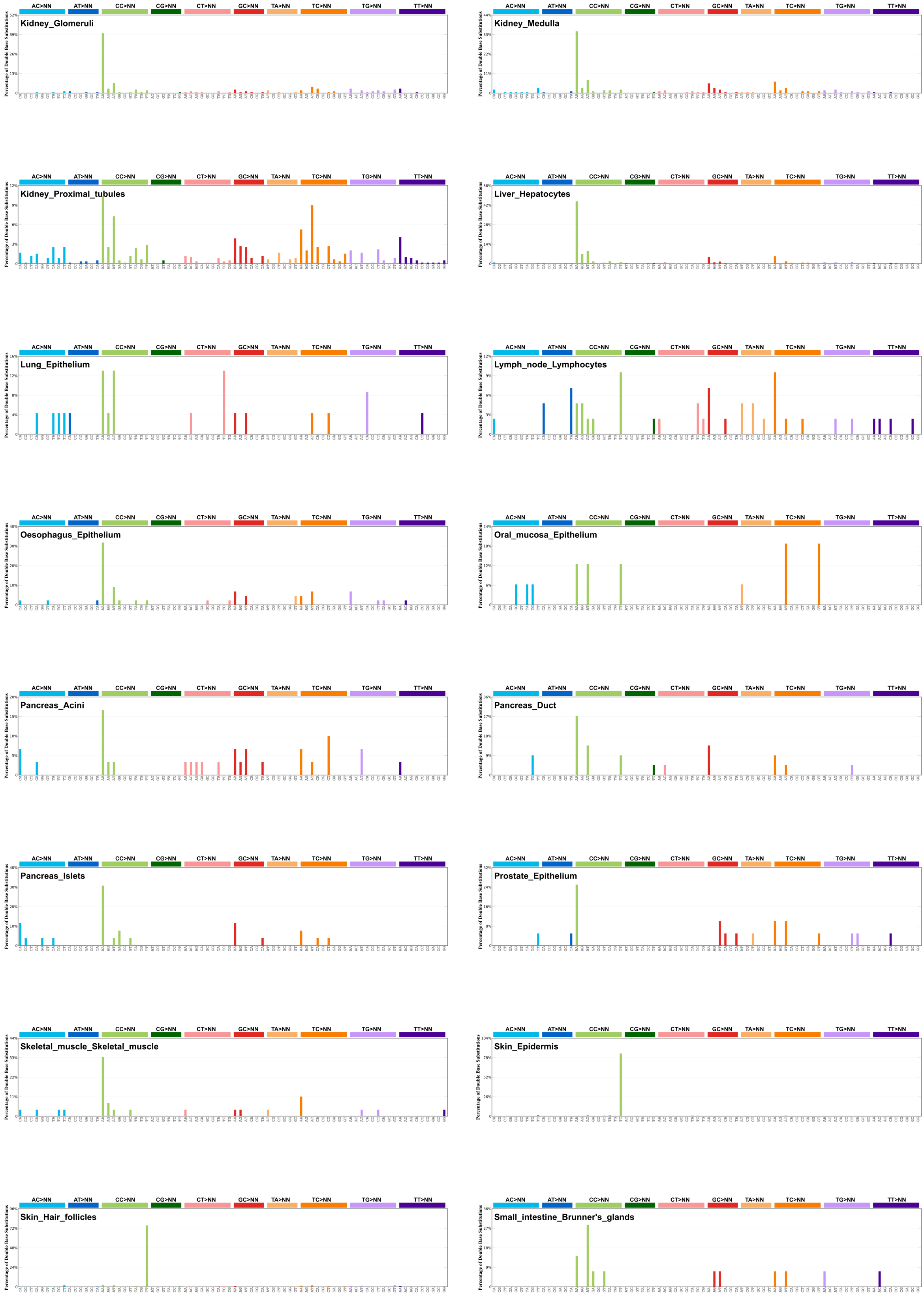
AC>NN
AT>NN
CC>NN
CG>NN
CT>NN
GC>NN
TA>NN
TC>NN
TG>NN
TT>NN
Percentage of Double Base Substitutions
Kidney_Glomeruli
Kidney_Medulla
Kidney_Proximal_tubules
Liver_Hepatocytes
Lung_Epithelium
Lymph_node_Lymphocytes
Oesophagus_Epithelium
Oral_mucosa_Epithelium
Pancreas_Acini
Pancreas_Duct
Pancreas_Islets
Prostate_Epithelium
Skeletal_muscle_Skeletal_muscle
Skin_Epidermis
Skin_Hair_follicles
Small_intestine_Brunner's_glands

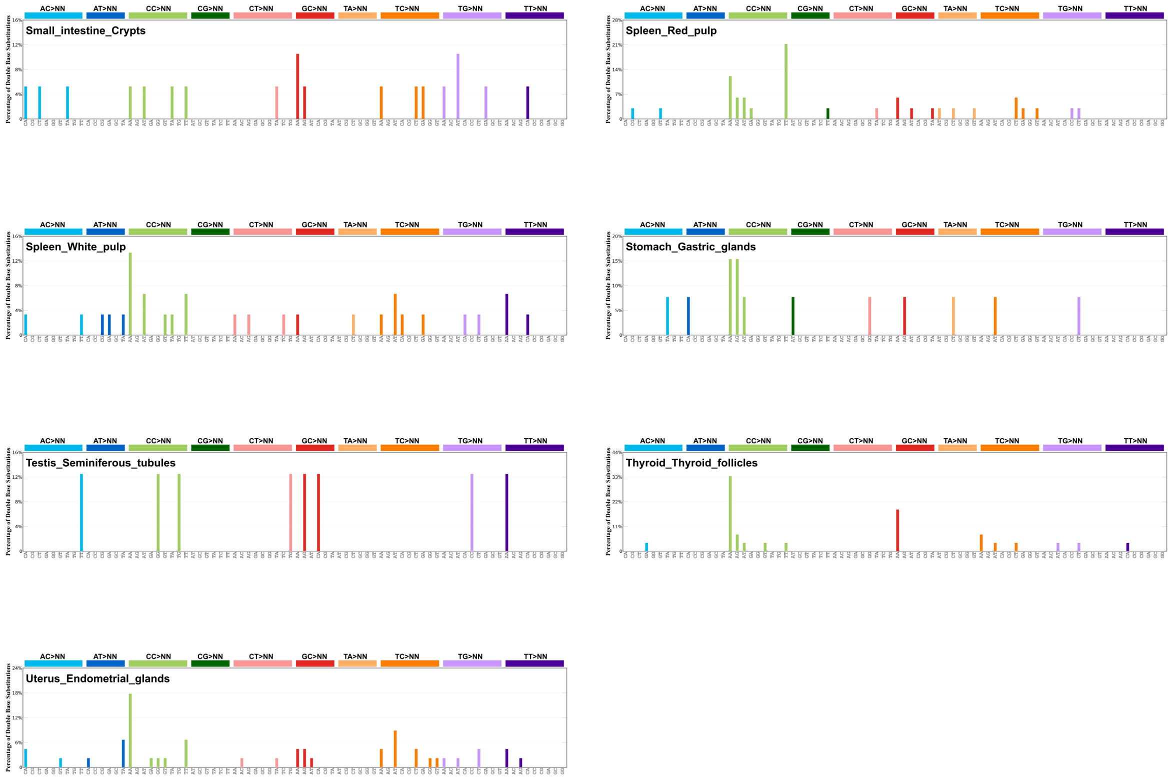


**Supplementary Figure 30. DBS profiles across 53 cell types.**

1. Bates, D., Mächler, M., Bolker, B. & Walker, S. Fitting Linear Mixed-Effects Models using lme4. *arXiv [stat.CO]* (2014) doi:10.48550/arXiv.1406.5823.
2. Linear and Nonlinear Mixed Effects Models [R package nlme version 3.1-170]. *Comprehensive R Archive Network (CRAN)* https://CRAN.R-project.org/package=nlme (2026).
3. *R: A Language and Environment for Statistical Computing. R Foundation for Statistical Computing.* (Vienna, Austria).
4. *R: A Language and Environment for Statistical Computing. R Foundation for Statistical Computing.* (Vienna, Austria).
5. Roberts, N. D. Patterns of somatic genome rearrangement in human cancer. Preprint at https://doi.org/10.17863/CAM.22674 (2018).
6. Díaz-Gay, M. *et al.* Assigning mutational signatures to individual samples and individual somatic mutations with SigProfilerAssignment. *Bioinformatics* **39**, (2023).
7. Jin, H. *et al.* Accurate and sensitive mutational signature analysis with MuSiCal. *Nat. Genet.* **56**, 541–552 (2024).
8. COSMIC. Catalogue Of Somatic Mutations In Cancer. (2024).
